# Sex-specific associations of childhood adversity with frontostriatal network organization and anhedonia in young adulthood

**DOI:** 10.64898/2026.07.24.740545

**Authors:** Ryan J. Shepherd, Matthias Pierce, Nils Muhlert, Mica Komarnyckyj, Meg Sheppard, Tobias Banaschewski, Gareth J. Barker, Arun L.W. Bokde, Rüdiger Brühl, Sylvane Desrivières, Herta Flor, Penny Gowland, Antoine Grigis, Andreas Heinz, Frauke Nees, Dimitri Papadopoulos Orfanos, Luise Poustka, Michael N. Smolka, Nathalie Holz, Nilakshi Vaidya, Henrik Walter, Robert Whelan, Paul Wirsching, Gunter Schumann, IMAGEN Consortium, Rebecca Elliott

## Abstract

**Introduction:** Anhedonia is a transdiagnostic psychiatric symptom linked to increased functional connectivity between the prefrontal cortex and striatum. Here, we examined how dimensions of early adversity contribute to this profile of connectivity.

**Methods:** In a European community sample of young adults (IMAGEN), we examined cross-sectional (n=613) and longitudinal (n=332) associations of adversity dimensions with resting-state fMRI-derived connectivity. We selected 10 ROIs from anhedonia literature, defined in the functional images as 4mm-radius spheres. We then used network-based regression models to identify clusters of ROI-ROI connections associated with threat and deprivation scores, using interaction terms to examine sex and age-specific associations. We also examined associations between adversity and anhedonia, operationalized using factor analysis of six items from self-report surveys.

**Results:** At age 18-22, we identified sex-specific associations between deprivation and connectivity for a cluster of 9 ROI-ROI connections (*p-FWE*=0.038), primarily involving the nucleus accumbens. Specifically, we observed positive associations between deprivation and connectivity in males, and negative associations in females. In the longitudinal analysis, negative deprivation associations in females attenuated with age for a cluster of 14 connections (*p-FWE*=0.009). A cluster of 17 connections also had initial positive associations with threat in females that attenuated with age (*p-FWE*=0.008). No such longitudinal changes were observed in males. Higher deprivation was linked to increased later anhedonia in males but not females (*p*=0.026).

**Conclusion:** Compared to females, young adult males may be more vulnerable to developing anhedonia after experiencing deprivation in childhood. Dimensions of early adversity are linked to distinct pathways of frontostriatal development.

## Introduction

Mechanisms underlying the emergence of poor mental health in young people are difficult to study due to significant heterogeneity in psychiatric phenotypes (1). Shifting focus from disorders to transdiagnostic constructs allows for a thorough investigation of how specific symptoms might emerge from disrupted neurodevelopment (2,3). Anhedonia, the loss of the ability to experience pleasure and interest in pursuing it (4,5), is a central and treatment-resistant component of several psychiatric disorders, particularly depression (6). Identifying factors that disrupt the development of the reward system in young people helps clarify why anhedonia and its associated disorders emerge.

Anhedonia is associated with blunted activation of the ventral striatum during reward tasks and distinct patterns of functional connectivity at rest (7–9). At rest, anhedonia is linked to frontostriatal hyperconnectivity, involving the nucleus accumbens, perigenual and subgenual anterior cingulate cortex, dorsolateral prefrontal cortex, and medial prefrontal cortex (10). These anhedonia-related patterns of connectivity align with the repeated identification of altered frontostriatal functional connectivity as a neural marker of depression (11,12). Recent work also suggests the existence of stable individual differences in topography and connectivity of the frontostriatal salience network, which predict future anhedonia in depressed patients (13). However, there is a lack of research which examines why these patterns of connectivity might emerge in young people.

Experiencing adversity in childhood shapes brain development in adolescence (14) and significantly increases the risk of poor mental health in adulthood (15). There is extensive evidence that accumulating multiple adversities is linked to worse outcomes overall (16,17). Mechanisms which link specific types of adversity to specific developmental outcomes are more difficult to study, particularly as there are many possible adversity exposures. The range of possible exposures leads to statistical challenges, and weak associations between specific adversities and outcomes (18). The dimensional model of adversity and psychopathology addresses these challenges by assigning specific adversities to dimensions of *threat* or *deprivation*, based on theory-driven knowledge of shared mechanisms impacting development (19). Experiences of threat such as abuse or witnessing violence, disrupt development by causing stress, potentially leading to maladaptive emotional processing (20). In contrast, experiences of deprivation, such as neglect or institutionalization, are characterized by insufficient emotional, social or cognitive input required for typical development (21). Deprivation has been linked to deficits of reward learning (22) and striatal responsiveness (23), whereas there is conflicting evidence that early threat contributes to blunted striatal reward responsiveness in some cases (24), and increased responsiveness in others (25). Although some prior research has investigated striatal reward processing differences associated with threat and deprivation adversities, none have examined how these dimensions differentially impact resting-state striatal connectivity development.

There is also significant heterogeneity in how adversity affects individuals’ development and health outcomes. A growing body of literature suggests that experiences of adversity and its effects on mental health differ by sex and gender (26–28). Biological sex may moderate the effects of adversity, particularly as sex hormones differentially influence brain development during puberty (29,30). Gender may also influence individuals’ capacity to adapt to adversity, as it plays a role in the resilience systems (31) which children and adolescents have access to. These systems involve psychological traits (e.g. prosocial behavior, perceived control) and external factors (e.g. friendship dynamics, experience in school), which are influenced by gender (32–34). Accounting for these potential sex and gender-specific differences is important when studying the impact of adversity on brain development and mental health.

In the present study, we examined cross-sectional and longitudinal associations between childhood adversity dimensions and the functional organization of an anhedonia network, informed by prior resting state literature (10). We also examined associations between childhood adversity and later adult anhedonia. Based on prior work highlighting the reward-blunting effects of deprivation (23), we hypothesized that deprivation would be associated with increased striatal to prefrontal connectivity and greater anhedonia. Given prior evidence of sex-specific effects of abuse and neglect on mental health (35), we expected sex differences in the associations of adversity with connectivity, but had no prior hypotheses regarding specific dimensions or the direction of associations.

## Methods

### Data source

The IMAGEN study (36) is a prospective longitudinal study of 2000 young people spread across sites in the UK, Ireland, France and Germany. A detailed description of recruitment and assessment procedures has been published previously (37). Baseline data was collected when participants were approximately aged 14, with three subsequent follow-ups around ages 16, 19 and 22. The present study used data from follow-up two and three. Researchers can freely access this dataset after submitting a research proposal and obtaining approval from the IMAGEN Consortium. Scripts used to carry out data cleaning and analyses are available in the following repository: https://github.com/r-j-shepherd/ELA_frontostriatal_connectivity.git.

### Childhood adversity

We operationalized early experiences of threat and deprivation using the abuse and neglect scale total scores from the Childhood Trauma Questionnaire (CTQ) (38,39). CTQ is a retrospective self-report measure of abuse and neglect occurring before the age of 18. CTQ includes three abuse subscales (emotional, physical, and sexual) and two neglect subscales (emotional and physical). CTQ also includes a Minimization and Denial subscale (range, 0-3), where a score above zero indicates potential bias towards under-reporting adversity. Threat and deprivation scores were converted to z-scores to allow comparability of effect sizes.

### Sample selection

We excluded participants who, at follow-up two, had: (1) unavailable pre-processed resting state fMRI data, (2) incomplete CTQ data, (3) incomplete demographic data, and (4) were under age 18. To account for potential false non-reporting of adversity, we also excluded participants who scored above zero on the Minimization and Denial subscale whilst scoring below the “none” thresholds for all CTQ subscales. Prior literature suggests that high Minimization and Denial scores impact the CTQ’s ability to discriminate between healthy and psychiatric samples and should therefore be taken into account in analyses of CTQ data (40,41). We used three subsamples of the Imagen data: a cross-sectional imaging sample of participants who met the aforementioned criteria (n=613), a longitudinal imaging sample including those who met this criteria and also had available pre-processed fMRI data at follow-up three (n=332), and a sample including those who met this criteria and also had complete anhedonia data at follow-up three (n=492).

### Anhedonia

Anhedonia was operationalized using factor analysis of six Likert items (***Table S1***) from four mental health surveys (CES-D, PHQ-8, PANAS, and WHO-5) administered at follow-up three, using a sample of participants who had complete data on selected items (n=1323, 677 missing). The sample was randomly split into 70% derivation (exploratory factor analysis; EFA) and 30% test samples (confirmatory factor analysis; CFA) (42). After selecting 6 items which were theoretically aligned with anhedonia, we carried out EFA (*psych* package in R), using the Maximum Likelihood (ML) method with oblimin rotation, to screen for items with poor loadings (<0.4) to the single anhedonia factor, then CFA (*lavaan* package in R), using Diagonally Weighted Least Squares (DWLS), to evaluate the factor structure identified in EFA. In the CFA model, we specified residual covariance for the two PANAS items and the two CES-D items, as this improved model fit statistics. We then fit the CFA model to the full sample and generated anhedonia factor scores for each participant.

### Resting-state fMRI preprocessing

MRI acquisition parameters for the IMAGEN study have been published previously (37). Preprocessing of raw fMRI data was carried out using FSL v5.0.9 and Advanced Normalisation Tools v1.9.2. This included motion correction, removal of non-brain tissue, and spatial smoothing using a 4mm gaussian kernel. Denoising was carried out using ICA⍰AROMA v0.3 (43) to identify and regress out motion⍰ and artefact⍰related components, followed by polynomial detrending (up to third order). Denoised functional data were then co-registered to each participant’s high-resolution T1 image, normalized to 2mm isotropic MNI standard space, and resliced to 3mm isotropic voxels.

We created 10 region-of-interest (ROI) 4mm-radius spheres using MNI coordinates from prior literature (44–47) (***Table S3***), in 1mm MNI152 space. This included the medial and ventromedial prefrontal cortex, and bilateral ROIs for the nucleus accumbens, perigenual anterior cingulate, subgenual anterior cingulate and dorsolateral prefrontal cortex (***Figure 1***). ROIs were resampled to match the resolution of the pre-processed 3mm isotropic resting state data.

**Figure 1:**
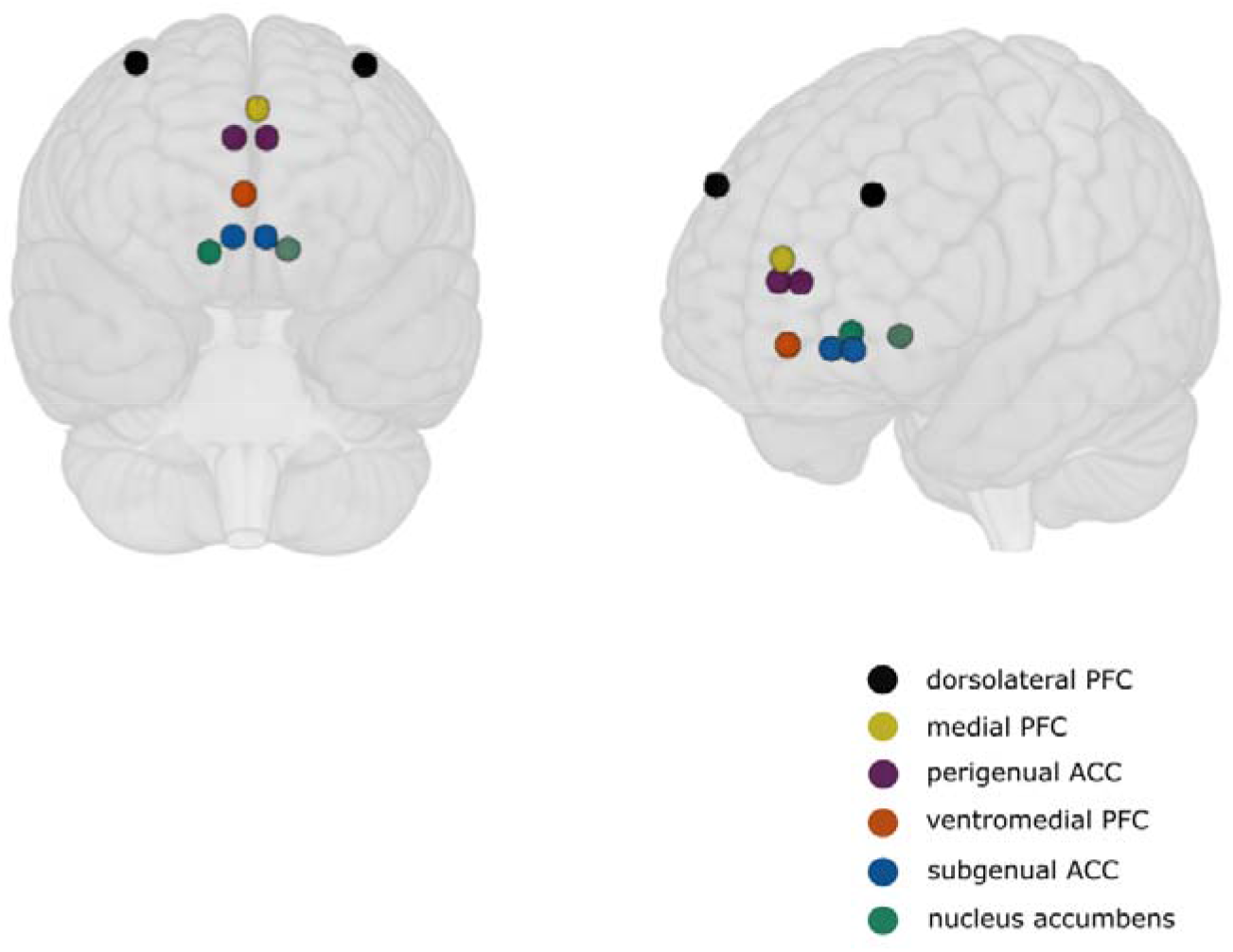
Position of 10 4mm-radius region-of-interest spheres in 1mm MNI standard space. Distinct regions are represented by colour and labelled in legend. PFC = prefrontal cortex, ACC = anterior cingulate cortex.

### Statistical analysis

Using the CONN toolbox (v22) (RRID:SCR_009550) (48) in SPM12 (49) the average BOLD timeseries was computed for each ROI sphere, and ROI-ROI connectivity matrices (i.e. Fisher-transformed correlation of timeseries between each ROI) were generated for each participant. Each connectivity matrix contained a total of 45 ROI-ROI connections, representing the combinations across all 10 ROIs.

We then used network-based regression models (*NBR* package in R) to identify clusters of ROI-ROI edges associated with threat and deprivation scores. NBR is built upon the network-based statistic theoretical framework (50). For each variable of interest, NBR identifies clusters of associated edges which are topologically linked as part of a network graph, using a significance threshold to select candidate edges and then permutation testing to generate Family-Wise Error corrected *p-values* (*p-FWE*) for each cluster.

Two *p-FWE* values were computed for each cluster: (1) a size-based *p*-value representing the probability of observing a cluster with equal or greater number of edges under the null hypothesis (*p-FWE_size_*), and (2) a strength-based *p*-value representing the probability of observing a cluster with equal or greater sum of test statistics (t-values) under the null hypothesis (*p-FWE_strength_*). We used a *p* threshold of 0.05 to identify candidate edges and then 5000 permutations to generate *p-FWE* values for each cluster.

To examine cross-sectional associations of adversity with functional connectivity at follow-up two across all 45 ROI-ROI edges, we used a simple linear NBR model, including covariates for sex, age, mean framewise displacement (to account for residual group differences in head motion (51)) and data collection site (to account for variability between MRI scanners). We used interaction terms to examine sex-specific associations of threat and deprivation with connectivity.

To examine longitudinal associations of early adversity with change in functional connectivity between follow-up two and three, we used two linear mixed effects NBR models (stratified by sex) with fixed effects for timepoint, mean framewise displacement and site; and a random effect to account for within-person clustering. We used interaction terms to examine whether threat and deprivation connectivity associations vary across age.

Sex-specific associations between adversity dimensions and anhedonia at follow-up three were examined using a linear regression model adjusted for sex, age and site; and interaction terms for sex and adversity dimensions.

Data cleaning and statistical analyses were carried out in R version 4.3.2. NBR results were visualized using BrainNet Viewer (http://www.nitrc.org/projects/bnv/) (52). To explain and visualize significant interaction clusters, we fit regression models to each cluster edge and used these models to extract beta coefficients for threat and deprivation, at each level of sex or timepoint (*emmeans* package in R). Two-sample t-tests were used to examine sex differences in threat and deprivation scores.

### Supplementary analyses

Two supplementary analyses were conducted. First, to assess the robustness of the identified clusters, we ran NBR models using different cluster-defining thresholds (p<0.05-0.005). Second, we examined sex-differences in attrition of individuals with moderate-severe abuse and neglect (classified using the CTQ severity thresholds (38)) using logistic regression. The model included interaction terms for sex and a binary moderate-severe adversity classification, with covariates for age at follow-up two, and site.

## Results

### Sample characteristics

All participants were of European descent. The mean age of participants at follow-up two was 18.48 years (SD=0.72 range=18-22) and at follow-up three was 21.99 (SD=0.63, range=20-24). Sample demographic data with mean adversity scores are included in ***Table 1***. Females had higher threat scores compared to males at follow-up two (t=2.9, p=0.004) and follow-up three (t=2.45, p=0.015). Males had higher deprivation scores compared to females at follow-up two (t=-2.23, p=0.03) but not follow-up three (p=0.19). Severe adversity experiences were uncommon; 58 participants (9.5%) met criteria for the “moderate to severe” CTQ threshold for any subscale at follow-up two, and 32 (9.6%) at follow-up three. We found marginal evidence that males who met these criteria were more likely to drop out than females (p=0.07) (***Table S13***).

**Table 1:** Table of sample characteristics, including sex, mean age (years) and IMAGEN data collection site. The range of threat scores was 15-75, and for deprivation, 10-50.

|  | **Follow-up**<br>**2** |  |  | **Follow-up**<br>**3** |  |  |
| --- | --- | --- | --- | --- | --- | --- |
|  | *n* | Mean adversity scores<br>(SD) |  | *n* | Mean adversity scores<br>(SD) |  |
|  |  | *Threat* | *Deprivation* |  | *Threat* | *Deprivation* |
| **Total** | 613 | 18.86<br>(5.14) | 15.51<br>(4.55) | 332 | 18.64<br>(5.33) | 15.55<br>(4.58) |
| **Sex** |  |  |  |  |  |  |
| Male | 296 | 18.25<br>(3.86) | 15.94<br>(4.55) | 155 | 17.90<br>(4.15) | 15.90<br>(4.72) |
| Female | 317 | 19.44<br>(6.05) | 15.12<br>(4.52) | 177 | 19.28<br>(6.12) | 15.24<br>(4.46) |
| **Site** |  |  |  |  |  |  |
| Berlin | 57 | 18.84<br>(4.70) | 15.95<br>(4.98) | 39 | 18.26<br>(3.51) | 15.79<br>(4.05) |
| Dresden | 111 | 17.91<br>(3.26) | 15.68<br>(4.68) | 54 | 17.70<br>(3.47) | 15.93<br>(5.13) |
| Dublin | 42 | 19.02<br>(4.34) | 15.19<br>(5.37) | 20 | 19.85<br>(5.54) | 16.55<br>(7.12) |
| Hamburg | 55 | 20.20<br>(6.29) | 15.60<br>(5.02) | 31 | 20.26<br>(7.42) | 15.84<br>(5.85) |
| London | 115 | 19.59<br>(6.00) | 15.01<br>(4.06) | 62 | 18.82<br>(5.15) | 14.61<br>(3.66) |
| Mannheim | 75 | 18.65<br>(4.89) | 15.47<br>(4.77) | 47 | 18.36<br>(3.99) | 14.98<br>(2.97) |
| Nottingham | 90 | 18.91<br>(6.41) | 15.77<br>(4.49) | 53 | 19.11<br>(7.78) | 15.58<br>(4.64) |
| *Paris* | 68 | 18.19<br>(3.96) | 15.56<br>(3.74) | 26 | 17.38<br>(3.70) | 16.42<br>(4.40) |
| **Mean age**<br>**(SD)** | 18.48<br>(0.72) |  |  | 21.99<br>(0.63) |  |  |

### Anhedonia factor analysis

An anhedonia factor was supported in the EFA, explaining 53% of the variance (eigenvalue = 3.20), and all items loaded strongly on to the factor (0.65-0.78) (see ***Table S2*** for each item loading). Internal consistency of the model was α=0.84.

The CFA model showed excellent fit χ²(7) = 2.82, p = 0.90; scaled χ²(7) = 6.84, p = 0.45, with additional indices indicating strong fit (CFI = 1, TLI = 1, RMSEA = 0, SRMR = 0.015).

### Cross-sectional sex-specific associations of childhood adversity dimensions with frontostriatal connectivity

At follow-up two, a cluster of 9 edges showed a significant interaction between deprivation scores and sex (*p-FWE_size_*=0.047 *p-FWE_strength_*=0.038), such that deprivation was associated with higher connectivity in males and lower connectivity in females (***Figure 2***). This cluster included connections between the right NAcc and left NAcc, bilateral pACC, and the right dlPFC; between the left NAcc and right dlPFC and vmPFC; and between the right dlPFC and vmPFC. This cluster was robust to stricter cluster defining thresholds (*p*<0.01, *p*<0.005) but had fewer connections at these thresholds (***Table S7-8***). There was one cluster identified for a potential interaction between threat scores and sex, but this was not significant (*p*-FWE_size_=0.61, *p*-FWE_strength_=0.59).

**Figure 2:**
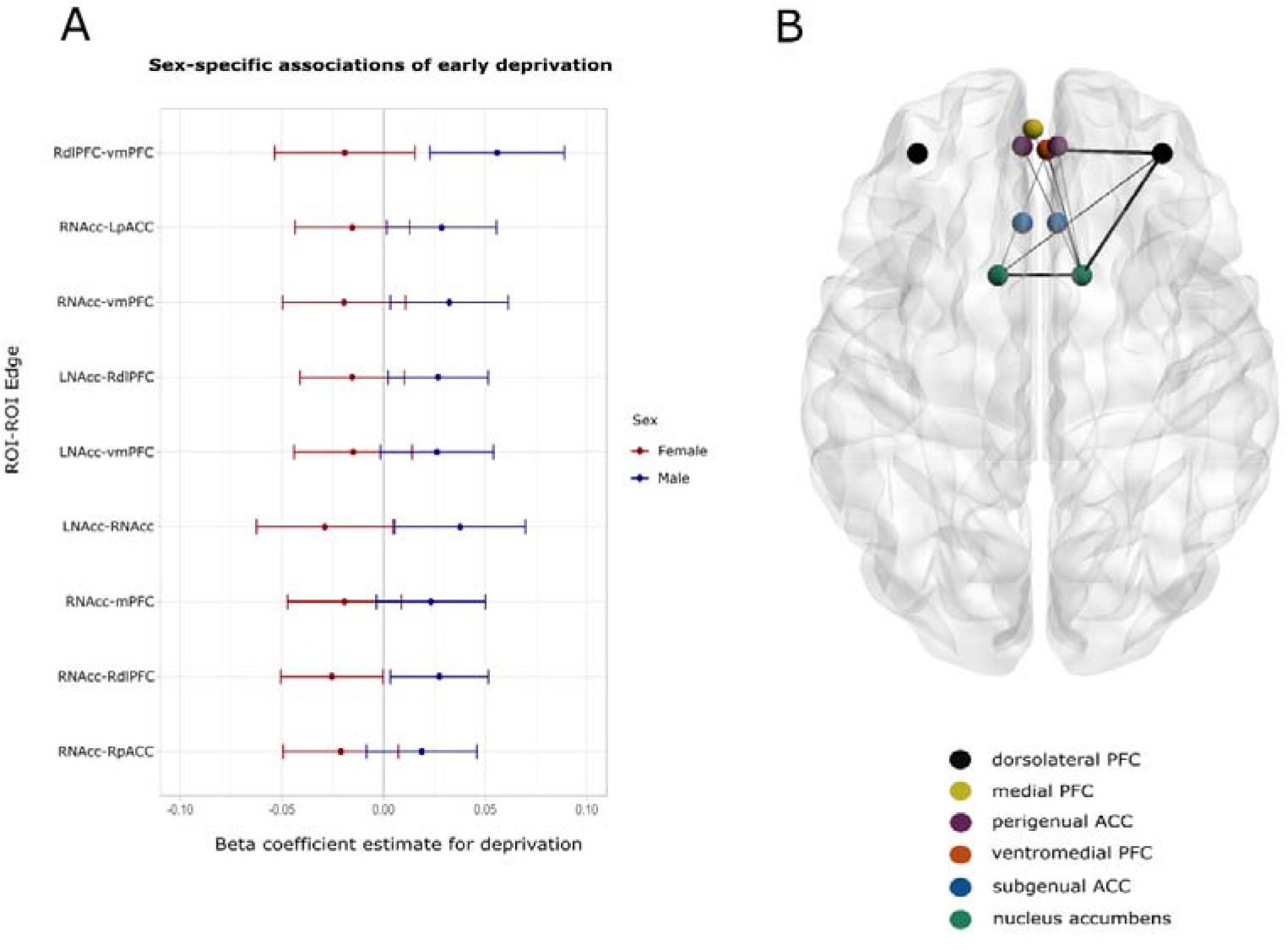
(**A**) Plot of estimated marginal slopes for deprivation score at each edge in sex x deprivation interaction cluster - extracted from linear models of connectivity for each edge in the NBR deprivation/sex interaction cluster. Error bars represent 95% confidence intervals. (**B**) 3D brain representation of a cluster of edges which showed evidence of a significant interaction between deprivation scores and sex. Distinct regions are represented by colour and labelled in legend. PFC = prefrontal cortex, ACC = anterior cingulate cortex. Line thickness corresponds to strength (i.e. the extent to which the edge exceeds the cluster defining threshold) of each edge in the cluster, with thicker lines indicating a stronger interaction effect.

There were no significant clusters for overall association with deprivation (*p-FWE_size_*=0.23, *p-FWE_strength_*=0.56), and no clusters identified for threat. There was a significant cluster of 33 edges which were associated with sex (*p-FWE_size_*<0.001, *p-FWE_strength_*<0.001), such that connectivity was increased for these edges in males compared to females. Site and mean framewise displacement were also associated with connectivity. Tables of strength statistics for edges in all clusters are included in Supplement (**Table S4**).

### Longitudinal sex stratified models of childhood adversity and change in connectivity

In the female sample mixed effects NBR model, there was a cluster of 17 edges which showed evidence of a significant interaction between threat scores and timepoint (p-FWE_size_=0.008, p-FWE_strength_=0.015), indicating initial positive associations between threat and connectivity which declined with increasing age (***Figure 3***). This cluster included connections between the left NAcc and bilateral pACC, bilateral sgACC, and right dlPFC; between the right NAcc and bilateral pACC, left sgACC, mPFC, and vmPFC; between the right pACC and right dlPFC; between the mPFC and bilateral sgACC, right dlPFC and vmPFC; and between the right dlPFC and vmPFC. This cluster was also robust to stricter cluster-defining thresholds (p<0.01, p<0.005), with fewer edges included (**Table S11-12)**.

**Figure 3:**
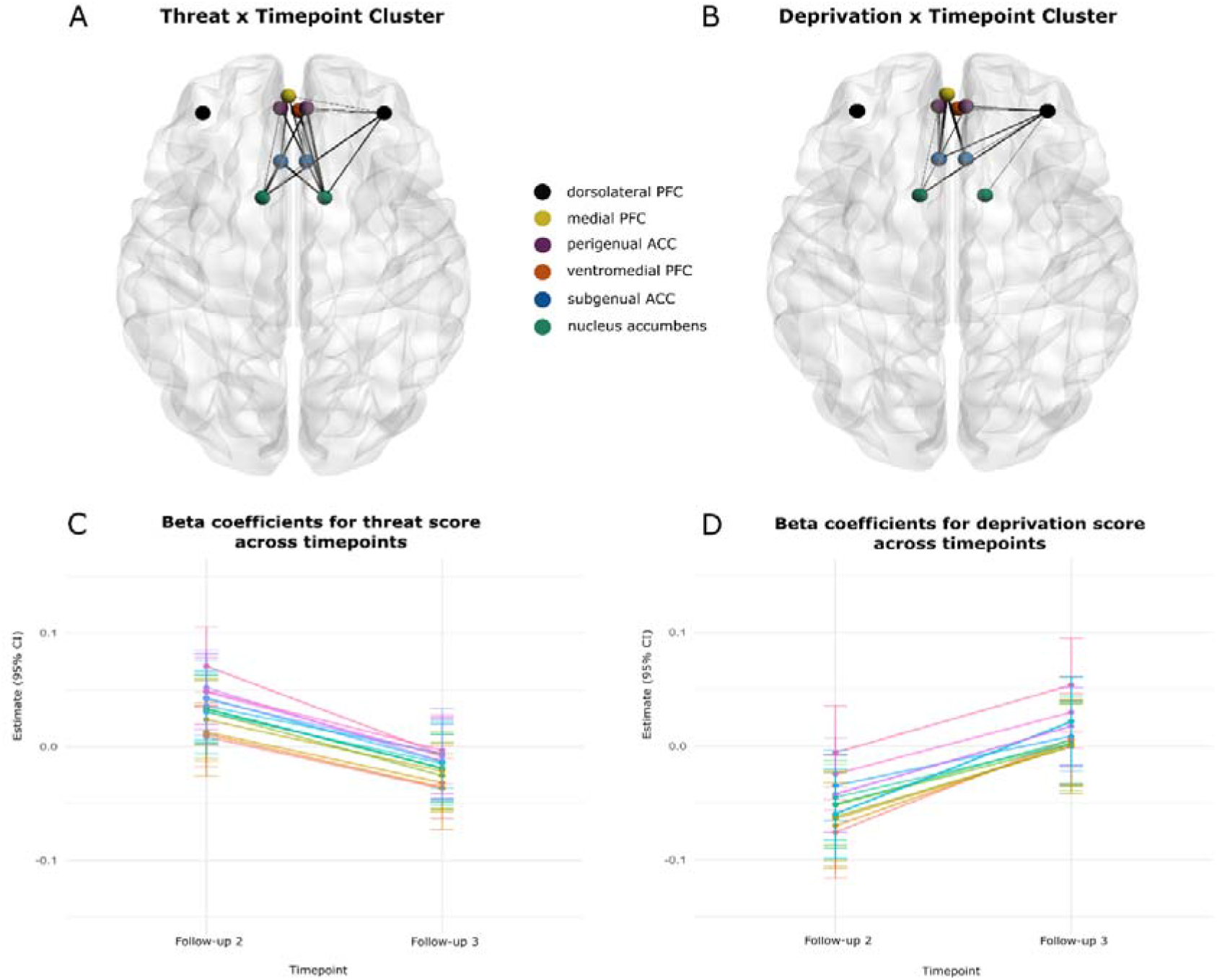
(**A-B**) 3D brain representations of clusters of edges which showed evidence of a significant interaction between (**A**) threat score and timepoint, and (**B**) deprivation score and timepoint. Distinct regions are represented by colour and labelled in legend. PFC = prefrontal cortex, ACC = anterior cingulate cortex. Line thickness corresponds to strength (i.e. the extent to which the edge exceeds the cluster defining threshold) of each edge in the cluster, with thicker lines indicating a stronger interaction effect. (**C-D**) Plots of estimated marginal slopes for (**C**) threat and (**D**) deprivation score at each timepoint for each edge in the adversity dimension x timepoint interaction clusters - extracted from linear mixed models of connectivity for each edge in the respective interaction cluster. Error bars represent 95% confidence intervals.

There was also a cluster of 14 edges which showed evidence of an interaction between deprivation scores and timepoint (p-FWE_size_=0.009, p-FWE_strength_=0.012), suggesting a negative association between deprivation and connectivity which became more positive with increasing age (***Figure 3***). This cluster included connections between the left NAcc and bilateral pACC, left sgACC and right dlPFC; between the right NAcc and right dlPFC; between the left sgACC and bilateral pACC, right dlPFC and mPFC; between right sgACC and right pACC, right dlPFC, and mPFC; and between right dlPFC and right pACC, and vmPFC. At stricter cluster-defining thresholds (p<0.01, p<0.005), this cluster was not significant.

The female sample model also had a significant cluster for overall association with threat (n_edges_=18, p-FWE_size_=0.019, p-FWE_strength_=0.053) and deprivation (n_edges_=22, p-FWE_size_=0.002, p-FWE_strength_=0.004). There were also significant clusters for site and mean framewise displacement. Edge strength statistics for each cluster are included in ***Table S6***.

In the male sample, there were no significant clusters for the deprivation/timepoint interaction term (p-FWE_size_=0.57, p-FWE_strength_=0.36), deprivation (p-FWE_size_=0.57, p-FWE_strength_=0.40), or threat (p-FWE_size_=0.42, p-FWE_strength_=0.21), and none identified for threat/timepoint. There was a significant cluster for timepoint (n_edges_=17, p-FWE_size_=0.007, p-FWE_strength_<0.001). There were also significant clusters for site and mean framewise displacement. Edge t-statistics for each cluster are included in ***Table S5***.

### Sex-specific associations between childhood adversity dimensions and anhedonia at follow-up three

We found evidence of a significant interaction between childhood deprivation scores and sex at follow-up three (p=0.028), such that there was a significant positive association between deprivation and anhedonia in males (β=0.209 95%CI [0.114, 0.305], *p*<0.001), and no clear association in females (p=0.27). Some Imagen sites were also associated with lower anhedonia scores (***Table S14***).

## Discussion

We report the first evidence that early deprivation shapes frontostriatal connectivity in opposite directions for males and females. In a frontostriatal subnetwork involving the nucleus accumbens, perigenual ACC, dorsolateral PFC, medial PFC and ventromedial PFC, higher deprivation was linked to increased connectivity in males and decreased connectivity in females. We also found, in a female subsample, that threat and deprivation had opposing longitudinal associations with frontostriatal connectivity, and that these associations became weaker with increasing age. In support of our hypothesis, increased early deprivation was also linked to greater anhedonia. However, this association was specific to males.

Several prior human studies have examined the relationship between early deprivation and frontostriatal connectivity. In a study of adolescents, increased connectivity between the nucleus accumbens and medial PFC mediated the relationship between early institutionalization and social problems (53). In preteens, early institutionalization has been linked to more diffuse frontostriatal structural connectivity, considered to arise as a result of delayed synaptic pruning (54). We found similar patterns of increased frontostriatal connectivity, specific to males who experienced early deprivation. However, other studies report a negative relationship. There is evidence early neglect is associated with reduced structural frontostriatal connectivity in children, which may be mitigated by foster placement (55). Early material deprivation has also been linked to patterns of reduced functional connectivity within the frontoparietal and default mode networks, which remain stable with age in adolescence (56). Overall, prior literature links early deprivation to differences in structural and functional network connectivity in young people. However, the direction of these associations varies by cohort age, deprivation domain, imaging modality and network definition. We provide evidence that these associations are also modified by sex and increasing age.

In animal models of early life stress, sex-specific associations of childhood adversity with frontostriatal and frontolimbic development have been reported. A rodent model of early maternal care disruption induced sex-specific alterations in the excitatory glutamatergic transmission of cortical neurons to the dorsomedial striatum, such that transmission was decreased in males, and increased in females. Specifically in these males, early disruption was also linked to impaired behavioral flexibility (57). This same model also induced blunted activation of the nucleus accumbens to rewards and anhedonia-like behavior, specifically in males (58). This supports the biological plausibility of a male-specific vulnerability to early deprivation in the context of striatal connectivity development and anhedonia. Sex-specific structural frontolimbic alterations have also been observed following unpredictable postnatal stress in rodents, with diverging patterns of increased connectivity in males and decreased connectivity in females (59). There is also evidence that suggests early maternal separation causes structural and functional frontolimbic connectivity alterations, which appear earlier in females compared to males (60). The sex differences identified in the present study may reflect a similar pattern of delayed neurodevelopmental response to deprivation in males.

Early deprivation is consistently linked to later anhedonia in young people (61–64). However, human evidence for a male-specific vulnerability to anhedonia following deprivation is limited. Some studies report no sex differences in this relationship (62), others suggest a female-specific vulnerability (61,64). These inconsistencies may be driven by sex differences in presentation across subtypes of anhedonia (social, consummatory, anticipatory, etc.). Anhedonia after early adversity may also manifest at different developmental stages in boys compared to girls, which is plausible given animal evidence (60).

A male-specific vulnerability to developing anhedonia following early deprivation could plausibly be driven by several neurobiological and psychosocial mechanisms. First, observed sex differences in the effects of deprivation may be explained by normative sex differences in brain development that emerge during puberty. Girls begin puberty at an earlier age compared to boys, and both experience different developmental milestones (65). It is plausible that deprivation induces similar altered connectivity trajectories across sexes, but differences in pubertal timing shift these trajectories earlier in females compared to males, resulting in opposite directions of associations observed cross-sectionally. An emerging theory posits that the relationship between early adversity exposures and adolescent anhedonia is mediated by chronic inflammation (68). This is informed by sex differences in immune function after puberty (69), and the deleterious effects of chronic inflammation on the availability of dopamine in the striatum (70). Further research is required to examine the directions of associations between pubertal development, specific immune factors and reward processing in adolescence.

Second, differences in vulnerability to early deprivation may be caused by gender differences in the context of resilience systems (31). Having access to these systems in childhood may buffer against the negative impact of adversity on development, and gender influences their accessibility. The gendered socialization of boys and girls leads to differences in friendship dynamics (71), social preferences (72), and the adoption of resilience-conferring traits such as prosocial behavior (73). Boys, in comparison to girls, are less encouraged to express their emotions when struggling (74). This contributes to differences in support-seeking behavior between genders (75), which likely influences the developmental consequences of early adversity. Considering evidence that the effects of early neglect on brain development are mitigated by compensatory support (55,76), it is possible that females are more able to compensate for early deprivation than males due to gender differences in friendship dynamics. The reduced frontostriatal connectivity in females with high deprivation in the present study may reflect such a compensatory adaptation.

Early threat was also associated with differences in frontostriatal development. In females, we found that early threat was linked to hyperconnectivity involving the anterior cingulate cortex, including its within-region connectivity and its connectivity to the nucleus accumbens and dorsolateral prefrontal cortex. This association seemed to diminish, or become negative, with increasing age. Studies examining the impact of early threat on functional frontostriatal connectivity are sparse, though there is evidence of a negative relationship, similarly for structural frontostriatal connectivity (77). In adult female survivors of childhood sexual abuse, reduced resting-state connectivity was observed in a subnetwork comprising the dorsal striatum, ventromedial PFC and ACC compared to controls. This was accompanied by reduced striatal dopamine transporter availability in participants with early abuse and current depression (78). Our findings suggest that this pattern of reduced frontostriatal connectivity is part of a threat-related developmental trajectory, rather than a static difference. The anterior cingulate cortex plays a central role in threat and safety processing (80), and its hyperconnectivity with prefrontal and striatal regions may reflect a maladapted threat monitoring system. This hyperconnectivity may precede a pattern of reduced connectivity later in life, aligning with evidence of accelerated and later decreased volume in frontolimbic brain structures linked to early adversity (81).

### Strengths and limitations

The present study has several strengths, particularly a large sample size which provided sufficient power to examine dimensions of adversity-, age- and sex-specific associations. Additionally, a longitudinal design allowed us to examine the impact of adversity on connectivity development, and to respect the temporal ordering of early adversity, functional network organization and anhedonia.

However, we acknowledge three main limitations. First, we used the Childhood Trauma Questionnaire to operationalize threat and deprivation, which does not capture the full breadth of threat and deprivation experiences. For example, experiences of witnessing domestic abuse are not recorded in this survey. Second, IMAGEN was designed to be ethnically homogenous (37), which limits the generalizability of our findings outside of White Europeans. Third, survey non-response and missed imaging visits at follow-up may introduce selection bias. Those with more severe mental health difficulties, potentially related to both early adversity and their current profile of frontostriatal connectivity, may be less likely to attend imaging visits. We also observed a trend towards higher attrition in males with moderate to severe adversity. Low prevalence of adversity exposure in the longitudinal male sample may have reduced power to detect longitudinal associations between adversity and connectivity.

### Conclusions

We provide evidence that early deprivation is linked to sex-specific pathways of functional frontostriatal network development and later anhedonia in young people. This suggests young adult males who experience early deprivation may be more vulnerable to developing anhedonia, compared to young adult females. In females, early threat and deprivation had distinct developmental associations with frontostriatal and anterior cingulate connectivity, suggesting patterns of accelerated and delayed development linked to adversity.

## Acknowledgements

RS is funded by the ESRC and BBSRC as part of the Soc-B Centre for Doctoral Training in Biosocial Research. MP is funded by a Henry Dale Fellowship from the Wellcome Trust and the Royal Society (224243/Z/21/Z).

RE, MK and MP were supported by the NIHR Manchester Biomedical Research Centre (NIHR203308). The views expressed are those of the authors and not necessarily those of the NIHR or the Department of Health and Social Care.

This work also received support from the following sources: the European Union-funded FP6 Integrated Project IMAGEN (Reinforcement-related behaviour in normal brain function and psychopathology) (LSHM-CT-2007-037286), the Horizon 2020 funded ERC Advanced Grant ‘STRATIFY’ (Brain network based stratification of reinforcement-related disorders) (695313), Horizon Europe ‘environMENTAL’, grant no: 101057429, UK Research and Innovation (UKRI) Horizon Europe funding guarantee (10041392 and 10038599), Human Brain Project (HBP SGA 2, 785907, and HBP SGA 3, 945539), the Chinese government via the Ministry of Science and Technology (MOST). The German Center for Mental Health (DZPG), the Bundesministerium für Bildung und Forschung (BMBF grants 01GS08152; 01EV0711; Forschungsnetz AERIAL 01EE1406A, 01EE1406B; Forschungsnetz IMAC-Mind 01GL1745B), the Deutsche Forschungsgemeinschaft (DFG project numbers 458317126 [COPE], 186318919 [FOR 1617], 178833530 [SFB 940], 386691645 [NE 1383/14-1], 402170461 [TRR 265], 454245598 [IRTG 2773]), the Medical Research Foundation and Medical Research Council (grants MR/R00465X/1 and MR/S020306/1), the National Institutes of Health (NIH) funded ENIGMA-grants 5U54EB020403-05, 1R56AG058854-01 and U54 EB020403 as well as NIH R01DA049238, the National Institutes of Health, Science Foundation Ireland (16/ERCD/3797). NSFC grant 82150710554. Further support was provided by grants from: the Eranet Neuron (Grant ANR-18-NEUR00002-01– ADORe); Agence Nationale de la Recherche (Grant ANR-12-SAMA-0004 -GeBra); Assistance-Publique Hôpitaux-de-Paris and INSERM (interface grant); Paris Descartes University (Grant collaborative-project-2010); Paris Sud University (Grant IDEX-2012); Fondation de l’Avenir (Grant AP-RM-17-013); Fondation de France (Grant 00081242); Fédération pour la Recherche sur le Cerveau, and Fondation pour la Recherche Médicale (Grants DPA20140629802 and ADOLIMIS DPP20151033945); the Ile-de-France Region (Action 16700103 -grant to QIM– VEAVE, n°23002745– 23002747).

## Disclosures

Dr Banaschewski served in an advisory or consultancy role for AGB Pharma, eye level, Infectopharm, Medice, Neurim Pharmaceuticals, Oberberg GmbH and Takeda. He received conference support or speaker’s fee by Janssen-Cilag, Medice and Takeda. He received royalities from Hogrefe, Kohlhammer, CIP Medien, Oxford University Press; the present work is unrelated to these relationships. Dr Barker has received honoraria from General Electric Healthcare for teaching on scanner programming courses. Dr Poustka served in an advisory or consultancy role for Roche and Viforpharm and received speaker’s fee by Shire. She received royalties from Hogrefe, Kohlhammer and Schattauer. The present work is unrelated to the above grants and relationships. The other authors report no biomedical financial interests or potential conflicts of interest.

## References

1. Feczko E, Miranda-Dominguez O, Marr M, Graham AM, Nigg JT, Fair DA (2019): The Heterogeneity problem: Approaches to identify psychiatric subtypes. Trends Cogn Sci 23: 584–601.

2. Chu BC, Temkin AB, Toffey K (2016): Transdiagnostic Mechanisms and Treatment for Children and Adolescents: An Emerging Field. In: Oxford Handbooks Editorial Board, editor. Oxford Handbook Topics in Psychology. Oxford University Press, p 0.

3. Dalgleish T, Black M, Johnston D, Bevan A (2020): Transdiagnostic Approaches to Mental Health Problems: Current Status and Future Directions. J Consult Clin Psychol 88: 179–195.

4. Husain M, Roiser JP (2018): Neuroscience of apathy and anhedonia: a transdiagnostic approach. Nat Rev Neurosci 19: 470–484.

5. Heinz A, Schmidt LG, Reischies FM (1994): Anhedonia in schizophrenic, depressed, or alcohol-dependent patients--neurobiological correlates. Pharmacopsychiatry 27 Suppl 1: 7–10.

6. Pizzagalli DA (2022): Anhedonia: Preclinical, Translational, and Clinical Integration, vol. 58. Switzerland: Springer Nature. Retrieved April 17, 2024, from https://link.springer.com/book/10.1007/978-3-031-09683-9

7. Pizzagalli DA (2022): Toward a Better Understanding of the Mechanisms and Pathophysiology of Anhedonia: Are We Ready for Translation? Am J Psychiatry 179: 458–469.

8. Gabbay V, Ely BA, Li Q, Bangaru SD, Panzer AM, Alonso CM, et al. (2013): Striatum-Based Circuitry of Adolescent Depression and Anhedonia. J Am Acad Child Adolesc Psychiatry 52: 628–41.e13.

9. Segarra N, Metastasio A, Ziauddeen H, Spencer J, Reinders NR, Dudas RB, et al. (2016): Abnormal Frontostriatal Activity During Unexpected Reward Receipt in Depression and Schizophrenia: Relationship to Anhedonia. Neuropsychopharmacology 41: 2001–2010.

10. Pizzagalli DA, Roberts AC (2022): Prefrontal cortex and depression. Neuropsychopharmacology 47: 225–246.

11. Cao Y, Ban M, Wang D, Kong L, He J, Chen S, et al. (2026): Association between anhedonia and ventral striatum-MPFC connectivity in first-episode, treatment-naïve major depressive disorder. J Affect Disord 392: 120214.

12. Morgan JK, Shaw DS, Olino TM, Musselman SC, Kurapati NT, Forbes EE (2016): History of Depression and Frontostriatal Connectivity During Reward Processing in Late Adolescent Boys. J Clin Child Adolesc Psychol 45: 59–68.

13. Lynch CJ, Elbau IG, Ng T, Ayaz A, Zhu S, Wolk D, et al. (2024): Frontostriatal salience network expansion in individuals in depression. Nature 633: 624–633.

14. Beck D, Whitmore L, MacSweeney N, Brieant A, Karl V, de Lange A-MG, et al. (2025): Dimensions of Early-Life Adversity Are Differentially Associated With Patterns of Delayed and Accelerated Brain Maturation. Biol Psychiatry 97: 64–72.

15. Baldwin JR, Wang B, Karwatowska L, Schoeler T, Tsaligopoulou A, Munafò MR, Pingault J-B (2023): Childhood maltreatment and mental health problems: A systematic review and meta-analysis of quasi-experimental studies. Am J Psychiatry 180: 117–126.

16. Evans GW, Li D, Whipple SS (2013): Cumulative risk and child development. Psychol Bull 139: 1342–1396.

17. Hashemi L, Fanslow J, Gulliver P, McIntosh T (2021): Exploring the health burden of cumulative and specific adverse childhood experiences in New Zealand: Results from a population-based study. Child Abuse Negl 122: 105372.

18. Green JG, McLaughlin KA, Berglund PA, Gruber MJ, Sampson NA, Zaslavsky AM, Kessler RC (2010): Childhood adversities and adult psychopathology in the National Comorbidity Survey Replication (NCS-R) I: Associations with first onset of DSM-IV disorders. Arch Gen Psychiatry 67: 113.

19. McLaughlin KA, Sheridan MA, Lambert HK (2014): Childhood Adversity and Neural Development: Deprivation and Threat as Distinct Dimensions of Early Experience. Neurosci Biobehav Rev 47: 578–591.

20. McLaughlin KA, Peverill M, Gold AL, Alves S, Sheridan MA (2015): Child Maltreatment and Neural Systems Underlying Emotion Regulation. J Am Acad Child Adolesc Psychiatry 54: 753–762.

21. Sheridan MA, McLaughlin KA (2014): Dimensions of Early Experience and Neural Development: Deprivation and Threat. Trends Cogn Sci 18: 580–585.

22. Sheridan MA, McLaughlin KA, Winter W, Fox N, Zeanah C, Nelson CA (2018): Early deprivation disruption of associative learning is a developmental pathway to depression and social problems. Nat Commun 9: 2216.

23. Hanson JL, Hariri AR, Williamson DE (2015): Blunted Ventral Striatum Development in Adolescence Reflects Emotional Neglect and Predicts Depressive Symptoms. Biol Psychiatry 78: 598–605.

24. Kasparek SW, Gastón-Panthaki A, Hanford LC, Lengua LJ, Sheridan MA, McLaughlin KA (2023): Does reward processing moderate or mediate the link between childhood adversity and psychopathology: A longitudinal study. Dev Psychopathol 35: 2338– 2351.

25. Hendrikse CJ, du Plessis S, Luckhoff HK, Vink M, van den Heuvel LL, Scheffler F, et al. (2022): Childhood trauma exposure and reward processing in healthy adults: A functional neuroimaging study. J Neurosci Res 100: 1452–1462.

26. Ho TC, Buthmann J, Chahal R, Miller JG, Gotlib IH (2024): Exploring Sex Differences in Trajectories of Pubertal Development and Mental Health Following Early Adversity. Psychoneuroendocrinology 161: 106944.

27. Supke M, Hahlweg K, Schulz W, Job A-K (2025): Sex-specific differences in the experience of adverse childhood experiences: transmission, protective, and risk factors from the perspectives of parents and their children-results of an 18-year German longitudinal study. Child Adolesc Psychiatry Ment Health 19: 46.

28. Hodes GE, Epperson CN (2019): Sex Differences in Vulnerability and Resilience to Stress Across the Life Span. Biol Psychiatry 86: 421–432.

29. Herting MM, Gautam P, Spielberg JM, Kan E, Dahl RE, Sowell ER (2014): The role of testosterone and estradiol in brain volume changes across adolescence: A longitudinal structural MRI study. Hum Brain Mapp 35: 5633–5645.

30. Ho TC, Colich NL, Sisk LM, Oskirko K, Jo B, Gotlib IH (2020): Sex differences in the effects of gonadal hormones on white matter microstructure development in adolescence. Dev Cogn Neurosci 42: 100773.

31. Davydov DM, Stewart R, Ritchie K, Chaudieu I (2010): Resilience and mental health. Clin Psychol Rev 30: 479–495.

32. Williams KEG, Krems JA, Ayers JD, Rankin AM (2022): Sex differences in friendship preferences. Evol Hum Behav 43: 44–52.

33. Xiao SX, Hashi EC, Korous KM, Eisenberg N (2019): Gender differences across multiple types of prosocial behavior in adolescence: A meta-analysis of the prosocial tendency measure-revised (PTM-R). J Adolesc 77: 41–58.

34. Vera Gil S (2024): The Influence of Gender on Academic Performance and Psychological Resilience, and the Relationship Between Both: Understanding the Differences Through Gender Stereotypes. Trends Psychol. 10.1007/s43076-024-00370-7

35. Prachason T, Mutlu I, Fusar-Poli L, Menne-Lothmann C, Decoster J, van Winkel R, et al. (2024): Gender differences in the associations between childhood adversity and psychopathology in the general population. Soc Psychiatry Psychiatr Epidemiol 59: 847–858.

36. Mascarell Maričić L, Walter H, Rosenthal A, Ripke S, Quinlan EB, Banaschewski T, et al. (2020): The IMAGEN study: a decade of imaging genetics in adolescents. Mol Psychiatry 25: 2648–2671.

37. Schumann G, Loth E, Banaschewski T, Barbot A, Barker G, Büchel C, et al. (2010): The IMAGEN study: reinforcement-related behaviour in normal brain function and psychopathology. Mol Psychiatry 15: 1128–1139.

38. Bernstein DP, Fink L, Handelsman L, Foote J, Lovejoy M, Wenzel K, et al. (1994): Initial reliability and validity of a new retrospective measure of child abuse and neglect. Am J Psychiatry 151: 1132–1136.

39. Banihashemi L, Peng CW, Verstynen T, Wallace ML, Lamont DN, Alkhars HM, et al. (2021): Opposing relationships of childhood threat and deprivation with stria terminalis white matter. Hum Brain Mapp 42: 2445–2460.

40. MacDonald K, Thomas ML, Sciolla AF, Schneider B, Pappas K, Bleijenberg G, et al. (2016): Minimization of Childhood Maltreatment Is Common and Consequential: Results from a Large, Multinational Sample Using the Childhood Trauma Questionnaire. PLoS ONE 11: e0146058.

41. Schmidt MR, Narayan AJ, Atzl VM, Rivera LM, Lieberman AF (2020): Childhood maltreatment on the Adverse Childhood Experiences (ACEs) scale versus the Childhood Trauma Questionnaire (CTQ) in a perinatal sample. J Aggress Maltreatment Trauma 29: 38–56.

42. Brown TA (2015): Confirmatory Factor Analysis for Applied Research, 2nd Ed. New York, NY, US: The Guilford Press, pp xvii, 462.

43. Pruim RHR, Mennes M, van Rooij D, Llera A, Buitelaar JK, Beckmann CF (2015): ICA-AROMA: A robust ICA-based strategy for removing motion artifacts from fMRI data. NeuroImage 112: 267–277.

44. Bartra O, McGuire JT, Kable JW (2013): The valuation system: A coordinate-based meta-analysis of BOLD fMRI experiments examining neural correlates of subjective value. NeuroImage 76: 412–427.

45. Dhami P, Atluri S, Lee JC, Knyahnytska Y, Croarkin PE, Blumberger DM, et al. (2020): Prefrontal Cortical Reactivity and Connectivity Markers Distinguish Youth Depression from Healthy Youth. Cereb Cortex N Y NY 30: 3884–3894.

46. Kelly AMC, Di Martino A, Uddin LQ, Shehzad Z, Gee DG, Reiss PT, et al. (2009): Development of Anterior Cingulate Functional Connectivity from Late Childhood to Early Adulthood. Cereb Cortex 19: 640–657.

47. Rupprechter S, Romaniuk L, Series P, Hirose Y, Hawkins E, Sandu A-L, et al. (2020): Blunted medial prefrontal cortico-limbic reward-related effective connectivity and depression. Brain J Neurol 143: 1946–1956.

48. Whitfield-Gabrieli S, Nieto-Castanon A (2012): Conn: a functional connectivity toolbox for correlated and anticorrelated brain networks. Brain Connect 2: 125–141.

49. Penny WD, Friston KJ, Ashburner JT, Kiebel SJ, Nichols TE (2011): Statistical Parametric Mapping: The Analysis of Functional Brain Images. Elsevier. Retrieved February 24, 2026, from https://shop.elsevier.com/books/statistical-parametric-mapping-the-analysis-of-functional-brain-images/penny/978-0-12-372560-8

50. Zalesky A, Fornito A, Bullmore ET (2010): Network-based statistic: Identifying differences in brain networks. NeuroImage 53: 1197–1207.

51. Goto M, Abe O, Miyati T, Yamasue H, Gomi T, Takeda T (2016): Head Motion and Correction Methods in Resting-state Functional MRI. Magn Reson Med Sci MRMS Off J Jpn Soc Magn Reson Med 15: 178–186.

52. Xia M, Wang J, He Y (2013): BrainNet Viewer: A Network Visualization Tool for Human Brain Connectomics. PLOS ONE 8: e68910.

53. Fareri DS, Gabard-Durnam L, Goff B, Flannery J, Gee DG, Lumian DS, et al. (2017): Altered ventral striatal-medial prefrontal cortex resting-state connectivity mediates adolescent social problems after early institutional care. Dev Psychopathol 29: 1865– 1876.

54. Behen ME, Muzik O, Saporta ASD, Wilson BJ, Pai D, Hua J, Chugani HT (2009): Abnormal fronto-striatal connectivity in children with histories of early deprivation: A diffusion tensor imaging study. Brain Imaging Behav 3: 292–297.

55. Bick J, Zhu T, Stamoulis C, Fox NA, Zeanah C, Nelson CA (2015): A Randomized Clinical Trial of Foster Care as an Intervention for Early Institutionalization: Long Term Improvements in White Matter Microstructure. JAMA Pediatr 169: 211.

56. Chahal R, Miller JG, Yuan JP, Buthmann JL, Gotlib IH (2022): An exploration of dimensions of early adversity and the development of functional brain network connectivity during adolescence: Implications for trajectories of internalizing symptoms. Dev Psychopathol 34: 557.

57. de Carvalho G, Khoja S, Haile MT, Chen LY (2023): Early life adversity impaired dorsal striatal synaptic transmission and behavioral adaptability to appropriate action selection in a sex-dependent manner. Front Synaptic Neurosci 15: 1128640.

58. Levis SC, Birnie MT, Bolton JL, Perrone CR, Montesinos JS, Baram TZ, Mahler SV (2022): Enduring disruption of reward and stress circuit activities by early-life adversity in male rats. Transl Psychiatry 12: 251.

59. White JD, Arefin TM, Pugliese A, Lee CH, Gassen J, Zhang J, Kaffman A (2020): Early life stress causes sex-specific changes in adult fronto-limbic connectivity that differentially drive learning. eLife 9: e58301.

60. Honeycutt JA, Demaestri C, Peterzell S, Silveri MM, Cai X, Kulkarni P, et al. (2020): Altered corticolimbic connectivity reveals sex-specific adolescent outcomes in a rat model of early life adversity. eLife 9: e52651.

61. Han J, Zhang L, Zhang C, Bi L, Wang L, Cai Y (2023): Adolescent’s anhedonia and association with childhood trauma among Chinese adolescents: a cross-sectional study. BMJ Open 13: e071521.

62. Cohen JR, McNeil SL, Shorey RC, Temple JR (2019): Maltreatment Subtypes, Depressed Mood, and Anhedonia: A Prospective Study with Adolescents. Psychol Trauma Theory Res Pract Policy 11: 704–712.

63. Fan J, Han Y, Xia J, Wang X, Liu Q, Liu Y, et al. (2023): Childhood Neglect rather than Abuse Is More Strongly Associated with Anhedonia across Major Depression and Obsessive-Compulsive Disorder Patients and University Students. Depress Anxiety 2023: 2429889.

64. Wang X, Lu J, Liu Q, Yu Q, Fan J, Gao F, et al. (2022): Childhood experiences of threat and deprivation predict distinct depressive symptoms: A parallel latent growth curve model. J Affect Disord 319: 244–251.

65. Vijayakumar N, Op de Macks Z, Shirtcliff EA, Pfeifer JH (2018): Puberty and the human brain: insights into adolescent development. Neurosci Biobehav Rev 92: 417–436.

66. Seib DR, Tobiansky DJ, Meitzen J, Floresco SB, Soma KK (2023): Neurosteroids and the mesocorticolimbic system. Neurosci Biobehav Rev 153: 105356.

67. Golden CEM, Martin AC, Kaur D, Mah A, Levy DH, Yamaguchi T, et al. (2025): Estrogen modulates reward prediction errors and reinforcement learning. Nat Neurosci 28: 2502–2514.

68. Gupta T, Eckstrand KL, Forbes EE (2024): Annual Research Review: Puberty and the development of anhedonia - considering childhood adversity and inflammation. J Child Psychol Psychiatry 65: 459–480.

69. Yang Y, Kozloski M (2011): Sex Differences in Age Trajectories of Physiological Dysregulation: Inflammation, Metabolic Syndrome, and Allostatic Load. J Gerontol Ser A 66A: 493–500.

70. Felger JC, Treadway MT (2017): Inflammation Effects on Motivation and Motor Activity: Role of Dopamine. Neuropsychopharmacol Off Publ Am Coll Neuropsychopharmacol 42: 216–241.

71. Dunbar RIM, Pearce E, Wlodarski R, Machin A (2024): Sex differences in close friendships and social style. Evol Hum Behav 45: 106631.

72. Williams KEG, Krems JA, Ayers JD, Rankin AM (2022): Sex differences in friendship preferences. Evol Hum Behav 43: 44–52.

73. Van der Graaff J, Carlo G, Crocetti E, Koot HM, Branje S (2018): Prosocial Behavior in Adolescence: Gender Differences in Development and Links with Empathy. J Youth Adolesc 47: 1086–1099.

74. Chaplin TM, Aldao A (2013): Gender Differences in Emotion Expression in Children: A Meta-Analytic Review. Psychol Bull 139: 735–765.

75. Reevy GM, Maslach C (2001): Use of Social Support: Gender and Personality Differences. Sex Roles 44: 437–459.

76. Sheridan MA, Mukerji CE, Wade M, Humphreys KL, Garrisi K, Goel S, et al. (2022): Early deprivation alters structural brain development from middle childhood to adolescence. Sci Adv 8: eabn4316.

77. Dennison MJ, Rosen ML, Sambrook KA, Jenness JL, Sheridan MA, McLaughlin KA (2019): Differential associations of distinct forms of childhood adversity with neurobehavioral measures of reward processing: A developmental pathway to depression. Child Dev 90: e96–e113.

78. Borchers LR, Dan R, Belleau EL, Kaiser RH, Clegg R, Goer F, et al. (2026): Dopaminergic Frontostriatal Pathways in Major Depressive Disorder and Childhood Sexual Abuse: A Multimodal Neuroimaging Investigation. Mol Psychiatry 31: 343– 351.

79. Ellis BJ, Figueredo AJ, Brumbach BH, Schlomer GL (2009): Fundamental Dimensions of Environmental Risk : The Impact of Harsh versus Unpredictable Environments on the Evolution and Development of Life History Strategies. Hum Nat Hawthorne N 20: 204–268.

80. Bliss-Moreau E, Santistevan AC, Bennett J, Moadab G, Amaral DG (2021): Anterior Cingulate Cortex Ablation Disrupts Affective Vigor and Vigilance. J Neurosci 41: 8075–8087.

81. Vannucci A, Fields A, Hansen E, Katz A, Kerwin J, Tachida A, et al. (2023): Interpersonal early adversity demonstrates dissimilarity from early socioeconomic disadvantage in the course of human brain development: A meta-analysis. Neurosci Biobehav Rev 150: 105210.

