## Supplementary Material for "Sex-specific associations of childhood adversity with frontostriatal network organization and anhedonia in young adulthood"

**Anhedonia factor analysis**

***Table S1:*** Items used for anhedonia factor analysis. Reverse coding is indicated by *.

| **Item** | **Phrasing** | **Response coding** |
| --- | --- | --- |
| CES-D, Q12* | *(during the past week) I was happy* | Scale of 1-4 |
| CES-D, Q16* | *(during the past week) I enjoyed life* | Scale of 1-4 |
| PHQ-8, Q1 | *(Over the last 2 weeks, how often have you been bothered by any of the following problems?)*    *Little interest or pleasure in doing things* | Scale of 1-4 |
| PANAS, Q1* | *(To what extent have you felt the following over the past week?)*    *Interested* | Scale of 1-5 |
| PANAS, Q3* | *(To what extent have you felt the following over the past week?)*    *Excited* | Scale of 1-5 |
| WHO-5, Q5* | *My daily life has been filled with things that interest me* | Scale of 1-6 |

***Table S2:*** EFA Results: factor loadings for each item.

| **Item** | **Factor Loading** | **Communality (H2)** | **Uniqueness (U2)** |
| --- | --- | --- | --- |
| WHO-5, Q5 | 0.76 | 0.58 | 0.42 |
| CES-D, Q16 | 0.78 | 0.61 | 0.39 |
| CES-D, Q12 | 0.77 | 0.58 | 0.41 |
| PHQ, Q1 | 0.67 | 0.45 | 0.55 |
| PANAS, Q1 | 0.74 | 0.54 | 0.46 |
| PANAS, Q3 | 0.65 | 0.42 | 0.58 |

**ROI Spheres**

**Table S3:** MNI coordinates from prior literature. ROIs where coordinates have been flipped on x-axis to get opposite hemisphere ROI are indicated by *.

| **ROI Name** | **MNI Coordinates** (x, y, z) | **Reference** |
| --- | --- | --- |
| Right nucleus accumbens | 12, 10, -6 | Bartra et al., 2013^1^ |
| Left nucleus accumbens | -12, 12, -6 | Bartra et al., 2013 |
| Right perigenual ACC | 5, 47, 11 | Kelly et al., 2008^2^ |
| Left perigenual ACC* | -5, 47, 11 | Kelly et al., 2008 |
| Right subgenual ACC | 5, 25, -10 | Kelly et al., 2008 |
| Left subgenual ACC* | -5, 25, -10 | Kelly et al., 2008 |
| Right dorsolateral PFC | 35, 45, 38 | Dhami et al., 2020^3^ |
| Left dorsolateral PFC | -35, 45, 38 | Dhami et al., 2020 |
| Medial PFC | -2, 52, 18 | Rupprechter et al., 2020^4^ |
| Ventromedial PFC | 2, 46, -8 | Bartra et al., 2013 |

**Main NBR Model Outputs**

The *strength* of each edge is calculated using the cluster defining threshold (CDT) and the t-statistic for each edge, representing the magnitude by which the edge t-statistic exceeds the cluster defining threshold.

**Table S4:** *Cross sectional model (CDT: p<0.05)*

| Variable | Cluster | No. Edges | p-FWE (Size) | p-FWE (Strength) | ROI1 | ROI2 | Strength |
| --- | --- | --- | --- | --- | --- | --- | --- |
| SexMale | 1 | 33 | 0.0006 | 0.0004 | LNacc | RNacc | 0.6177569 |
| SexMale | 1 | 33 | 0.0006 | 0.0004 | LNacc | LpACC | 1.5110822 |
| SexMale | 1 | 33 | 0.0006 | 0.0004 | RNacc | LpACC | 0.6914857 |
| SexMale | 1 | 33 | 0.0006 | 0.0004 | LNacc | RpACC | 1.3518196 |
| SexMale | 1 | 33 | 0.0006 | 0.0004 | RNacc | RpACC | 1.384468 |
| SexMale | 1 | 33 | 0.0006 | 0.0004 | LpACC | RpACC | 2.2315146 |
| SexMale | 1 | 33 | 0.0006 | 0.0004 | LNacc | LsgACC | 0.0229653 |
| SexMale | 1 | 33 | 0.0006 | 0.0004 | RNacc | LsgACC | 0.6516 |
| SexMale | 1 | 33 | 0.0006 | 0.0004 | LsgACC | RsgACC | 0.6393333 |
| SexMale | 1 | 33 | 0.0006 | 0.0004 | LNacc | LdlPFC | 2.5944114 |
| SexMale | 1 | 33 | 0.0006 | 0.0004 | RNacc | LdlPFC | 1.6696206 |
| SexMale | 1 | 33 | 0.0006 | 0.0004 | LpACC | LdlPFC | 1.6546375 |
| SexMale | 1 | 33 | 0.0006 | 0.0004 | RpACC | LdlPFC | 1.9540583 |
| SexMale | 1 | 33 | 0.0006 | 0.0004 | LNacc | RdlPFC | 2.5057765 |
| SexMale | 1 | 33 | 0.0006 | 0.0004 | RNacc | RdlPFC | 2.0002226 |
| SexMale | 1 | 33 | 0.0006 | 0.0004 | LpACC | RdlPFC | 0.7798511 |
| SexMale | 1 | 33 | 0.0006 | 0.0004 | RpACC | RdlPFC | 0.5690364 |
| SexMale | 1 | 33 | 0.0006 | 0.0004 | LsgACC | RdlPFC | 0.6116068 |
| SexMale | 1 | 33 | 0.0006 | 0.0004 | LdlPFC | RdlPFC | 0.7510648 |
| SexMale | 1 | 33 | 0.0006 | 0.0004 | LNacc | mPFC | 1.14166 |
| SexMale | 1 | 33 | 0.0006 | 0.0004 | LpACC | mPFC | 0.473684 |
| SexMale | 1 | 33 | 0.0006 | 0.0004 | RpACC | mPFC | 0.3262377 |
| SexMale | 1 | 33 | 0.0006 | 0.0004 | LdlPFC | mPFC | 0.8742898 |
| SexMale | 1 | 33 | 0.0006 | 0.0004 | RdlPFC | mPFC | 0.1375676 |
| SexMale | 1 | 33 | 0.0006 | 0.0004 | LNacc | vmPFC | 3.092153 |
| SexMale | 1 | 33 | 0.0006 | 0.0004 | RNacc | vmPFC | 1.5396539 |
| SexMale | 1 | 33 | 0.0006 | 0.0004 | LpACC | vmPFC | 0.8620343 |
| SexMale | 1 | 33 | 0.0006 | 0.0004 | RpACC | vmPFC | 0.9548626 |
| SexMale | 1 | 33 | 0.0006 | 0.0004 | LsgACC | vmPFC | 1.3710529 |
| SexMale | 1 | 33 | 0.0006 | 0.0004 | RsgACC | vmPFC | 0.9922957 |
| SexMale | 1 | 33 | 0.0006 | 0.0004 | LdlPFC | vmPFC | 0.9815679 |
| SexMale | 1 | 33 | 0.0006 | 0.0004 | RdlPFC | vmPFC | 0.3368163 |
| SexMale | 1 | 33 | 0.0006 | 0.0004 | mPFC | vmPFC | 0.2608974 |
| deprivation_z | 2 | 3 | 0.2396 | 0.5682 | LpACC | RpACC | -0.136677 |
| deprivation_z | 2 | 3 | 0.2396 | 0.5682 | RNacc | LdlPFC | -0.0264875 |
| deprivation_z | 2 | 3 | 0.2396 | 0.5682 | RNacc | RdlPFC | -0.0173202 |
| deprivation_z | 2 | 3 | 0.2396 | 0.5682 | LdlPFC | RdlPFC | -0.0068198 |
| deprivation_z | 2 | 3 | 0.2396 | 0.5682 | LsgACC | mPFC | -0.1711122 |
| age_fu2 | 1 | 7 | 0.08 | 0.1198 | LsgACC | LdlPFC | 0.0082693 |
| age_fu2 | 1 | 7 | 0.08 | 0.1198 | RsgACC | LdlPFC | 0.3756195 |
| age_fu2 | 1 | 7 | 0.08 | 0.1198 | RpACC | RdlPFC | 0.3561416 |
| age_fu2 | 1 | 7 | 0.08 | 0.1198 | RsgACC | RdlPFC | 0.2527332 |
| age_fu2 | 1 | 7 | 0.08 | 0.1198 | RdlPFC | mPFC | 0.4098852 |
| age_fu2 | 1 | 7 | 0.08 | 0.1198 | LNacc | vmPFC | -0.1262096 |
| age_fu2 | 1 | 7 | 0.08 | 0.1198 | RpACC | vmPFC | -0.3491843 |
| siteDRESDEN | 1 | 1 | 0.6158 | 0.331 | LNacc | RNacc | -0.483576 |
| siteDRESDEN | 1 | 1 | 0.6158 | 0.331 | LpACC | LsgACC | -0.3954289 |
| siteDRESDEN | 1 | 1 | 0.6158 | 0.331 | RpACC | LsgACC | -0.1817581 |
| siteDRESDEN | 1 | 1 | 0.6158 | 0.331 | LpACC | RsgACC | -0.305336 |
| siteDRESDEN | 1 | 1 | 0.6158 | 0.331 | RpACC | RsgACC | -0.1954823 |
| siteDRESDEN | 1 | 1 | 0.6158 | 0.331 | RpACC | mPFC | -0.1208703 |
| siteDRESDEN | 1 | 1 | 0.6158 | 0.331 | LsgACC | mPFC | -0.2090473 |
| siteDRESDEN | 1 | 1 | 0.6158 | 0.331 | RsgACC | mPFC | -0.323567 |
| siteDRESDEN | 1 | 1 | 0.6158 | 0.331 | LpACC | vmPFC | -1.4497925 |
| siteDRESDEN | 1 | 1 | 0.6158 | 0.331 | RpACC | vmPFC | -1.0238661 |
| siteDRESDEN | 1 | 1 | 0.6158 | 0.331 | mPFC | vmPFC | -0.205378 |
| siteDUBLIN | 1 | 1 | 0.6096 | 0.0564 | LNacc | RNacc | 3.3745765 |
| siteDUBLIN | 1 | 1 | 0.6096 | 0.0564 | LpACC | RpACC | 1.8907722 |
| siteDUBLIN | 1 | 1 | 0.6096 | 0.0564 | LpACC | LsgACC | 1.2559427 |
| siteDUBLIN | 1 | 1 | 0.6096 | 0.0564 | RpACC | LsgACC | 1.2148694 |
| siteDUBLIN | 1 | 1 | 0.6096 | 0.0564 | LpACC | RsgACC | 1.2513778 |
| siteDUBLIN | 1 | 1 | 0.6096 | 0.0564 | RpACC | RsgACC | 1.6996476 |
| siteDUBLIN | 1 | 1 | 0.6096 | 0.0564 | LsgACC | RsgACC | 0.7171838 |
| siteDUBLIN | 1 | 1 | 0.6096 | 0.0564 | LdlPFC | RdlPFC | 2.674201 |
| siteDUBLIN | 1 | 1 | 0.6096 | 0.0564 | LpACC | mPFC | 2.6352953 |
| siteDUBLIN | 1 | 1 | 0.6096 | 0.0564 | RpACC | mPFC | 1.6880787 |
| siteDUBLIN | 1 | 1 | 0.6096 | 0.0564 | LsgACC | mPFC | 1.7152074 |
| siteDUBLIN | 1 | 1 | 0.6096 | 0.0564 | RsgACC | mPFC | 1.7208996 |
| siteDUBLIN | 1 | 1 | 0.6096 | 0.0564 | LdlPFC | mPFC | 0.1155516 |
| siteDUBLIN | 1 | 1 | 0.6096 | 0.0564 | RpACC | vmPFC | 0.362831 |
| siteDUBLIN | 1 | 1 | 0.6096 | 0.0564 | LsgACC | vmPFC | 0.4708177 |
| siteDUBLIN | 1 | 1 | 0.6096 | 0.0564 | mPFC | vmPFC | 1.0275344 |
| siteHAMBURG | 3 | 12 | 0.0266 | 0.0138 | LsgACC | RsgACC | 1.0590068 |
| siteHAMBURG | 3 | 12 | 0.0266 | 0.0138 | LpACC | LdlPFC | 0.1619147 |
| siteHAMBURG | 3 | 12 | 0.0266 | 0.0138 | RpACC | LdlPFC | 0.9363091 |
| siteHAMBURG | 3 | 12 | 0.0266 | 0.0138 | LpACC | RdlPFC | 0.1957555 |
| siteHAMBURG | 3 | 12 | 0.0266 | 0.0138 | RpACC | RdlPFC | 1.1659544 |
| siteHAMBURG | 3 | 12 | 0.0266 | 0.0138 | LsgACC | RdlPFC | 0.2692128 |
| siteHAMBURG | 3 | 12 | 0.0266 | 0.0138 | LdlPFC | RdlPFC | 1.9047274 |
| siteHAMBURG | 3 | 12 | 0.0266 | 0.0138 | LsgACC | mPFC | -0.043171 |
| siteHAMBURG | 3 | 12 | 0.0266 | 0.0138 | RsgACC | mPFC | -0.0058765 |
| siteHAMBURG | 3 | 12 | 0.0266 | 0.0138 | LdlPFC | mPFC | 1.3230757 |
| siteHAMBURG | 3 | 12 | 0.0266 | 0.0138 | RdlPFC | mPFC | 0.5712416 |
| siteHAMBURG | 3 | 12 | 0.0266 | 0.0138 | LpACC | vmPFC | -0.2700713 |
| siteLONDON | 1 | 36 | 0 | 0 | LNacc | RNacc | 7.2585548 |
| siteLONDON | 1 | 36 | 0 | 0 | LNacc | LpACC | 5.8601176 |
| siteLONDON | 1 | 36 | 0 | 0 | RNacc | LpACC | 3.4715485 |
| siteLONDON | 1 | 36 | 0 | 0 | LNacc | RpACC | 5.5008753 |
| siteLONDON | 1 | 36 | 0 | 0 | RNacc | RpACC | 4.2948265 |
| siteLONDON | 1 | 36 | 0 | 0 | LpACC | RpACC | 5.3791486 |
| siteLONDON | 1 | 36 | 0 | 0 | LNacc | LsgACC | 0.9758073 |
| siteLONDON | 1 | 36 | 0 | 0 | RNacc | LsgACC | 1.5974208 |
| siteLONDON | 1 | 36 | 0 | 0 | LNacc | RsgACC | 2.5699636 |
| siteLONDON | 1 | 36 | 0 | 0 | RNacc | RsgACC | 2.640254 |
| siteLONDON | 1 | 36 | 0 | 0 | LsgACC | RsgACC | 8.2258087 |
| siteLONDON | 1 | 36 | 0 | 0 | LNacc | LdlPFC | 2.8656884 |
| siteLONDON | 1 | 36 | 0 | 0 | RNacc | LdlPFC | 0.6690324 |
| siteLONDON | 1 | 36 | 0 | 0 | LpACC | LdlPFC | 2.0792032 |
| siteLONDON | 1 | 36 | 0 | 0 | RpACC | LdlPFC | 2.107233 |
| siteLONDON | 1 | 36 | 0 | 0 | LsgACC | LdlPFC | 2.9306038 |
| siteLONDON | 1 | 36 | 0 | 0 | RsgACC | LdlPFC | 3.9899914 |
| siteLONDON | 1 | 36 | 0 | 0 | LNacc | RdlPFC | 2.9519362 |
| siteLONDON | 1 | 36 | 0 | 0 | RNacc | RdlPFC | 1.9279582 |
| siteLONDON | 1 | 36 | 0 | 0 | LpACC | RdlPFC | 1.5536306 |
| siteLONDON | 1 | 36 | 0 | 0 | RpACC | RdlPFC | 2.3283378 |
| siteLONDON | 1 | 36 | 0 | 0 | LsgACC | RdlPFC | 3.0339533 |
| siteLONDON | 1 | 36 | 0 | 0 | RsgACC | RdlPFC | 2.851169 |
| siteLONDON | 1 | 36 | 0 | 0 | LdlPFC | RdlPFC | 4.3493595 |
| siteLONDON | 1 | 36 | 0 | 0 | LNacc | mPFC | 4.340618 |
| siteLONDON | 1 | 36 | 0 | 0 | RNacc | mPFC | 2.9032476 |
| siteLONDON | 1 | 36 | 0 | 0 | LpACC | mPFC | 4.3152257 |
| siteLONDON | 1 | 36 | 0 | 0 | RpACC | mPFC | 2.5340244 |
| siteLONDON | 1 | 36 | 0 | 0 | LdlPFC | mPFC | 2.3191361 |
| siteLONDON | 1 | 36 | 0 | 0 | RdlPFC | mPFC | 1.9000947 |
| siteLONDON | 1 | 36 | 0 | 0 | LNacc | vmPFC | 3.1936347 |
| siteLONDON | 1 | 36 | 0 | 0 | RNacc | vmPFC | 2.6305602 |
| siteLONDON | 1 | 36 | 0 | 0 | LsgACC | vmPFC | 1.2124344 |
| siteLONDON | 1 | 36 | 0 | 0 | RsgACC | vmPFC | 1.8347281 |
| siteLONDON | 1 | 36 | 0 | 0 | LdlPFC | vmPFC | 1.0196131 |
| siteLONDON | 1 | 36 | 0 | 0 | RdlPFC | vmPFC | 0.7831319 |
| siteMANNHEIM | 1 | 23 | 0.0024 | 0.0002 | LNacc | LpACC | 0.8218353 |
| siteMANNHEIM | 1 | 23 | 0.0024 | 0.0002 | RNacc | LpACC | 0.6956514 |
| siteMANNHEIM | 1 | 23 | 0.0024 | 0.0002 | LNacc | RpACC | 0.9523134 |
| siteMANNHEIM | 1 | 23 | 0.0024 | 0.0002 | RNacc | RpACC | 1.1844427 |
| siteMANNHEIM | 1 | 23 | 0.0024 | 0.0002 | LpACC | LdlPFC | 1.2743181 |
| siteMANNHEIM | 1 | 23 | 0.0024 | 0.0002 | RpACC | LdlPFC | 2.1612965 |
| siteMANNHEIM | 1 | 23 | 0.0024 | 0.0002 | LsgACC | LdlPFC | 0.2260695 |
| siteMANNHEIM | 1 | 23 | 0.0024 | 0.0002 | RsgACC | LdlPFC | 1.9698739 |
| siteMANNHEIM | 1 | 23 | 0.0024 | 0.0002 | LNacc | RdlPFC | 0.0949193 |
| siteMANNHEIM | 1 | 23 | 0.0024 | 0.0002 | RNacc | RdlPFC | 1.0434548 |
| siteMANNHEIM | 1 | 23 | 0.0024 | 0.0002 | LpACC | RdlPFC | 0.9474849 |
| siteMANNHEIM | 1 | 23 | 0.0024 | 0.0002 | RpACC | RdlPFC | 2.5732554 |
| siteMANNHEIM | 1 | 23 | 0.0024 | 0.0002 | LsgACC | RdlPFC | 0.6943781 |
| siteMANNHEIM | 1 | 23 | 0.0024 | 0.0002 | RsgACC | RdlPFC | 1.2837461 |
| siteMANNHEIM | 1 | 23 | 0.0024 | 0.0002 | LdlPFC | RdlPFC | 2.1432265 |
| siteMANNHEIM | 1 | 23 | 0.0024 | 0.0002 | LNacc | mPFC | 0.6007128 |
| siteMANNHEIM | 1 | 23 | 0.0024 | 0.0002 | RNacc | mPFC | 0.87145 |
| siteMANNHEIM | 1 | 23 | 0.0024 | 0.0002 | LsgACC | mPFC | -0.1575704 |
| siteMANNHEIM | 1 | 23 | 0.0024 | 0.0002 | LdlPFC | mPFC | 2.065182 |
| siteMANNHEIM | 1 | 23 | 0.0024 | 0.0002 | RdlPFC | mPFC | 1.0858879 |
| siteMANNHEIM | 1 | 23 | 0.0024 | 0.0002 | RsgACC | vmPFC | -0.0050044 |
| siteMANNHEIM | 1 | 23 | 0.0024 | 0.0002 | LdlPFC | vmPFC | 1.9170394 |
| siteMANNHEIM | 1 | 23 | 0.0024 | 0.0002 | RdlPFC | vmPFC | 1.7577013 |
| siteNOTTINGHAM | 1 | 1 | 0.6196 | 0.127 | LNacc | RNacc | 1.9319901 |
| siteNOTTINGHAM | 1 | 1 | 0.6196 | 0.127 | LpACC | RpACC | 2.1867247 |
| siteNOTTINGHAM | 1 | 1 | 0.6196 | 0.127 | LpACC | LsgACC | 1.8796246 |
| siteNOTTINGHAM | 1 | 1 | 0.6196 | 0.127 | RpACC | LsgACC | 1.7233307 |
| siteNOTTINGHAM | 1 | 1 | 0.6196 | 0.127 | LpACC | RsgACC | 2.0030951 |
| siteNOTTINGHAM | 1 | 1 | 0.6196 | 0.127 | RpACC | RsgACC | 1.9007793 |
| siteNOTTINGHAM | 1 | 1 | 0.6196 | 0.127 | LsgACC | RsgACC | 2.523074 |
| siteNOTTINGHAM | 1 | 1 | 0.6196 | 0.127 | LpACC | mPFC | 2.0848241 |
| siteNOTTINGHAM | 1 | 1 | 0.6196 | 0.127 | RpACC | mPFC | 1.6757494 |
| siteNOTTINGHAM | 1 | 1 | 0.6196 | 0.127 | LsgACC | mPFC | 1.0563245 |
| siteNOTTINGHAM | 1 | 1 | 0.6196 | 0.127 | RsgACC | mPFC | 1.2601595 |
| siteNOTTINGHAM | 1 | 1 | 0.6196 | 0.127 | RpACC | vmPFC | 0.3229855 |
| siteNOTTINGHAM | 1 | 1 | 0.6196 | 0.127 | LsgACC | vmPFC | 1.1925697 |
| siteNOTTINGHAM | 1 | 1 | 0.6196 | 0.127 | RsgACC | vmPFC | 1.4557511 |
| siteNOTTINGHAM | 1 | 1 | 0.6196 | 0.127 | mPFC | vmPFC | 0.3460398 |
| sitePARIS | 1 | 11 | 0.028 | 0.02 | LNacc | RNacc | -0.2047912 |
| sitePARIS | 1 | 11 | 0.028 | 0.02 | LNacc | LpACC | 0.0034761 |
| sitePARIS | 1 | 11 | 0.028 | 0.02 | LpACC | LsgACC | -0.7809277 |
| sitePARIS | 1 | 11 | 0.028 | 0.02 | RpACC | LsgACC | -0.4581171 |
| sitePARIS | 1 | 11 | 0.028 | 0.02 | LpACC | RsgACC | -0.9093944 |
| sitePARIS | 1 | 11 | 0.028 | 0.02 | RpACC | RsgACC | -1.0588591 |
| sitePARIS | 1 | 11 | 0.028 | 0.02 | LsgACC | RsgACC | 0.3047263 |
| sitePARIS | 1 | 11 | 0.028 | 0.02 | LpACC | LdlPFC | 0.0276439 |
| sitePARIS | 1 | 11 | 0.028 | 0.02 | LsgACC | mPFC | -0.9049866 |
| sitePARIS | 1 | 11 | 0.028 | 0.02 | RsgACC | mPFC | -1.1676603 |
| sitePARIS | 1 | 11 | 0.028 | 0.02 | LpACC | vmPFC | -0.6385232 |
| mean_fd | 1 | 44 | 0 | 0 | LNacc | RNacc | 10.843918 |
| mean_fd | 1 | 44 | 0 | 0 | LNacc | LpACC | 5.6287803 |
| mean_fd | 1 | 44 | 0 | 0 | RNacc | LpACC | 6.4248849 |
| mean_fd | 1 | 44 | 0 | 0 | LNacc | RpACC | 6.4043721 |
| mean_fd | 1 | 44 | 0 | 0 | RNacc | RpACC | 5.9848622 |
| mean_fd | 1 | 44 | 0 | 0 | LpACC | RpACC | 6.6083014 |
| mean_fd | 1 | 44 | 0 | 0 | LNacc | LsgACC | 8.6340369 |
| mean_fd | 1 | 44 | 0 | 0 | RNacc | LsgACC | 7.9095953 |
| mean_fd | 1 | 44 | 0 | 0 | LpACC | LsgACC | 2.8131291 |
| mean_fd | 1 | 44 | 0 | 0 | RpACC | LsgACC | 3.2477454 |
| mean_fd | 1 | 44 | 0 | 0 | LNacc | RsgACC | 9.0626575 |
| mean_fd | 1 | 44 | 0 | 0 | RNacc | RsgACC | 7.87923 |
| mean_fd | 1 | 44 | 0 | 0 | LpACC | RsgACC | 3.1851487 |
| mean_fd | 1 | 44 | 0 | 0 | RpACC | RsgACC | 2.7195667 |
| mean_fd | 1 | 44 | 0 | 0 | LsgACC | RsgACC | 7.1972389 |
| mean_fd | 1 | 44 | 0 | 0 | LNacc | LdlPFC | 5.1144438 |
| mean_fd | 1 | 44 | 0 | 0 | RNacc | LdlPFC | 4.6750674 |
| mean_fd | 1 | 44 | 0 | 0 | LpACC | LdlPFC | 4.5950272 |
| mean_fd | 1 | 44 | 0 | 0 | RpACC | LdlPFC | 3.7489599 |
| mean_fd | 1 | 44 | 0 | 0 | LsgACC | LdlPFC | 6.2223231 |
| mean_fd | 1 | 44 | 0 | 0 | RsgACC | LdlPFC | 7.126192 |
| mean_fd | 1 | 44 | 0 | 0 | LNacc | RdlPFC | 5.2887888 |
| mean_fd | 1 | 44 | 0 | 0 | RNacc | RdlPFC | 5.1001 |
| mean_fd | 1 | 44 | 0 | 0 | LpACC | RdlPFC | 5.3215431 |
| mean_fd | 1 | 44 | 0 | 0 | RpACC | RdlPFC | 4.1669928 |
| mean_fd | 1 | 44 | 0 | 0 | LsgACC | RdlPFC | 5.9142838 |
| mean_fd | 1 | 44 | 0 | 0 | RsgACC | RdlPFC | 5.82483 |
| mean_fd | 1 | 44 | 0 | 0 | LNacc | mPFC | 5.8408514 |
| mean_fd | 1 | 44 | 0 | 0 | RNacc | mPFC | 6.5280813 |
| mean_fd | 1 | 44 | 0 | 0 | LpACC | mPFC | 3.0438564 |
| mean_fd | 1 | 44 | 0 | 0 | RpACC | mPFC | 3.9190576 |
| mean_fd | 1 | 44 | 0 | 0 | LsgACC | mPFC | 1.9657853 |
| mean_fd | 1 | 44 | 0 | 0 | RsgACC | mPFC | 1.693441 |
| mean_fd | 1 | 44 | 0 | 0 | LdlPFC | mPFC | 4.5856249 |
| mean_fd | 1 | 44 | 0 | 0 | RdlPFC | mPFC | 5.1279768 |
| mean_fd | 1 | 44 | 0 | 0 | LNacc | vmPFC | 8.8332664 |
| mean_fd | 1 | 44 | 0 | 0 | RNacc | vmPFC | 8.6408801 |
| mean_fd | 1 | 44 | 0 | 0 | LpACC | vmPFC | 1.1636529 |
| mean_fd | 1 | 44 | 0 | 0 | RpACC | vmPFC | 1.4882813 |
| mean_fd | 1 | 44 | 0 | 0 | LsgACC | vmPFC | 5.3783632 |
| mean_fd | 1 | 44 | 0 | 0 | RsgACC | vmPFC | 3.9077109 |
| mean_fd | 1 | 44 | 0 | 0 | LdlPFC | vmPFC | 4.1473038 |
| mean_fd | 1 | 44 | 0 | 0 | RdlPFC | vmPFC | 5.1664214 |
| mean_fd | 1 | 44 | 0 | 0 | mPFC | vmPFC | 1.1701332 |
| threat_z:SexMale | 6 | 1 | 0.613 | 0.5892 | RsgACC | LdlPFC | 0.0299003 |
| threat_z:SexMale | 6 | 1 | 0.613 | 0.5892 | RdlPFC | vmPFC | -0.1438181 |
| SexMale:deprivation_z | 1 | 9 | 0.047 | 0.0384 | LNacc | RNacc | 0.848498 |
| SexMale:deprivation_z | 1 | 9 | 0.047 | 0.0384 | RNacc | LpACC | 0.2426219 |
| SexMale:deprivation_z | 1 | 9 | 0.047 | 0.0384 | RNacc | RpACC | 0.0291526 |
| SexMale:deprivation_z | 1 | 9 | 0.047 | 0.0384 | LNacc | RdlPFC | 0.3658004 |
| SexMale:deprivation_z | 1 | 9 | 0.047 | 0.0384 | RNacc | RdlPFC | 1.0138737 |
| SexMale:deprivation_z | 1 | 9 | 0.047 | 0.0384 | RNacc | mPFC | 0.1897306 |
| SexMale:deprivation_z | 1 | 9 | 0.047 | 0.0384 | LNacc | vmPFC | 0.0487816 |
| SexMale:deprivation_z | 1 | 9 | 0.047 | 0.0384 | RNacc | vmPFC | 0.4596471 |
| SexMale:deprivation_z | 1 | 9 | 0.047 | 0.0384 | RdlPFC | vmPFC | 1.1103929 |

**Table S5:** *Longitudinal Male model (CDT: p<0.05)*

| Variable | Cluster | No. Edges | p-FWE (Size) | p-FWE (Strength) | ROI1 | ROI2 | Strength |
| --- | --- | --- | --- | --- | --- | --- | --- |
| timepointFU3 | 1 | 17 | 0.0074 | 0.0008 | LNacc | RNacc | -2.5002782 |
| timepointFU3 | 1 | 17 | 0.0074 | 0.0008 | LNacc | LpACC | -0.3438912 |
| timepointFU3 | 1 | 17 | 0.0074 | 0.0008 | RNacc | LpACC | -0.8242203 |
| timepointFU3 | 1 | 17 | 0.0074 | 0.0008 | LNacc | RpACC | -0.6998232 |
| timepointFU3 | 1 | 17 | 0.0074 | 0.0008 | RNacc | RpACC | -1.2331906 |
| timepointFU3 | 1 | 17 | 0.0074 | 0.0008 | LNacc | LsgACC | -0.2689943 |
| timepointFU3 | 1 | 17 | 0.0074 | 0.0008 | RNacc | LsgACC | -0.6201477 |
| timepointFU3 | 1 | 17 | 0.0074 | 0.0008 | LNacc | RsgACC | -2.4303644 |
| timepointFU3 | 1 | 17 | 0.0074 | 0.0008 | RNacc | RsgACC | -2.551971 |
| timepointFU3 | 1 | 17 | 0.0074 | 0.0008 | LsgACC | RsgACC | -3.9338968 |
| timepointFU3 | 1 | 17 | 0.0074 | 0.0008 | LpACC | RdlPFC | 0.5118437 |
| timepointFU3 | 1 | 17 | 0.0074 | 0.0008 | RpACC | RdlPFC | 0.2075295 |
| timepointFU3 | 1 | 17 | 0.0074 | 0.0008 | LNacc | mPFC | -0.3956789 |
| timepointFU3 | 1 | 17 | 0.0074 | 0.0008 | RNacc | mPFC | -0.0417044 |
| timepointFU3 | 1 | 17 | 0.0074 | 0.0008 | LNacc | vmPFC | -0.4768363 |
| timepointFU3 | 1 | 17 | 0.0074 | 0.0008 | RNacc | vmPFC | -0.7469811 |
| timepointFU3 | 1 | 17 | 0.0074 | 0.0008 | RsgACC | vmPFC | -0.4445349 |
| threat_z | 6 | 1 | 0.4226 | 0.2052 | RsgACC | LdlPFC | 0.2970969 |
| deprivation_z | 8 | 1 | 0.5708 | 0.3982 | RdlPFC | vmPFC | 0.1941229 |
| mean_fd | 1 | 44 | 0.0568 | 0.0012 | LNacc | RNacc | 6.6355609 |
| mean_fd | 1 | 44 | 0.0568 | 0.0012 | LNacc | LpACC | 3.9977214 |
| mean_fd | 1 | 44 | 0.0568 | 0.0012 | RNacc | LpACC | 3.2610743 |
| mean_fd | 1 | 44 | 0.0568 | 0.0012 | LNacc | RpACC | 4.6453651 |
| mean_fd | 1 | 44 | 0.0568 | 0.0012 | RNacc | RpACC | 4.3018374 |
| mean_fd | 1 | 44 | 0.0568 | 0.0012 | LpACC | RpACC | 4.4897983 |
| mean_fd | 1 | 44 | 0.0568 | 0.0012 | LNacc | LsgACC | 4.7586814 |
| mean_fd | 1 | 44 | 0.0568 | 0.0012 | RNacc | LsgACC | 4.7607921 |
| mean_fd | 1 | 44 | 0.0568 | 0.0012 | LpACC | LsgACC | 1.1840776 |
| mean_fd | 1 | 44 | 0.0568 | 0.0012 | RpACC | LsgACC | 2.7881622 |
| mean_fd | 1 | 44 | 0.0568 | 0.0012 | LNacc | RsgACC | 5.2987094 |
| mean_fd | 1 | 44 | 0.0568 | 0.0012 | RNacc | RsgACC | 5.0251156 |
| mean_fd | 1 | 44 | 0.0568 | 0.0012 | LpACC | RsgACC | 1.679708 |
| mean_fd | 1 | 44 | 0.0568 | 0.0012 | RpACC | RsgACC | 2.3718055 |
| mean_fd | 1 | 44 | 0.0568 | 0.0012 | LsgACC | RsgACC | 3.1251005 |
| mean_fd | 1 | 44 | 0.0568 | 0.0012 | LNacc | LdlPFC | 0.7934276 |
| mean_fd | 1 | 44 | 0.0568 | 0.0012 | RNacc | LdlPFC | 1.4376102 |
| mean_fd | 1 | 44 | 0.0568 | 0.0012 | LpACC | LdlPFC | 1.5925302 |
| mean_fd | 1 | 44 | 0.0568 | 0.0012 | RpACC | LdlPFC | 1.0282169 |
| mean_fd | 1 | 44 | 0.0568 | 0.0012 | LsgACC | LdlPFC | 1.747473 |
| mean_fd | 1 | 44 | 0.0568 | 0.0012 | RsgACC | LdlPFC | 1.2306461 |
| mean_fd | 1 | 44 | 0.0568 | 0.0012 | LNacc | RdlPFC | 1.3559921 |
| mean_fd | 1 | 44 | 0.0568 | 0.0012 | RNacc | RdlPFC | 0.5966538 |
| mean_fd | 1 | 44 | 0.0568 | 0.0012 | LpACC | RdlPFC | 2.5092695 |
| mean_fd | 1 | 44 | 0.0568 | 0.0012 | RpACC | RdlPFC | 1.2893092 |
| mean_fd | 1 | 44 | 0.0568 | 0.0012 | LsgACC | RdlPFC | 1.2260409 |
| mean_fd | 1 | 44 | 0.0568 | 0.0012 | RsgACC | RdlPFC | 1.0844336 |
| mean_fd | 1 | 44 | 0.0568 | 0.0012 | LNacc | mPFC | 2.8136513 |
| mean_fd | 1 | 44 | 0.0568 | 0.0012 | RNacc | mPFC | 2.0412818 |
| mean_fd | 1 | 44 | 0.0568 | 0.0012 | LpACC | mPFC | 0.5743274 |
| mean_fd | 1 | 44 | 0.0568 | 0.0012 | RpACC | mPFC | 1.0824498 |
| mean_fd | 1 | 44 | 0.0568 | 0.0012 | LsgACC | mPFC | 1.1926133 |
| mean_fd | 1 | 44 | 0.0568 | 0.0012 | RsgACC | mPFC | 0.7011259 |
| mean_fd | 1 | 44 | 0.0568 | 0.0012 | LdlPFC | mPFC | 1.6742104 |
| mean_fd | 1 | 44 | 0.0568 | 0.0012 | RdlPFC | mPFC | 2.5133146 |
| mean_fd | 1 | 44 | 0.0568 | 0.0012 | LNacc | vmPFC | 5.2575589 |
| mean_fd | 1 | 44 | 0.0568 | 0.0012 | RNacc | vmPFC | 5.0390205 |
| mean_fd | 1 | 44 | 0.0568 | 0.0012 | LpACC | vmPFC | 1.1455999 |
| mean_fd | 1 | 44 | 0.0568 | 0.0012 | RpACC | vmPFC | 2.0998061 |
| mean_fd | 1 | 44 | 0.0568 | 0.0012 | LsgACC | vmPFC | 3.4539484 |
| mean_fd | 1 | 44 | 0.0568 | 0.0012 | RsgACC | vmPFC | 2.6869869 |
| mean_fd | 1 | 44 | 0.0568 | 0.0012 | LdlPFC | vmPFC | 1.6380732 |
| mean_fd | 1 | 44 | 0.0568 | 0.0012 | RdlPFC | vmPFC | 1.6521554 |
| mean_fd | 1 | 44 | 0.0568 | 0.0012 | mPFC | vmPFC | 0.7571225 |
| siteDUBLIN | 1 | 1 | 1 | 1 | LNacc | RNacc | 0.9316909 |
| siteDUBLIN | 1 | 1 | 1 | 1 | LpACC | RpACC | 1.7084576 |
| siteDUBLIN | 1 | 1 | 1 | 1 | LpACC | LsgACC | 0.5056047 |
| siteDUBLIN | 1 | 1 | 1 | 1 | RpACC | LsgACC | 0.4134765 |
| siteDUBLIN | 1 | 1 | 1 | 1 | LpACC | RsgACC | 0.1263315 |
| siteDUBLIN | 1 | 1 | 1 | 1 | RpACC | RsgACC | 0.6075797 |
| siteDUBLIN | 1 | 1 | 1 | 1 | LdlPFC | RdlPFC | 0.8063516 |
| siteDUBLIN | 1 | 1 | 1 | 1 | LpACC | mPFC | 2.3057265 |
| siteDUBLIN | 1 | 1 | 1 | 1 | RpACC | mPFC | 1.1761783 |
| siteDUBLIN | 1 | 1 | 1 | 1 | LsgACC | mPFC | 0.3688082 |
| siteDUBLIN | 1 | 1 | 1 | 1 | LpACC | vmPFC | 1.1001427 |
| siteDUBLIN | 1 | 1 | 1 | 1 | RpACC | vmPFC | 1.7113915 |
| siteDUBLIN | 1 | 1 | 1 | 1 | mPFC | vmPFC | 0.9862247 |
| siteHAMBURG | 1 | 11 | 0.5954 | 0.0064 | LNacc | LpACC | 0.1235471 |
| siteHAMBURG | 1 | 11 | 0.5954 | 0.0064 | LNacc | RpACC | 0.4682508 |
| siteHAMBURG | 1 | 11 | 0.5954 | 0.0064 | LpACC | RpACC | 0.1793688 |
| siteHAMBURG | 1 | 11 | 0.5954 | 0.0064 | LNacc | LdlPFC | 0.0746621 |
| siteHAMBURG | 1 | 11 | 0.5954 | 0.0064 | RpACC | LdlPFC | 0.362334 |
| siteHAMBURG | 1 | 11 | 0.5954 | 0.0064 | LpACC | RdlPFC | 0.1998896 |
| siteHAMBURG | 1 | 11 | 0.5954 | 0.0064 | RpACC | RdlPFC | 0.7081157 |
| siteHAMBURG | 1 | 11 | 0.5954 | 0.0064 | LdlPFC | RdlPFC | 2.2935584 |
| siteHAMBURG | 1 | 11 | 0.5954 | 0.0064 | LNacc | mPFC | 0.0634945 |
| siteHAMBURG | 1 | 11 | 0.5954 | 0.0064 | LpACC | mPFC | 0.4560082 |
| siteHAMBURG | 1 | 11 | 0.5954 | 0.0064 | RdlPFC | mPFC | 0.2063775 |
| siteLONDON | 1 | 26 | 0.9998 | 0.9852 | LNacc | RNacc | 2.9065467 |
| siteLONDON | 1 | 26 | 0.9998 | 0.9852 | LNacc | LpACC | 0.5475494 |
| siteLONDON | 1 | 26 | 0.9998 | 0.9852 | LNacc | RpACC | 0.3097 |
| siteLONDON | 1 | 26 | 0.9998 | 0.9852 | RNacc | RpACC | 0.0764475 |
| siteLONDON | 1 | 26 | 0.9998 | 0.9852 | LpACC | RpACC | 1.876716 |
| siteLONDON | 1 | 26 | 0.9998 | 0.9852 | LNacc | RsgACC | 0.2388659 |
| siteLONDON | 1 | 26 | 0.9998 | 0.9852 | RNacc | RsgACC | 0.1612069 |
| siteLONDON | 1 | 26 | 0.9998 | 0.9852 | LsgACC | RsgACC | 1.001035 |
| siteLONDON | 1 | 26 | 0.9998 | 0.9852 | LNacc | LdlPFC | 0.6536319 |
| siteLONDON | 1 | 26 | 0.9998 | 0.9852 | RNacc | LdlPFC | 0.1271579 |
| siteLONDON | 1 | 26 | 0.9998 | 0.9852 | RpACC | LdlPFC | 0.38501 |
| siteLONDON | 1 | 26 | 0.9998 | 0.9852 | LsgACC | LdlPFC | 0.3077823 |
| siteLONDON | 1 | 26 | 0.9998 | 0.9852 | RsgACC | LdlPFC | 0.5855099 |
| siteLONDON | 1 | 26 | 0.9998 | 0.9852 | LNacc | RdlPFC | 0.2730982 |
| siteLONDON | 1 | 26 | 0.9998 | 0.9852 | LpACC | RdlPFC | 0.4132867 |
| siteLONDON | 1 | 26 | 0.9998 | 0.9852 | RpACC | RdlPFC | 1.1321888 |
| siteLONDON | 1 | 26 | 0.9998 | 0.9852 | LsgACC | RdlPFC | 0.1373557 |
| siteLONDON | 1 | 26 | 0.9998 | 0.9852 | RsgACC | RdlPFC | 0.4367257 |
| siteLONDON | 1 | 26 | 0.9998 | 0.9852 | LdlPFC | RdlPFC | 0.8151202 |
| siteLONDON | 1 | 26 | 0.9998 | 0.9852 | LNacc | mPFC | 0.1815396 |
| siteLONDON | 1 | 26 | 0.9998 | 0.9852 | LpACC | mPFC | 1.0778103 |
| siteLONDON | 1 | 26 | 0.9998 | 0.9852 | RpACC | mPFC | 0.2464384 |
| siteLONDON | 1 | 26 | 0.9998 | 0.9852 | LdlPFC | mPFC | 0.1108061 |
| siteLONDON | 1 | 26 | 0.9998 | 0.9852 | RdlPFC | mPFC | 0.3750218 |
| siteLONDON | 1 | 26 | 0.9998 | 0.9852 | LNacc | vmPFC | 0.228995 |
| siteLONDON | 1 | 26 | 0.9998 | 0.9852 | RNacc | vmPFC | 0.0974887 |
| siteMANNHEIM | 7 | 1 | 1 | 0.8566 | LdlPFC | RdlPFC | 0.3981237 |
| siteNOTTINGHAM | 1 | 13 | 0.6514 | 0.9662 | LNacc | RNacc | 0.9414668 |
| siteNOTTINGHAM | 1 | 13 | 0.6514 | 0.9662 | LNacc | LpACC | 0.0342047 |
| siteNOTTINGHAM | 1 | 13 | 0.6514 | 0.9662 | LpACC | RpACC | 0.8161472 |
| siteNOTTINGHAM | 1 | 13 | 0.6514 | 0.9662 | LpACC | LsgACC | 0.9885312 |
| siteNOTTINGHAM | 1 | 13 | 0.6514 | 0.9662 | RpACC | LsgACC | 1.3845916 |
| siteNOTTINGHAM | 1 | 13 | 0.6514 | 0.9662 | LpACC | RsgACC | 1.2614534 |
| siteNOTTINGHAM | 1 | 13 | 0.6514 | 0.9662 | RpACC | RsgACC | 1.0050825 |
| siteNOTTINGHAM | 1 | 13 | 0.6514 | 0.9662 | LNacc | LdlPFC | 0.1717734 |
| siteNOTTINGHAM | 1 | 13 | 0.6514 | 0.9662 | LdlPFC | RdlPFC | 0.0392357 |
| siteNOTTINGHAM | 1 | 13 | 0.6514 | 0.9662 | LpACC | mPFC | 0.8068087 |
| siteNOTTINGHAM | 1 | 13 | 0.6514 | 0.9662 | RpACC | mPFC | 0.2180479 |
| siteNOTTINGHAM | 1 | 13 | 0.6514 | 0.9662 | LpACC | vmPFC | 0.4048144 |
| siteNOTTINGHAM | 1 | 13 | 0.6514 | 0.9662 | RpACC | vmPFC | 0.463899 |
| sitePARIS | 1 | 9 | 0.8988 | 0.0106 | LNacc | LpACC | 0.3748513 |
| sitePARIS | 1 | 9 | 0.8988 | 0.0106 | RNacc | LpACC | 0.1959389 |
| sitePARIS | 1 | 9 | 0.8988 | 0.0106 | LNacc | RpACC | 0.4114475 |
| sitePARIS | 1 | 9 | 0.8988 | 0.0106 | RNacc | RpACC | 0.1170282 |
| sitePARIS | 1 | 9 | 0.8988 | 0.0106 | LdlPFC | RdlPFC | 0.6581298 |
| sitePARIS | 1 | 9 | 0.8988 | 0.0106 | LNacc | mPFC | 0.5529618 |
| sitePARIS | 1 | 9 | 0.8988 | 0.0106 | RNacc | mPFC | 0.4348864 |
| sitePARIS | 1 | 9 | 0.8988 | 0.0106 | LpACC | vmPFC | 0.1666068 |
| sitePARIS | 1 | 9 | 0.8988 | 0.0106 | RpACC | vmPFC | 1.2990503 |
| sitePARIS | 1 | 9 | 0.8988 | 0.0106 | mPFC | vmPFC | 0.7504074 |
| timepointFU3:deprivation_z | 1 | 1 | 0.5698 | 0.345 | LNacc | RNacc | -0.3523139 |

**Table S6:** *Longitudinal Female model (CDT: p<0.05)*

| Variable | Cluster | No. Edges | p-FWE (Size) | p-FWE (Strength) | ROI1 | ROI2 | Strength |
| --- | --- | --- | --- | --- | --- | --- | --- |
| timepointFU3 | 2 | 1 | 0.6774 | 0.6142 | LpACC | RpACC | -1.0081395 |
| timepointFU3 | 2 | 1 | 0.6774 | 0.6142 | LsgACC | RsgACC | -3.5898594 |
| timepointFU3 | 2 | 1 | 0.6774 | 0.6142 | RNacc | LdlPFC | -0.0922634 |
| threat_z | 1 | 18 | 0.0188 | 0.0528 | LNacc | LpACC | 0.3431063 |
| threat_z | 1 | 18 | 0.0188 | 0.0528 | RNacc | LpACC | 0.2286175 |
| threat_z | 1 | 18 | 0.0188 | 0.0528 | LNacc | RpACC | 0.2471142 |
| threat_z | 1 | 18 | 0.0188 | 0.0528 | RNacc | RpACC | 0.1268503 |
| threat_z | 1 | 18 | 0.0188 | 0.0528 | LNacc | LsgACC | 2.0552636 |
| threat_z | 1 | 18 | 0.0188 | 0.0528 | RNacc | LsgACC | 1.1857733 |
| threat_z | 1 | 18 | 0.0188 | 0.0528 | LpACC | LsgACC | 0.5916031 |
| threat_z | 1 | 18 | 0.0188 | 0.0528 | RpACC | LsgACC | 0.2473981 |
| threat_z | 1 | 18 | 0.0188 | 0.0528 | LNacc | RsgACC | 0.7994222 |
| threat_z | 1 | 18 | 0.0188 | 0.0528 | RNacc | RsgACC | 0.0510349 |
| threat_z | 1 | 18 | 0.0188 | 0.0528 | LpACC | RsgACC | 0.0779395 |
| threat_z | 1 | 18 | 0.0188 | 0.0528 | RNacc | mPFC | 0.385907 |
| threat_z | 1 | 18 | 0.0188 | 0.0528 | LsgACC | mPFC | 0.8737063 |
| threat_z | 1 | 18 | 0.0188 | 0.0528 | RsgACC | mPFC | 0.6037124 |
| threat_z | 1 | 18 | 0.0188 | 0.0528 | LNacc | vmPFC | 1.0777085 |
| threat_z | 1 | 18 | 0.0188 | 0.0528 | RNacc | vmPFC | 1.2369662 |
| threat_z | 1 | 18 | 0.0188 | 0.0528 | LsgACC | vmPFC | 0.4515195 |
| threat_z | 1 | 18 | 0.0188 | 0.0528 | mPFC | vmPFC | 0.0559259 |
| deprivation_z | 1 | 22 | 0.0022 | 0.004 | LNacc | RNacc | -0.0744958 |
| deprivation_z | 1 | 22 | 0.0022 | 0.004 | LNacc | LpACC | -0.3957424 |
| deprivation_z | 1 | 22 | 0.0022 | 0.004 | LNacc | RpACC | -0.872973 |
| deprivation_z | 1 | 22 | 0.0022 | 0.004 | RNacc | RpACC | -0.3306594 |
| deprivation_z | 1 | 22 | 0.0022 | 0.004 | LpACC | RpACC | -0.1531164 |
| deprivation_z | 1 | 22 | 0.0022 | 0.004 | LNacc | LsgACC | -1.0602637 |
| deprivation_z | 1 | 22 | 0.0022 | 0.004 | RNacc | LsgACC | -0.7490921 |
| deprivation_z | 1 | 22 | 0.0022 | 0.004 | LpACC | LsgACC | -1.6892049 |
| deprivation_z | 1 | 22 | 0.0022 | 0.004 | RpACC | LsgACC | -1.1966695 |
| deprivation_z | 1 | 22 | 0.0022 | 0.004 | RNacc | RsgACC | -0.2094267 |
| deprivation_z | 1 | 22 | 0.0022 | 0.004 | LpACC | RsgACC | -0.6802479 |
| deprivation_z | 1 | 22 | 0.0022 | 0.004 | RpACC | RsgACC | -0.6658667 |
| deprivation_z | 1 | 22 | 0.0022 | 0.004 | LsgACC | RsgACC | -0.0588071 |
| deprivation_z | 1 | 22 | 0.0022 | 0.004 | RNacc | RdlPFC | -0.2468973 |
| deprivation_z | 1 | 22 | 0.0022 | 0.004 | LsgACC | RdlPFC | -0.47161 |
| deprivation_z | 1 | 22 | 0.0022 | 0.004 | LNacc | mPFC | -0.4596445 |
| deprivation_z | 1 | 22 | 0.0022 | 0.004 | RNacc | mPFC | -0.6659858 |
| deprivation_z | 1 | 22 | 0.0022 | 0.004 | RpACC | mPFC | -0.0608718 |
| deprivation_z | 1 | 22 | 0.0022 | 0.004 | LsgACC | mPFC | -1.7588239 |
| deprivation_z | 1 | 22 | 0.0022 | 0.004 | RsgACC | mPFC | -1.0143865 |
| deprivation_z | 1 | 22 | 0.0022 | 0.004 | RNacc | vmPFC | -0.1064311 |
| deprivation_z | 1 | 22 | 0.0022 | 0.004 | LsgACC | vmPFC | -0.4643591 |
| mean_fd | 1 | 42 | 0.0934 | 0.0008 | LNacc | RNacc | 7.1099473 |
| mean_fd | 1 | 42 | 0.0934 | 0.0008 | LNacc | LpACC | 3.9878472 |
| mean_fd | 1 | 42 | 0.0934 | 0.0008 | RNacc | LpACC | 2.0050033 |
| mean_fd | 1 | 42 | 0.0934 | 0.0008 | LNacc | RpACC | 2.9537199 |
| mean_fd | 1 | 42 | 0.0934 | 0.0008 | RNacc | RpACC | 1.2808196 |
| mean_fd | 1 | 42 | 0.0934 | 0.0008 | LpACC | RpACC | 4.226456 |
| mean_fd | 1 | 42 | 0.0934 | 0.0008 | LNacc | LsgACC | 6.1851801 |
| mean_fd | 1 | 42 | 0.0934 | 0.0008 | RNacc | LsgACC | 5.6722134 |
| mean_fd | 1 | 42 | 0.0934 | 0.0008 | LpACC | LsgACC | 1.9523382 |
| mean_fd | 1 | 42 | 0.0934 | 0.0008 | RpACC | LsgACC | 0.7922802 |
| mean_fd | 1 | 42 | 0.0934 | 0.0008 | LNacc | RsgACC | 4.5875577 |
| mean_fd | 1 | 42 | 0.0934 | 0.0008 | RNacc | RsgACC | 4.7483325 |
| mean_fd | 1 | 42 | 0.0934 | 0.0008 | LpACC | RsgACC | 0.9399221 |
| mean_fd | 1 | 42 | 0.0934 | 0.0008 | RpACC | RsgACC | 0.2333218 |
| mean_fd | 1 | 42 | 0.0934 | 0.0008 | LsgACC | RsgACC | 5.3089757 |
| mean_fd | 1 | 42 | 0.0934 | 0.0008 | LNacc | LdlPFC | 2.257496 |
| mean_fd | 1 | 42 | 0.0934 | 0.0008 | RNacc | LdlPFC | 1.5490737 |
| mean_fd | 1 | 42 | 0.0934 | 0.0008 | LpACC | LdlPFC | 3.549098 |
| mean_fd | 1 | 42 | 0.0934 | 0.0008 | RpACC | LdlPFC | 2.8918836 |
| mean_fd | 1 | 42 | 0.0934 | 0.0008 | LsgACC | LdlPFC | 2.6314189 |
| mean_fd | 1 | 42 | 0.0934 | 0.0008 | RsgACC | LdlPFC | 1.8793466 |
| mean_fd | 1 | 42 | 0.0934 | 0.0008 | LNacc | RdlPFC | 2.2191406 |
| mean_fd | 1 | 42 | 0.0934 | 0.0008 | RNacc | RdlPFC | 2.88277 |
| mean_fd | 1 | 42 | 0.0934 | 0.0008 | LpACC | RdlPFC | 4.1531007 |
| mean_fd | 1 | 42 | 0.0934 | 0.0008 | RpACC | RdlPFC | 1.7603658 |
| mean_fd | 1 | 42 | 0.0934 | 0.0008 | LsgACC | RdlPFC | 3.3902347 |
| mean_fd | 1 | 42 | 0.0934 | 0.0008 | RsgACC | RdlPFC | 2.0856159 |
| mean_fd | 1 | 42 | 0.0934 | 0.0008 | LNacc | mPFC | 4.360861 |
| mean_fd | 1 | 42 | 0.0934 | 0.0008 | RNacc | mPFC | 2.820591 |
| mean_fd | 1 | 42 | 0.0934 | 0.0008 | LpACC | mPFC | 3.3591266 |
| mean_fd | 1 | 42 | 0.0934 | 0.0008 | RpACC | mPFC | 2.7900868 |
| mean_fd | 1 | 42 | 0.0934 | 0.0008 | LsgACC | mPFC | 1.1669742 |
| mean_fd | 1 | 42 | 0.0934 | 0.0008 | LdlPFC | mPFC | 3.4019302 |
| mean_fd | 1 | 42 | 0.0934 | 0.0008 | RdlPFC | mPFC | 3.7142922 |
| mean_fd | 1 | 42 | 0.0934 | 0.0008 | LNacc | vmPFC | 3.6893233 |
| mean_fd | 1 | 42 | 0.0934 | 0.0008 | RNacc | vmPFC | 4.3604771 |
| mean_fd | 1 | 42 | 0.0934 | 0.0008 | LpACC | vmPFC | 1.3553369 |
| mean_fd | 1 | 42 | 0.0934 | 0.0008 | LsgACC | vmPFC | 4.6818606 |
| mean_fd | 1 | 42 | 0.0934 | 0.0008 | RsgACC | vmPFC | 3.0758673 |
| mean_fd | 1 | 42 | 0.0934 | 0.0008 | LdlPFC | vmPFC | 1.7620298 |
| mean_fd | 1 | 42 | 0.0934 | 0.0008 | RdlPFC | vmPFC | 1.5029928 |
| mean_fd | 1 | 42 | 0.0934 | 0.0008 | mPFC | vmPFC | 0.027852 |
| siteDRESDEN | 3 | 3 | 0.9912 | 0.833 | LpACC | RsgACC | -0.2366493 |
| siteDRESDEN | 3 | 3 | 0.9912 | 0.833 | RpACC | RsgACC | -0.1285041 |
| siteDRESDEN | 3 | 3 | 0.9912 | 0.833 | RsgACC | mPFC | -0.4024765 |
| siteDRESDEN | 3 | 3 | 0.9912 | 0.833 | LdlPFC | vmPFC | 0.3611885 |
| siteDRESDEN | 3 | 3 | 0.9912 | 0.833 | RdlPFC | vmPFC | 0.3744561 |
| siteDUBLIN | 1 | 15 | 0.8478 | 0.7918 | LNacc | RNacc | 1.5658523 |
| siteDUBLIN | 1 | 15 | 0.8478 | 0.7918 | LNacc | RpACC | 0.6807217 |
| siteDUBLIN | 1 | 15 | 0.8478 | 0.7918 | RNacc | RpACC | 0.0936522 |
| siteDUBLIN | 1 | 15 | 0.8478 | 0.7918 | LpACC | RpACC | 2.4346322 |
| siteDUBLIN | 1 | 15 | 0.8478 | 0.7918 | LpACC | LsgACC | 1.4411497 |
| siteDUBLIN | 1 | 15 | 0.8478 | 0.7918 | RpACC | LsgACC | 2.178337 |
| siteDUBLIN | 1 | 15 | 0.8478 | 0.7918 | LpACC | RsgACC | 0.7603742 |
| siteDUBLIN | 1 | 15 | 0.8478 | 0.7918 | RpACC | RsgACC | 1.9130584 |
| siteDUBLIN | 1 | 15 | 0.8478 | 0.7918 | LsgACC | RsgACC | 1.8199884 |
| siteDUBLIN | 1 | 15 | 0.8478 | 0.7918 | LNacc | LdlPFC | 0.0099204 |
| siteDUBLIN | 1 | 15 | 0.8478 | 0.7918 | LdlPFC | RdlPFC | 2.0703803 |
| siteDUBLIN | 1 | 15 | 0.8478 | 0.7918 | LpACC | mPFC | 1.8919471 |
| siteDUBLIN | 1 | 15 | 0.8478 | 0.7918 | RpACC | mPFC | 0.844686 |
| siteDUBLIN | 1 | 15 | 0.8478 | 0.7918 | LsgACC | mPFC | 0.3728932 |
| siteDUBLIN | 1 | 15 | 0.8478 | 0.7918 | LsgACC | vmPFC | 0.5801241 |
| siteHAMBURG | 3 | 8 | 0.9994 | 0.0926 | LpACC | RsgACC | -0.6664888 |
| siteHAMBURG | 3 | 8 | 0.9994 | 0.0926 | LpACC | LdlPFC | 1.336992 |
| siteHAMBURG | 3 | 8 | 0.9994 | 0.0926 | RpACC | LdlPFC | 1.8691289 |
| siteHAMBURG | 3 | 8 | 0.9994 | 0.0926 | LdlPFC | RdlPFC | 1.080496 |
| siteHAMBURG | 3 | 8 | 0.9994 | 0.0926 | LsgACC | mPFC | -0.4259486 |
| siteHAMBURG | 3 | 8 | 0.9994 | 0.0926 | RsgACC | mPFC | -0.6374889 |
| siteHAMBURG | 3 | 8 | 0.9994 | 0.0926 | LdlPFC | mPFC | 2.2723568 |
| siteHAMBURG | 3 | 8 | 0.9994 | 0.0926 | LdlPFC | vmPFC | 1.032881 |
| siteLONDON | 1 | 20 | 1 | 0.9918 | LNacc | RNacc | 4.9874699 |
| siteLONDON | 1 | 20 | 1 | 0.9918 | LNacc | LpACC | 0.5245046 |
| siteLONDON | 1 | 20 | 1 | 0.9918 | LNacc | RpACC | 0.9759203 |
| siteLONDON | 1 | 20 | 1 | 0.9918 | RNacc | RpACC | 0.7256151 |
| siteLONDON | 1 | 20 | 1 | 0.9918 | LpACC | RpACC | 2.5326062 |
| siteLONDON | 1 | 20 | 1 | 0.9918 | LNacc | LsgACC | 0.4594279 |
| siteLONDON | 1 | 20 | 1 | 0.9918 | RNacc | LsgACC | 0.3998251 |
| siteLONDON | 1 | 20 | 1 | 0.9918 | LNacc | RsgACC | 0.5939565 |
| siteLONDON | 1 | 20 | 1 | 0.9918 | RNacc | RsgACC | 0.1087372 |
| siteLONDON | 1 | 20 | 1 | 0.9918 | LsgACC | RsgACC | 3.833925 |
| siteLONDON | 1 | 20 | 1 | 0.9918 | LpACC | LdlPFC | 0.32398 |
| siteLONDON | 1 | 20 | 1 | 0.9918 | RpACC | LdlPFC | 1.1497969 |
| siteLONDON | 1 | 20 | 1 | 0.9918 | LsgACC | LdlPFC | 1.0617437 |
| siteLONDON | 1 | 20 | 1 | 0.9918 | RsgACC | LdlPFC | 1.5522754 |
| siteLONDON | 1 | 20 | 1 | 0.9918 | RNacc | RdlPFC | 0.095913 |
| siteLONDON | 1 | 20 | 1 | 0.9918 | LsgACC | RdlPFC | 0.4020972 |
| siteLONDON | 1 | 20 | 1 | 0.9918 | RsgACC | RdlPFC | 0.3111637 |
| siteLONDON | 1 | 20 | 1 | 0.9918 | LdlPFC | RdlPFC | 1.9543613 |
| siteLONDON | 1 | 20 | 1 | 0.9918 | LpACC | mPFC | 2.5569729 |
| siteLONDON | 1 | 20 | 1 | 0.9918 | LdlPFC | vmPFC | 0.4104831 |
| siteMANNHEIM | 1 | 20 | 0.478 | 0.9732 | LNacc | LpACC | 0.3836287 |
| siteMANNHEIM | 1 | 20 | 0.478 | 0.9732 | LNacc | RpACC | 0.387726 |
| siteMANNHEIM | 1 | 20 | 0.478 | 0.9732 | LpACC | RsgACC | -0.1776001 |
| siteMANNHEIM | 1 | 20 | 0.478 | 0.9732 | LpACC | LdlPFC | 1.9147932 |
| siteMANNHEIM | 1 | 20 | 0.478 | 0.9732 | RpACC | LdlPFC | 2.3894317 |
| siteMANNHEIM | 1 | 20 | 0.478 | 0.9732 | LsgACC | LdlPFC | 0.7529972 |
| siteMANNHEIM | 1 | 20 | 0.478 | 0.9732 | RsgACC | LdlPFC | 1.0287114 |
| siteMANNHEIM | 1 | 20 | 0.478 | 0.9732 | RNacc | RdlPFC | 0.305078 |
| siteMANNHEIM | 1 | 20 | 0.478 | 0.9732 | LpACC | RdlPFC | 1.1903754 |
| siteMANNHEIM | 1 | 20 | 0.478 | 0.9732 | RpACC | RdlPFC | 1.4773468 |
| siteMANNHEIM | 1 | 20 | 0.478 | 0.9732 | RsgACC | RdlPFC | 0.2582688 |
| siteMANNHEIM | 1 | 20 | 0.478 | 0.9732 | LdlPFC | RdlPFC | 0.602945 |
| siteMANNHEIM | 1 | 20 | 0.478 | 0.9732 | LNacc | mPFC | 0.1259911 |
| siteMANNHEIM | 1 | 20 | 0.478 | 0.9732 | RsgACC | mPFC | -0.0924502 |
| siteMANNHEIM | 1 | 20 | 0.478 | 0.9732 | LdlPFC | mPFC | 1.9909636 |
| siteMANNHEIM | 1 | 20 | 0.478 | 0.9732 | RdlPFC | mPFC | 0.583755 |
| siteMANNHEIM | 1 | 20 | 0.478 | 0.9732 | RpACC | vmPFC | -0.2264936 |
| siteMANNHEIM | 1 | 20 | 0.478 | 0.9732 | RsgACC | vmPFC | -0.1189089 |
| siteMANNHEIM | 1 | 20 | 0.478 | 0.9732 | LdlPFC | vmPFC | 1.1902853 |
| siteMANNHEIM | 1 | 20 | 0.478 | 0.9732 | mPFC | vmPFC | -0.0720622 |
| siteNOTTINGHAM | 1 | 1 | 1 | 1 | LNacc | RNacc | 1.3202052 |
| siteNOTTINGHAM | 1 | 1 | 1 | 1 | LpACC | RpACC | 0.0485222 |
| siteNOTTINGHAM | 1 | 1 | 1 | 1 | LsgACC | RsgACC | 3.4079872 |
| siteNOTTINGHAM | 1 | 1 | 1 | 1 | LpACC | RdlPFC | -0.0713545 |
| siteNOTTINGHAM | 1 | 1 | 1 | 1 | LpACC | mPFC | 0.789037 |
| siteNOTTINGHAM | 1 | 1 | 1 | 1 | RdlPFC | vmPFC | -0.3228984 |
| sitePARIS | 1 | 17 | 0.0004 | 0.0036 | LNacc | RNacc | -1.498735 |
| sitePARIS | 1 | 17 | 0.0004 | 0.0036 | RNacc | LsgACC | -0.036768 |
| sitePARIS | 1 | 17 | 0.0004 | 0.0036 | LpACC | LsgACC | -0.2745787 |
| sitePARIS | 1 | 17 | 0.0004 | 0.0036 | RpACC | LsgACC | -0.3916982 |
| sitePARIS | 1 | 17 | 0.0004 | 0.0036 | LNacc | RsgACC | -0.4950952 |
| sitePARIS | 1 | 17 | 0.0004 | 0.0036 | RNacc | RsgACC | -0.5162655 |
| sitePARIS | 1 | 17 | 0.0004 | 0.0036 | LpACC | RsgACC | -0.8523144 |
| sitePARIS | 1 | 17 | 0.0004 | 0.0036 | RpACC | RsgACC | -0.8352132 |
| sitePARIS | 1 | 17 | 0.0004 | 0.0036 | LpACC | LdlPFC | 1.1543635 |
| sitePARIS | 1 | 17 | 0.0004 | 0.0036 | RpACC | LdlPFC | 0.9097483 |
| sitePARIS | 1 | 17 | 0.0004 | 0.0036 | LdlPFC | RdlPFC | 0.6518721 |
| sitePARIS | 1 | 17 | 0.0004 | 0.0036 | LsgACC | mPFC | -0.7168051 |
| sitePARIS | 1 | 17 | 0.0004 | 0.0036 | RsgACC | mPFC | -0.6339171 |
| sitePARIS | 1 | 17 | 0.0004 | 0.0036 | LdlPFC | mPFC | 0.434521 |
| sitePARIS | 1 | 17 | 0.0004 | 0.0036 | LpACC | vmPFC | -0.2000881 |
| sitePARIS | 1 | 17 | 0.0004 | 0.0036 | RpACC | vmPFC | -0.0827885 |
| sitePARIS | 1 | 17 | 0.0004 | 0.0036 | LsgACC | vmPFC | -0.0440047 |
| timepointFU3:threat_z | 1 | 17 | 0.0082 | 0.015 | LNacc | LpACC | -0.27255 |
| timepointFU3:threat_z | 1 | 17 | 0.0082 | 0.015 | RNacc | LpACC | -0.7347037 |
| timepointFU3:threat_z | 1 | 17 | 0.0082 | 0.015 | LNacc | RpACC | -0.7325871 |
| timepointFU3:threat_z | 1 | 17 | 0.0082 | 0.015 | RNacc | RpACC | -1.0481856 |
| timepointFU3:threat_z | 1 | 17 | 0.0082 | 0.015 | LNacc | LsgACC | -1.2337394 |
| timepointFU3:threat_z | 1 | 17 | 0.0082 | 0.015 | RNacc | LsgACC | -0.8887246 |
| timepointFU3:threat_z | 1 | 17 | 0.0082 | 0.015 | LNacc | RsgACC | -0.0729703 |
| timepointFU3:threat_z | 1 | 17 | 0.0082 | 0.015 | LNacc | RdlPFC | -0.7445791 |
| timepointFU3:threat_z | 1 | 17 | 0.0082 | 0.015 | RNacc | RdlPFC | -0.5785318 |
| timepointFU3:threat_z | 1 | 17 | 0.0082 | 0.015 | RpACC | RdlPFC | -0.3306245 |
| timepointFU3:threat_z | 1 | 17 | 0.0082 | 0.015 | RNacc | mPFC | -0.1836329 |
| timepointFU3:threat_z | 1 | 17 | 0.0082 | 0.015 | LsgACC | mPFC | -0.5199616 |
| timepointFU3:threat_z | 1 | 17 | 0.0082 | 0.015 | RsgACC | mPFC | -0.5599196 |
| timepointFU3:threat_z | 1 | 17 | 0.0082 | 0.015 | RdlPFC | mPFC | -0.1071426 |
| timepointFU3:threat_z | 1 | 17 | 0.0082 | 0.015 | RNacc | vmPFC | -0.9641077 |
| timepointFU3:threat_z | 1 | 17 | 0.0082 | 0.015 | RdlPFC | vmPFC | -0.1914931 |
| timepointFU3:threat_z | 1 | 17 | 0.0082 | 0.015 | mPFC | vmPFC | -0.139362 |
| timepointFU3:deprivation_z | 1 | 14 | 0.009 | 0.0166 | LNacc | LpACC | 0.0068072 |
| timepointFU3:deprivation_z | 1 | 14 | 0.009 | 0.0166 | LNacc | RpACC | 0.3135676 |
| timepointFU3:deprivation_z | 1 | 14 | 0.009 | 0.0166 | LNacc | LsgACC | 0.1884631 |
| timepointFU3:deprivation_z | 1 | 14 | 0.009 | 0.0166 | LpACC | LsgACC | 0.7726515 |
| timepointFU3:deprivation_z | 1 | 14 | 0.009 | 0.0166 | RpACC | LsgACC | 0.2830365 |
| timepointFU3:deprivation_z | 1 | 14 | 0.009 | 0.0166 | RpACC | RsgACC | 0.1091638 |
| timepointFU3:deprivation_z | 1 | 14 | 0.009 | 0.0166 | LNacc | RdlPFC | 0.6737109 |
| timepointFU3:deprivation_z | 1 | 14 | 0.009 | 0.0166 | RNacc | RdlPFC | 0.0650219 |
| timepointFU3:deprivation_z | 1 | 14 | 0.009 | 0.0166 | RpACC | RdlPFC | 0.4467336 |
| timepointFU3:deprivation_z | 1 | 14 | 0.009 | 0.0166 | LsgACC | RdlPFC | 0.5670275 |
| timepointFU3:deprivation_z | 1 | 14 | 0.009 | 0.0166 | RsgACC | RdlPFC | 0.0839457 |
| timepointFU3:deprivation_z | 1 | 14 | 0.009 | 0.0166 | LsgACC | mPFC | 1.0167739 |
| timepointFU3:deprivation_z | 1 | 14 | 0.009 | 0.0166 | RsgACC | mPFC | 1.0386782 |
| timepointFU3:deprivation_z | 1 | 14 | 0.009 | 0.0166 | RdlPFC | vmPFC | 0.2748434 |

**Sensitivity analysis: Testing robustness of interaction clusters to different cluster-defining thresholds**

**Table S7:** *Cross sectional model (CDT: p<0.01)*

| Variable | Cluster | No. Edges | p-FWE (Size) | p-FWE (Strength) | ROI1 | ROI2 | Strength |
| --- | --- | --- | --- | --- | --- | --- | --- |
| SexMale | 1 | 24 | 0.0004 | 0.0004 | LNacc | LpACC | 0.8909454 |
| SexMale | 1 | 24 | 0.0004 | 0.0004 | RNacc | LpACC | 0.0713489 |
| SexMale | 1 | 24 | 0.0004 | 0.0004 | LNacc | RpACC | 0.7316827 |
| SexMale | 1 | 24 | 0.0004 | 0.0004 | RNacc | RpACC | 0.7643312 |
| SexMale | 1 | 24 | 0.0004 | 0.0004 | LpACC | RpACC | 1.6113778 |
| SexMale | 1 | 24 | 0.0004 | 0.0004 | RNacc | LsgACC | 0.0314632 |
| SexMale | 1 | 24 | 0.0004 | 0.0004 | LsgACC | RsgACC | 0.0191965 |
| SexMale | 1 | 24 | 0.0004 | 0.0004 | LNacc | LdlPFC | 1.9742745 |
| SexMale | 1 | 24 | 0.0004 | 0.0004 | RNacc | LdlPFC | 1.0494837 |
| SexMale | 1 | 24 | 0.0004 | 0.0004 | LpACC | LdlPFC | 1.0345007 |
| SexMale | 1 | 24 | 0.0004 | 0.0004 | RpACC | LdlPFC | 1.3339215 |
| SexMale | 1 | 24 | 0.0004 | 0.0004 | LNacc | RdlPFC | 1.8856397 |
| SexMale | 1 | 24 | 0.0004 | 0.0004 | RNacc | RdlPFC | 1.3800857 |
| SexMale | 1 | 24 | 0.0004 | 0.0004 | LpACC | RdlPFC | 0.1597143 |
| SexMale | 1 | 24 | 0.0004 | 0.0004 | LdlPFC | RdlPFC | 0.130928 |
| SexMale | 1 | 24 | 0.0004 | 0.0004 | LNacc | mPFC | 0.5215232 |
| SexMale | 1 | 24 | 0.0004 | 0.0004 | LdlPFC | mPFC | 0.254153 |
| SexMale | 1 | 24 | 0.0004 | 0.0004 | LNacc | vmPFC | 2.4720162 |
| SexMale | 1 | 24 | 0.0004 | 0.0004 | RNacc | vmPFC | 0.9195171 |
| SexMale | 1 | 24 | 0.0004 | 0.0004 | LpACC | vmPFC | 0.2418974 |
| SexMale | 1 | 24 | 0.0004 | 0.0004 | RpACC | vmPFC | 0.3347258 |
| SexMale | 1 | 24 | 0.0004 | 0.0004 | LsgACC | vmPFC | 0.7509161 |
| SexMale | 1 | 24 | 0.0004 | 0.0004 | RsgACC | vmPFC | 0.3721589 |
| SexMale | 1 | 24 | 0.0004 | 0.0004 | LdlPFC | vmPFC | 0.3614311 |
| siteDRESDEN | 3 | 2 | 0.0878 | 0.0298 | LpACC | vmPFC | -0.8296557 |
| siteDRESDEN | 3 | 2 | 0.0878 | 0.0298 | RpACC | vmPFC | -0.4037293 |
| siteDUBLIN | 1 | 1 | 0.1934 | 0.0084 | LNacc | RNacc | 2.7544396 |
| siteDUBLIN | 1 | 1 | 0.1934 | 0.0084 | LpACC | RpACC | 1.2706354 |
| siteDUBLIN | 1 | 1 | 0.1934 | 0.0084 | LpACC | LsgACC | 0.6358059 |
| siteDUBLIN | 1 | 1 | 0.1934 | 0.0084 | RpACC | LsgACC | 0.5947326 |
| siteDUBLIN | 1 | 1 | 0.1934 | 0.0084 | LpACC | RsgACC | 0.631241 |
| siteDUBLIN | 1 | 1 | 0.1934 | 0.0084 | RpACC | RsgACC | 1.0795108 |
| siteDUBLIN | 1 | 1 | 0.1934 | 0.0084 | LsgACC | RsgACC | 0.097047 |
| siteDUBLIN | 1 | 1 | 0.1934 | 0.0084 | LdlPFC | RdlPFC | 2.0540642 |
| siteDUBLIN | 1 | 1 | 0.1934 | 0.0084 | LpACC | mPFC | 2.0151585 |
| siteDUBLIN | 1 | 1 | 0.1934 | 0.0084 | RpACC | mPFC | 1.0679419 |
| siteDUBLIN | 1 | 1 | 0.1934 | 0.0084 | LsgACC | mPFC | 1.0950706 |
| siteDUBLIN | 1 | 1 | 0.1934 | 0.0084 | RsgACC | mPFC | 1.1007628 |
| siteDUBLIN | 1 | 1 | 0.1934 | 0.0084 | mPFC | vmPFC | 0.4073975 |
| siteHAMBURG | 4 | 4 | 0.0272 | 0.0088 | LsgACC | RsgACC | 0.43887 |
| siteHAMBURG | 4 | 4 | 0.0272 | 0.0088 | RpACC | LdlPFC | 0.3161723 |
| siteHAMBURG | 4 | 4 | 0.0272 | 0.0088 | RpACC | RdlPFC | 0.5458176 |
| siteHAMBURG | 4 | 4 | 0.0272 | 0.0088 | LdlPFC | RdlPFC | 1.2845906 |
| siteHAMBURG | 4 | 4 | 0.0272 | 0.0088 | LdlPFC | mPFC | 0.7029389 |
| siteLONDON | 1 | 36 | 0 | 0 | LNacc | RNacc | 6.638418 |
| siteLONDON | 1 | 36 | 0 | 0 | LNacc | LpACC | 5.2399808 |
| siteLONDON | 1 | 36 | 0 | 0 | RNacc | LpACC | 2.8514117 |
| siteLONDON | 1 | 36 | 0 | 0 | LNacc | RpACC | 4.8807385 |
| siteLONDON | 1 | 36 | 0 | 0 | RNacc | RpACC | 3.6746897 |
| siteLONDON | 1 | 36 | 0 | 0 | LpACC | RpACC | 4.7590118 |
| siteLONDON | 1 | 36 | 0 | 0 | LNacc | LsgACC | 0.3556705 |
| siteLONDON | 1 | 36 | 0 | 0 | RNacc | LsgACC | 0.977284 |
| siteLONDON | 1 | 36 | 0 | 0 | LNacc | RsgACC | 1.9498268 |
| siteLONDON | 1 | 36 | 0 | 0 | RNacc | RsgACC | 2.0201172 |
| siteLONDON | 1 | 36 | 0 | 0 | LsgACC | RsgACC | 7.6056718 |
| siteLONDON | 1 | 36 | 0 | 0 | LNacc | LdlPFC | 2.2455516 |
| siteLONDON | 1 | 36 | 0 | 0 | RNacc | LdlPFC | 0.0488956 |
| siteLONDON | 1 | 36 | 0 | 0 | LpACC | LdlPFC | 1.4590664 |
| siteLONDON | 1 | 36 | 0 | 0 | RpACC | LdlPFC | 1.4870962 |
| siteLONDON | 1 | 36 | 0 | 0 | LsgACC | LdlPFC | 2.310467 |
| siteLONDON | 1 | 36 | 0 | 0 | RsgACC | LdlPFC | 3.3698546 |
| siteLONDON | 1 | 36 | 0 | 0 | LNacc | RdlPFC | 2.3317994 |
| siteLONDON | 1 | 36 | 0 | 0 | RNacc | RdlPFC | 1.3078213 |
| siteLONDON | 1 | 36 | 0 | 0 | LpACC | RdlPFC | 0.9334938 |
| siteLONDON | 1 | 36 | 0 | 0 | RpACC | RdlPFC | 1.708201 |
| siteLONDON | 1 | 36 | 0 | 0 | LsgACC | RdlPFC | 2.4138165 |
| siteLONDON | 1 | 36 | 0 | 0 | RsgACC | RdlPFC | 2.2310322 |
| siteLONDON | 1 | 36 | 0 | 0 | LdlPFC | RdlPFC | 3.7292227 |
| siteLONDON | 1 | 36 | 0 | 0 | LNacc | mPFC | 3.7204812 |
| siteLONDON | 1 | 36 | 0 | 0 | RNacc | mPFC | 2.2831107 |
| siteLONDON | 1 | 36 | 0 | 0 | LpACC | mPFC | 3.6950889 |
| siteLONDON | 1 | 36 | 0 | 0 | RpACC | mPFC | 1.9138876 |
| siteLONDON | 1 | 36 | 0 | 0 | LdlPFC | mPFC | 1.6989993 |
| siteLONDON | 1 | 36 | 0 | 0 | RdlPFC | mPFC | 1.2799579 |
| siteLONDON | 1 | 36 | 0 | 0 | LNacc | vmPFC | 2.5734979 |
| siteLONDON | 1 | 36 | 0 | 0 | RNacc | vmPFC | 2.0104234 |
| siteLONDON | 1 | 36 | 0 | 0 | LsgACC | vmPFC | 0.5922976 |
| siteLONDON | 1 | 36 | 0 | 0 | RsgACC | vmPFC | 1.2145913 |
| siteLONDON | 1 | 36 | 0 | 0 | LdlPFC | vmPFC | 0.3994763 |
| siteLONDON | 1 | 36 | 0 | 0 | RdlPFC | vmPFC | 0.162995 |
| siteMANNHEIM | 1 | 18 | 0.0002 | 0 | LNacc | LpACC | 0.2016985 |
| siteMANNHEIM | 1 | 18 | 0.0002 | 0 | RNacc | LpACC | 0.0755146 |
| siteMANNHEIM | 1 | 18 | 0.0002 | 0 | LNacc | RpACC | 0.3321766 |
| siteMANNHEIM | 1 | 18 | 0.0002 | 0 | RNacc | RpACC | 0.5643059 |
| siteMANNHEIM | 1 | 18 | 0.0002 | 0 | LpACC | LdlPFC | 0.6541813 |
| siteMANNHEIM | 1 | 18 | 0.0002 | 0 | RpACC | LdlPFC | 1.5411596 |
| siteMANNHEIM | 1 | 18 | 0.0002 | 0 | RsgACC | LdlPFC | 1.3497371 |
| siteMANNHEIM | 1 | 18 | 0.0002 | 0 | RNacc | RdlPFC | 0.423318 |
| siteMANNHEIM | 1 | 18 | 0.0002 | 0 | LpACC | RdlPFC | 0.3273481 |
| siteMANNHEIM | 1 | 18 | 0.0002 | 0 | RpACC | RdlPFC | 1.9531186 |
| siteMANNHEIM | 1 | 18 | 0.0002 | 0 | LsgACC | RdlPFC | 0.0742413 |
| siteMANNHEIM | 1 | 18 | 0.0002 | 0 | RsgACC | RdlPFC | 0.6636093 |
| siteMANNHEIM | 1 | 18 | 0.0002 | 0 | LdlPFC | RdlPFC | 1.5230897 |
| siteMANNHEIM | 1 | 18 | 0.0002 | 0 | RNacc | mPFC | 0.2513132 |
| siteMANNHEIM | 1 | 18 | 0.0002 | 0 | LdlPFC | mPFC | 1.4450451 |
| siteMANNHEIM | 1 | 18 | 0.0002 | 0 | RdlPFC | mPFC | 0.4657511 |
| siteMANNHEIM | 1 | 18 | 0.0002 | 0 | LdlPFC | vmPFC | 1.2969026 |
| siteMANNHEIM | 1 | 18 | 0.0002 | 0 | RdlPFC | vmPFC | 1.1375645 |
| siteNOTTINGHAM | 1 | 1 | 0.2088 | 0.0278 | LNacc | RNacc | 1.3118533 |
| siteNOTTINGHAM | 1 | 1 | 0.2088 | 0.0278 | LpACC | RpACC | 1.5665879 |
| siteNOTTINGHAM | 1 | 1 | 0.2088 | 0.0278 | LpACC | LsgACC | 1.2594878 |
| siteNOTTINGHAM | 1 | 1 | 0.2088 | 0.0278 | RpACC | LsgACC | 1.1031939 |
| siteNOTTINGHAM | 1 | 1 | 0.2088 | 0.0278 | LpACC | RsgACC | 1.3829583 |
| siteNOTTINGHAM | 1 | 1 | 0.2088 | 0.0278 | RpACC | RsgACC | 1.2806425 |
| siteNOTTINGHAM | 1 | 1 | 0.2088 | 0.0278 | LsgACC | RsgACC | 1.9029372 |
| siteNOTTINGHAM | 1 | 1 | 0.2088 | 0.0278 | LpACC | mPFC | 1.4646872 |
| siteNOTTINGHAM | 1 | 1 | 0.2088 | 0.0278 | RpACC | mPFC | 1.0556126 |
| siteNOTTINGHAM | 1 | 1 | 0.2088 | 0.0278 | LsgACC | mPFC | 0.4361876 |
| siteNOTTINGHAM | 1 | 1 | 0.2088 | 0.0278 | RsgACC | mPFC | 0.6400227 |
| siteNOTTINGHAM | 1 | 1 | 0.2088 | 0.0278 | LsgACC | vmPFC | 0.5724328 |
| siteNOTTINGHAM | 1 | 1 | 0.2088 | 0.0278 | RsgACC | vmPFC | 0.8356143 |
| sitePARIS | 3 | 6 | 0.0106 | 0.0164 | LpACC | LsgACC | -0.1607908 |
| sitePARIS | 3 | 6 | 0.0106 | 0.0164 | LpACC | RsgACC | -0.2892575 |
| sitePARIS | 3 | 6 | 0.0106 | 0.0164 | RpACC | RsgACC | -0.4387223 |
| sitePARIS | 3 | 6 | 0.0106 | 0.0164 | LsgACC | mPFC | -0.2848498 |
| sitePARIS | 3 | 6 | 0.0106 | 0.0164 | RsgACC | mPFC | -0.5475235 |
| sitePARIS | 3 | 6 | 0.0106 | 0.0164 | LpACC | vmPFC | -0.0183864 |
| mean_fd | 1 | 44 | 0 | 0 | LNacc | RNacc | 10.223781 |
| mean_fd | 1 | 44 | 0 | 0 | LNacc | LpACC | 5.0086435 |
| mean_fd | 1 | 44 | 0 | 0 | RNacc | LpACC | 5.8047481 |
| mean_fd | 1 | 44 | 0 | 0 | LNacc | RpACC | 5.7842353 |
| mean_fd | 1 | 44 | 0 | 0 | RNacc | RpACC | 5.3647254 |
| mean_fd | 1 | 44 | 0 | 0 | LpACC | RpACC | 5.9881646 |
| mean_fd | 1 | 44 | 0 | 0 | LNacc | LsgACC | 8.0139001 |
| mean_fd | 1 | 44 | 0 | 0 | RNacc | LsgACC | 7.2894585 |
| mean_fd | 1 | 44 | 0 | 0 | LpACC | LsgACC | 2.1929923 |
| mean_fd | 1 | 44 | 0 | 0 | RpACC | LsgACC | 2.6276086 |
| mean_fd | 1 | 44 | 0 | 0 | LNacc | RsgACC | 8.4425207 |
| mean_fd | 1 | 44 | 0 | 0 | RNacc | RsgACC | 7.2590931 |
| mean_fd | 1 | 44 | 0 | 0 | LpACC | RsgACC | 2.5650119 |
| mean_fd | 1 | 44 | 0 | 0 | RpACC | RsgACC | 2.0994298 |
| mean_fd | 1 | 44 | 0 | 0 | LsgACC | RsgACC | 6.5771021 |
| mean_fd | 1 | 44 | 0 | 0 | LNacc | LdlPFC | 4.494307 |
| mean_fd | 1 | 44 | 0 | 0 | RNacc | LdlPFC | 4.0549306 |
| mean_fd | 1 | 44 | 0 | 0 | LpACC | LdlPFC | 3.9748904 |
| mean_fd | 1 | 44 | 0 | 0 | RpACC | LdlPFC | 3.1288231 |
| mean_fd | 1 | 44 | 0 | 0 | LsgACC | LdlPFC | 5.6021863 |
| mean_fd | 1 | 44 | 0 | 0 | RsgACC | LdlPFC | 6.5060552 |
| mean_fd | 1 | 44 | 0 | 0 | LNacc | RdlPFC | 4.668652 |
| mean_fd | 1 | 44 | 0 | 0 | RNacc | RdlPFC | 4.4799632 |
| mean_fd | 1 | 44 | 0 | 0 | LpACC | RdlPFC | 4.7014062 |
| mean_fd | 1 | 44 | 0 | 0 | RpACC | RdlPFC | 3.546856 |
| mean_fd | 1 | 44 | 0 | 0 | LsgACC | RdlPFC | 5.294147 |
| mean_fd | 1 | 44 | 0 | 0 | RsgACC | RdlPFC | 5.2046931 |
| mean_fd | 1 | 44 | 0 | 0 | LNacc | mPFC | 5.2207146 |
| mean_fd | 1 | 44 | 0 | 0 | RNacc | mPFC | 5.9079445 |
| mean_fd | 1 | 44 | 0 | 0 | LpACC | mPFC | 2.4237196 |
| mean_fd | 1 | 44 | 0 | 0 | RpACC | mPFC | 3.2989208 |
| mean_fd | 1 | 44 | 0 | 0 | LsgACC | mPFC | 1.3456485 |
| mean_fd | 1 | 44 | 0 | 0 | RsgACC | mPFC | 1.0733042 |
| mean_fd | 1 | 44 | 0 | 0 | LdlPFC | mPFC | 3.9654881 |
| mean_fd | 1 | 44 | 0 | 0 | RdlPFC | mPFC | 4.50784 |
| mean_fd | 1 | 44 | 0 | 0 | LNacc | vmPFC | 8.2131296 |
| mean_fd | 1 | 44 | 0 | 0 | RNacc | vmPFC | 8.0207433 |
| mean_fd | 1 | 44 | 0 | 0 | LpACC | vmPFC | 0.5435161 |
| mean_fd | 1 | 44 | 0 | 0 | RpACC | vmPFC | 0.8681445 |
| mean_fd | 1 | 44 | 0 | 0 | LsgACC | vmPFC | 4.7582264 |
| mean_fd | 1 | 44 | 0 | 0 | RsgACC | vmPFC | 3.2875741 |
| mean_fd | 1 | 44 | 0 | 0 | LdlPFC | vmPFC | 3.5271669 |
| mean_fd | 1 | 44 | 0 | 0 | RdlPFC | vmPFC | 4.5462846 |
| mean_fd | 1 | 44 | 0 | 0 | mPFC | vmPFC | 0.5499964 |
| SexMale:deprivation_z | 1 | 3 | 0.044 | 0.0316 | LNacc | RNacc | 0.2283612 |
| SexMale:deprivation_z | 1 | 3 | 0.044 | 0.0316 | RNacc | RdlPFC | 0.3937369 |
| SexMale:deprivation_z | 1 | 3 | 0.044 | 0.0316 | RdlPFC | vmPFC | 0.4902561 |

***Table S8:*** *Cross sectional model (CDT: p<0.005)*

| Variable | Cluster | No. Edges | p-FWE (Size) | p-FWE (Strength) | ROI1 | ROI2 | Strength |
| --- | --- | --- | --- | --- | --- | --- | --- |
| SexMale | 1 | 19 | 0 | 0 | LNacc | LpACC | 0.6575312 |
| SexMale | 1 | 19 | 0 | 0 | LNacc | RpACC | 0.4982686 |
| SexMale | 1 | 19 | 0 | 0 | RNacc | RpACC | 0.530917 |
| SexMale | 1 | 19 | 0 | 0 | LpACC | RpACC | 1.3779636 |
| SexMale | 1 | 19 | 0 | 0 | LNacc | LdlPFC | 1.7408604 |
| SexMale | 1 | 19 | 0 | 0 | RNacc | LdlPFC | 0.8160696 |
| SexMale | 1 | 19 | 0 | 0 | LpACC | LdlPFC | 0.8010865 |
| SexMale | 1 | 19 | 0 | 0 | RpACC | LdlPFC | 1.1005073 |
| SexMale | 1 | 19 | 0 | 0 | LNacc | RdlPFC | 1.6522255 |
| SexMale | 1 | 19 | 0 | 0 | RNacc | RdlPFC | 1.1466716 |
| SexMale | 1 | 19 | 0 | 0 | LNacc | mPFC | 0.288109 |
| SexMale | 1 | 19 | 0 | 0 | LdlPFC | mPFC | 0.0207388 |
| SexMale | 1 | 19 | 0 | 0 | LNacc | vmPFC | 2.238602 |
| SexMale | 1 | 19 | 0 | 0 | RNacc | vmPFC | 0.6861029 |
| SexMale | 1 | 19 | 0 | 0 | LpACC | vmPFC | 0.0084833 |
| SexMale | 1 | 19 | 0 | 0 | RpACC | vmPFC | 0.1013116 |
| SexMale | 1 | 19 | 0 | 0 | LsgACC | vmPFC | 0.5175019 |
| SexMale | 1 | 19 | 0 | 0 | RsgACC | vmPFC | 0.1387447 |
| SexMale | 1 | 19 | 0 | 0 | LdlPFC | vmPFC | 0.1280169 |
| siteDRESDEN | 3 | 2 | 0.044 | 0.0256 | LpACC | vmPFC | -0.5962415 |
| siteDRESDEN | 3 | 2 | 0.044 | 0.0256 | RpACC | vmPFC | -0.1703151 |
| siteDUBLIN | 1 | 1 | 0.1104 | 0.0046 | LNacc | RNacc | 2.5210255 |
| siteDUBLIN | 1 | 1 | 0.1104 | 0.0046 | LpACC | RpACC | 1.0372212 |
| siteDUBLIN | 1 | 1 | 0.1104 | 0.0046 | LpACC | LsgACC | 0.4023917 |
| siteDUBLIN | 1 | 1 | 0.1104 | 0.0046 | RpACC | LsgACC | 0.3613184 |
| siteDUBLIN | 1 | 1 | 0.1104 | 0.0046 | LpACC | RsgACC | 0.3978268 |
| siteDUBLIN | 1 | 1 | 0.1104 | 0.0046 | RpACC | RsgACC | 0.8460966 |
| siteDUBLIN | 1 | 1 | 0.1104 | 0.0046 | LdlPFC | RdlPFC | 1.82065 |
| siteDUBLIN | 1 | 1 | 0.1104 | 0.0046 | LpACC | mPFC | 1.7817443 |
| siteDUBLIN | 1 | 1 | 0.1104 | 0.0046 | RpACC | mPFC | 0.8345277 |
| siteDUBLIN | 1 | 1 | 0.1104 | 0.0046 | LsgACC | mPFC | 0.8616564 |
| siteDUBLIN | 1 | 1 | 0.1104 | 0.0046 | RsgACC | mPFC | 0.8673486 |
| siteDUBLIN | 1 | 1 | 0.1104 | 0.0046 | mPFC | vmPFC | 0.1739834 |
| siteHAMBURG | 4 | 4 | 0.015 | 0.008 | LsgACC | RsgACC | 0.2054558 |
| siteHAMBURG | 4 | 4 | 0.015 | 0.008 | RpACC | LdlPFC | 0.0827581 |
| siteHAMBURG | 4 | 4 | 0.015 | 0.008 | RpACC | RdlPFC | 0.3124034 |
| siteHAMBURG | 4 | 4 | 0.015 | 0.008 | LdlPFC | RdlPFC | 1.0511764 |
| siteHAMBURG | 4 | 4 | 0.015 | 0.008 | LdlPFC | mPFC | 0.4695247 |
| siteLONDON | 1 | 34 | 0 | 0 | LNacc | RNacc | 6.4050038 |
| siteLONDON | 1 | 34 | 0 | 0 | LNacc | LpACC | 5.0065666 |
| siteLONDON | 1 | 34 | 0 | 0 | RNacc | LpACC | 2.6179975 |
| siteLONDON | 1 | 34 | 0 | 0 | LNacc | RpACC | 4.6473243 |
| siteLONDON | 1 | 34 | 0 | 0 | RNacc | RpACC | 3.4412755 |
| siteLONDON | 1 | 34 | 0 | 0 | LpACC | RpACC | 4.5255976 |
| siteLONDON | 1 | 34 | 0 | 0 | LNacc | LsgACC | 0.1222563 |
| siteLONDON | 1 | 34 | 0 | 0 | RNacc | LsgACC | 0.7438698 |
| siteLONDON | 1 | 34 | 0 | 0 | LNacc | RsgACC | 1.7164126 |
| siteLONDON | 1 | 34 | 0 | 0 | RNacc | RsgACC | 1.786703 |
| siteLONDON | 1 | 34 | 0 | 0 | LsgACC | RsgACC | 7.3722576 |
| siteLONDON | 1 | 34 | 0 | 0 | LNacc | LdlPFC | 2.0121374 |
| siteLONDON | 1 | 34 | 0 | 0 | LpACC | LdlPFC | 1.2256522 |
| siteLONDON | 1 | 34 | 0 | 0 | RpACC | LdlPFC | 1.253682 |
| siteLONDON | 1 | 34 | 0 | 0 | LsgACC | LdlPFC | 2.0770528 |
| siteLONDON | 1 | 34 | 0 | 0 | RsgACC | LdlPFC | 3.1364404 |
| siteLONDON | 1 | 34 | 0 | 0 | LNacc | RdlPFC | 2.0983852 |
| siteLONDON | 1 | 34 | 0 | 0 | RNacc | RdlPFC | 1.0744072 |
| siteLONDON | 1 | 34 | 0 | 0 | LpACC | RdlPFC | 0.7000796 |
| siteLONDON | 1 | 34 | 0 | 0 | RpACC | RdlPFC | 1.4747868 |
| siteLONDON | 1 | 34 | 0 | 0 | LsgACC | RdlPFC | 2.1804023 |
| siteLONDON | 1 | 34 | 0 | 0 | RsgACC | RdlPFC | 1.997618 |
| siteLONDON | 1 | 34 | 0 | 0 | LdlPFC | RdlPFC | 3.4958085 |
| siteLONDON | 1 | 34 | 0 | 0 | LNacc | mPFC | 3.487067 |
| siteLONDON | 1 | 34 | 0 | 0 | RNacc | mPFC | 2.0496966 |
| siteLONDON | 1 | 34 | 0 | 0 | LpACC | mPFC | 3.4616747 |
| siteLONDON | 1 | 34 | 0 | 0 | RpACC | mPFC | 1.6804734 |
| siteLONDON | 1 | 34 | 0 | 0 | LdlPFC | mPFC | 1.4655851 |
| siteLONDON | 1 | 34 | 0 | 0 | RdlPFC | mPFC | 1.0465437 |
| siteLONDON | 1 | 34 | 0 | 0 | LNacc | vmPFC | 2.3400837 |
| siteLONDON | 1 | 34 | 0 | 0 | RNacc | vmPFC | 1.7770092 |
| siteLONDON | 1 | 34 | 0 | 0 | LsgACC | vmPFC | 0.3588834 |
| siteLONDON | 1 | 34 | 0 | 0 | RsgACC | vmPFC | 0.9811771 |
| siteLONDON | 1 | 34 | 0 | 0 | LdlPFC | vmPFC | 0.1660621 |
| siteMANNHEIM | 1 | 15 | 0 | 0 | LNacc | RpACC | 0.0987624 |
| siteMANNHEIM | 1 | 15 | 0 | 0 | RNacc | RpACC | 0.3308917 |
| siteMANNHEIM | 1 | 15 | 0 | 0 | LpACC | LdlPFC | 0.4207671 |
| siteMANNHEIM | 1 | 15 | 0 | 0 | RpACC | LdlPFC | 1.3077454 |
| siteMANNHEIM | 1 | 15 | 0 | 0 | RsgACC | LdlPFC | 1.1163229 |
| siteMANNHEIM | 1 | 15 | 0 | 0 | RNacc | RdlPFC | 0.1899038 |
| siteMANNHEIM | 1 | 15 | 0 | 0 | LpACC | RdlPFC | 0.0939339 |
| siteMANNHEIM | 1 | 15 | 0 | 0 | RpACC | RdlPFC | 1.7197044 |
| siteMANNHEIM | 1 | 15 | 0 | 0 | RsgACC | RdlPFC | 0.4301951 |
| siteMANNHEIM | 1 | 15 | 0 | 0 | LdlPFC | RdlPFC | 1.2896755 |
| siteMANNHEIM | 1 | 15 | 0 | 0 | RNacc | mPFC | 0.017899 |
| siteMANNHEIM | 1 | 15 | 0 | 0 | LdlPFC | mPFC | 1.211631 |
| siteMANNHEIM | 1 | 15 | 0 | 0 | RdlPFC | mPFC | 0.2323369 |
| siteMANNHEIM | 1 | 15 | 0 | 0 | LdlPFC | vmPFC | 1.0634884 |
| siteMANNHEIM | 1 | 15 | 0 | 0 | RdlPFC | vmPFC | 0.9041503 |
| siteNOTTINGHAM | 1 | 1 | 0.1132 | 0.019 | LNacc | RNacc | 1.0784391 |
| siteNOTTINGHAM | 1 | 1 | 0.1132 | 0.019 | LpACC | RpACC | 1.3331737 |
| siteNOTTINGHAM | 1 | 1 | 0.1132 | 0.019 | LpACC | LsgACC | 1.0260736 |
| siteNOTTINGHAM | 1 | 1 | 0.1132 | 0.019 | RpACC | LsgACC | 0.8697797 |
| siteNOTTINGHAM | 1 | 1 | 0.1132 | 0.019 | LpACC | RsgACC | 1.1495441 |
| siteNOTTINGHAM | 1 | 1 | 0.1132 | 0.019 | RpACC | RsgACC | 1.0472283 |
| siteNOTTINGHAM | 1 | 1 | 0.1132 | 0.019 | LsgACC | RsgACC | 1.669523 |
| siteNOTTINGHAM | 1 | 1 | 0.1132 | 0.019 | LpACC | mPFC | 1.231273 |
| siteNOTTINGHAM | 1 | 1 | 0.1132 | 0.019 | RpACC | mPFC | 0.8221984 |
| siteNOTTINGHAM | 1 | 1 | 0.1132 | 0.019 | LsgACC | mPFC | 0.2027735 |
| siteNOTTINGHAM | 1 | 1 | 0.1132 | 0.019 | RsgACC | mPFC | 0.4066085 |
| siteNOTTINGHAM | 1 | 1 | 0.1132 | 0.019 | LsgACC | vmPFC | 0.3390187 |
| siteNOTTINGHAM | 1 | 1 | 0.1132 | 0.019 | RsgACC | vmPFC | 0.6022001 |
| sitePARIS | 3 | 4 | 0.0106 | 0.0282 | LpACC | RsgACC | -0.0558434 |
| sitePARIS | 3 | 4 | 0.0106 | 0.0282 | RpACC | RsgACC | -0.2053081 |
| sitePARIS | 3 | 4 | 0.0106 | 0.0282 | LsgACC | mPFC | -0.0514356 |
| sitePARIS | 3 | 4 | 0.0106 | 0.0282 | RsgACC | mPFC | -0.3141093 |
| mean_fd | 1 | 44 | 0 | 0 | LNacc | RNacc | 9.9903671 |
| mean_fd | 1 | 44 | 0 | 0 | LNacc | LpACC | 4.7752293 |
| mean_fd | 1 | 44 | 0 | 0 | RNacc | LpACC | 5.5713339 |
| mean_fd | 1 | 44 | 0 | 0 | LNacc | RpACC | 5.5508211 |
| mean_fd | 1 | 44 | 0 | 0 | RNacc | RpACC | 5.1313112 |
| mean_fd | 1 | 44 | 0 | 0 | LpACC | RpACC | 5.7547504 |
| mean_fd | 1 | 44 | 0 | 0 | LNacc | LsgACC | 7.7804859 |
| mean_fd | 1 | 44 | 0 | 0 | RNacc | LsgACC | 7.0560443 |
| mean_fd | 1 | 44 | 0 | 0 | LpACC | LsgACC | 1.9595781 |
| mean_fd | 1 | 44 | 0 | 0 | RpACC | LsgACC | 2.3941944 |
| mean_fd | 1 | 44 | 0 | 0 | LNacc | RsgACC | 8.2091065 |
| mean_fd | 1 | 44 | 0 | 0 | RNacc | RsgACC | 7.0256789 |
| mean_fd | 1 | 44 | 0 | 0 | LpACC | RsgACC | 2.3315977 |
| mean_fd | 1 | 44 | 0 | 0 | RpACC | RsgACC | 1.8660157 |
| mean_fd | 1 | 44 | 0 | 0 | LsgACC | RsgACC | 6.3436879 |
| mean_fd | 1 | 44 | 0 | 0 | LNacc | LdlPFC | 4.2608928 |
| mean_fd | 1 | 44 | 0 | 0 | RNacc | LdlPFC | 3.8215164 |
| mean_fd | 1 | 44 | 0 | 0 | LpACC | LdlPFC | 3.7414762 |
| mean_fd | 1 | 44 | 0 | 0 | RpACC | LdlPFC | 2.8954089 |
| mean_fd | 1 | 44 | 0 | 0 | LsgACC | LdlPFC | 5.3687721 |
| mean_fd | 1 | 44 | 0 | 0 | RsgACC | LdlPFC | 6.272641 |
| mean_fd | 1 | 44 | 0 | 0 | LNacc | RdlPFC | 4.4352378 |
| mean_fd | 1 | 44 | 0 | 0 | RNacc | RdlPFC | 4.246549 |
| mean_fd | 1 | 44 | 0 | 0 | LpACC | RdlPFC | 4.4679921 |
| mean_fd | 1 | 44 | 0 | 0 | RpACC | RdlPFC | 3.3134418 |
| mean_fd | 1 | 44 | 0 | 0 | LsgACC | RdlPFC | 5.0607328 |
| mean_fd | 1 | 44 | 0 | 0 | RsgACC | RdlPFC | 4.971279 |
| mean_fd | 1 | 44 | 0 | 0 | LNacc | mPFC | 4.9873004 |
| mean_fd | 1 | 44 | 0 | 0 | RNacc | mPFC | 5.6745303 |
| mean_fd | 1 | 44 | 0 | 0 | LpACC | mPFC | 2.1903054 |
| mean_fd | 1 | 44 | 0 | 0 | RpACC | mPFC | 3.0655066 |
| mean_fd | 1 | 44 | 0 | 0 | LsgACC | mPFC | 1.1122343 |
| mean_fd | 1 | 44 | 0 | 0 | RsgACC | mPFC | 0.83989 |
| mean_fd | 1 | 44 | 0 | 0 | LdlPFC | mPFC | 3.7320739 |
| mean_fd | 1 | 44 | 0 | 0 | RdlPFC | mPFC | 4.2744258 |
| mean_fd | 1 | 44 | 0 | 0 | LNacc | vmPFC | 7.9797154 |
| mean_fd | 1 | 44 | 0 | 0 | RNacc | vmPFC | 7.7873291 |
| mean_fd | 1 | 44 | 0 | 0 | LpACC | vmPFC | 0.3101019 |
| mean_fd | 1 | 44 | 0 | 0 | RpACC | vmPFC | 0.6347303 |
| mean_fd | 1 | 44 | 0 | 0 | LsgACC | vmPFC | 4.5248122 |
| mean_fd | 1 | 44 | 0 | 0 | RsgACC | vmPFC | 3.0541599 |
| mean_fd | 1 | 44 | 0 | 0 | LdlPFC | vmPFC | 3.2937528 |
| mean_fd | 1 | 44 | 0 | 0 | RdlPFC | vmPFC | 4.3128704 |
| mean_fd | 1 | 44 | 0 | 0 | mPFC | vmPFC | 0.3165822 |
| SexMale:deprivation_z | 2 | 2 | 0.0416 | 0.042 | RNacc | RdlPFC | 0.1603227 |
| SexMale:deprivation_z | 2 | 2 | 0.0416 | 0.042 | RdlPFC | vmPFC | 0.2568419 |

***Table S9:*** *Longitudinal Male model (CDT: p<0.01)*

| Variable | Cluster | No. Edges | p-FWE (Size) | p-FWE (Strength) | ROI1 | ROI2 | Strength |
| --- | --- | --- | --- | --- | --- | --- | --- |
| timepointFU3 | 1 | 8 | 0.004 | 0 | LNacc | RNacc | -1.8757483 |
| timepointFU3 | 1 | 8 | 0.004 | 0 | RNacc | LpACC | -0.1996905 |
| timepointFU3 | 1 | 8 | 0.004 | 0 | LNacc | RpACC | -0.0752934 |
| timepointFU3 | 1 | 8 | 0.004 | 0 | RNacc | RpACC | -0.6086607 |
| timepointFU3 | 1 | 8 | 0.004 | 0 | LNacc | RsgACC | -1.8058345 |
| timepointFU3 | 1 | 8 | 0.004 | 0 | RNacc | RsgACC | -1.9274411 |
| timepointFU3 | 1 | 8 | 0.004 | 0 | LsgACC | RsgACC | -3.309367 |
| timepointFU3 | 1 | 8 | 0.004 | 0 | RNacc | vmPFC | -0.1224513 |
| mean_fd | 1 | 42 | 0.014 | 0 | LNacc | RNacc | 6.0110311 |
| mean_fd | 1 | 42 | 0.014 | 0 | LNacc | LpACC | 3.3731916 |
| mean_fd | 1 | 42 | 0.014 | 0 | RNacc | LpACC | 2.6365445 |
| mean_fd | 1 | 42 | 0.014 | 0 | LNacc | RpACC | 4.0208352 |
| mean_fd | 1 | 42 | 0.014 | 0 | RNacc | RpACC | 3.6773075 |
| mean_fd | 1 | 42 | 0.014 | 0 | LpACC | RpACC | 3.8652684 |
| mean_fd | 1 | 42 | 0.014 | 0 | LNacc | LsgACC | 4.1341516 |
| mean_fd | 1 | 42 | 0.014 | 0 | RNacc | LsgACC | 4.1362622 |
| mean_fd | 1 | 42 | 0.014 | 0 | LpACC | LsgACC | 0.5595478 |
| mean_fd | 1 | 42 | 0.014 | 0 | RpACC | LsgACC | 2.1636324 |
| mean_fd | 1 | 42 | 0.014 | 0 | LNacc | RsgACC | 4.6741795 |
| mean_fd | 1 | 42 | 0.014 | 0 | RNacc | RsgACC | 4.4005857 |
| mean_fd | 1 | 42 | 0.014 | 0 | LpACC | RsgACC | 1.0551781 |
| mean_fd | 1 | 42 | 0.014 | 0 | RpACC | RsgACC | 1.7472756 |
| mean_fd | 1 | 42 | 0.014 | 0 | LsgACC | RsgACC | 2.5005706 |
| mean_fd | 1 | 42 | 0.014 | 0 | LNacc | LdlPFC | 0.1688978 |
| mean_fd | 1 | 42 | 0.014 | 0 | RNacc | LdlPFC | 0.8130803 |
| mean_fd | 1 | 42 | 0.014 | 0 | LpACC | LdlPFC | 0.9680004 |
| mean_fd | 1 | 42 | 0.014 | 0 | RpACC | LdlPFC | 0.4036871 |
| mean_fd | 1 | 42 | 0.014 | 0 | LsgACC | LdlPFC | 1.1229432 |
| mean_fd | 1 | 42 | 0.014 | 0 | RsgACC | LdlPFC | 0.6061163 |
| mean_fd | 1 | 42 | 0.014 | 0 | LNacc | RdlPFC | 0.7314622 |
| mean_fd | 1 | 42 | 0.014 | 0 | LpACC | RdlPFC | 1.8847396 |
| mean_fd | 1 | 42 | 0.014 | 0 | RpACC | RdlPFC | 0.6647794 |
| mean_fd | 1 | 42 | 0.014 | 0 | LsgACC | RdlPFC | 0.601511 |
| mean_fd | 1 | 42 | 0.014 | 0 | RsgACC | RdlPFC | 0.4599038 |
| mean_fd | 1 | 42 | 0.014 | 0 | LNacc | mPFC | 2.1891215 |
| mean_fd | 1 | 42 | 0.014 | 0 | RNacc | mPFC | 1.416752 |
| mean_fd | 1 | 42 | 0.014 | 0 | RpACC | mPFC | 0.45792 |
| mean_fd | 1 | 42 | 0.014 | 0 | LsgACC | mPFC | 0.5680835 |
| mean_fd | 1 | 42 | 0.014 | 0 | RsgACC | mPFC | 0.0765961 |
| mean_fd | 1 | 42 | 0.014 | 0 | LdlPFC | mPFC | 1.0496806 |
| mean_fd | 1 | 42 | 0.014 | 0 | RdlPFC | mPFC | 1.8887848 |
| mean_fd | 1 | 42 | 0.014 | 0 | LNacc | vmPFC | 4.6330291 |
| mean_fd | 1 | 42 | 0.014 | 0 | RNacc | vmPFC | 4.4144907 |
| mean_fd | 1 | 42 | 0.014 | 0 | LpACC | vmPFC | 0.5210701 |
| mean_fd | 1 | 42 | 0.014 | 0 | RpACC | vmPFC | 1.4752763 |
| mean_fd | 1 | 42 | 0.014 | 0 | LsgACC | vmPFC | 2.8294185 |
| mean_fd | 1 | 42 | 0.014 | 0 | RsgACC | vmPFC | 2.0624571 |
| mean_fd | 1 | 42 | 0.014 | 0 | LdlPFC | vmPFC | 1.0135434 |
| mean_fd | 1 | 42 | 0.014 | 0 | RdlPFC | vmPFC | 1.0276256 |
| mean_fd | 1 | 42 | 0.014 | 0 | mPFC | vmPFC | 0.1325927 |
| siteDUBLIN | 1 | 1 | 1 | 1 | LNacc | RNacc | 0.3071611 |
| siteDUBLIN | 1 | 1 | 1 | 1 | LpACC | RpACC | 1.0839278 |
| siteDUBLIN | 1 | 1 | 1 | 1 | LdlPFC | RdlPFC | 0.1818218 |
| siteDUBLIN | 1 | 1 | 1 | 1 | LpACC | mPFC | 1.6811967 |
| siteDUBLIN | 1 | 1 | 1 | 1 | RpACC | mPFC | 0.5516485 |
| siteDUBLIN | 1 | 1 | 1 | 1 | LpACC | vmPFC | 0.4756129 |
| siteDUBLIN | 1 | 1 | 1 | 1 | RpACC | vmPFC | 1.0868617 |
| siteDUBLIN | 1 | 1 | 1 | 1 | mPFC | vmPFC | 0.3616949 |
| siteHAMBURG | 4 | 2 | 1 | 0.534 | RpACC | RdlPFC | 0.0835859 |
| siteHAMBURG | 4 | 2 | 1 | 0.534 | LdlPFC | RdlPFC | 1.6690285 |
| siteLONDON | 1 | 6 | 0.9 | 0.808 | LNacc | RNacc | 2.2820168 |
| siteLONDON | 1 | 6 | 0.9 | 0.808 | LpACC | RpACC | 1.2521861 |
| siteLONDON | 1 | 6 | 0.9 | 0.808 | LsgACC | RsgACC | 0.3765052 |
| siteLONDON | 1 | 6 | 0.9 | 0.808 | LNacc | LdlPFC | 0.0291021 |
| siteLONDON | 1 | 6 | 0.9 | 0.808 | RpACC | RdlPFC | 0.5076589 |
| siteLONDON | 1 | 6 | 0.9 | 0.808 | LdlPFC | RdlPFC | 0.1905903 |
| siteLONDON | 1 | 6 | 0.9 | 0.808 | LpACC | mPFC | 0.4532805 |
| siteNOTTINGHAM | 1 | 1 | 1 | 1 | LNacc | RNacc | 0.316937 |
| siteNOTTINGHAM | 1 | 1 | 1 | 1 | LpACC | RpACC | 0.1916174 |
| siteNOTTINGHAM | 1 | 1 | 1 | 1 | LpACC | LsgACC | 0.3640014 |
| siteNOTTINGHAM | 1 | 1 | 1 | 1 | RpACC | LsgACC | 0.7600617 |
| siteNOTTINGHAM | 1 | 1 | 1 | 1 | LpACC | RsgACC | 0.6369236 |
| siteNOTTINGHAM | 1 | 1 | 1 | 1 | RpACC | RsgACC | 0.3805526 |
| siteNOTTINGHAM | 1 | 1 | 1 | 1 | LpACC | mPFC | 0.1822788 |
| sitePARIS | 4 | 2 | 0.998 | 0.346 | LdlPFC | RdlPFC | 0.0336 |
| sitePARIS | 4 | 2 | 0.998 | 0.346 | RpACC | vmPFC | 0.6745204 |
| sitePARIS | 4 | 2 | 0.998 | 0.346 | mPFC | vmPFC | 0.1258775 |

***Table S10:*** *Longitudinal Male model (CDT: p<0.005)*

| predictor | component | n_edges | p_ncomp | p_strn | ROI1 | ROI2 | t_value |
| --- | --- | --- | --- | --- | --- | --- | --- |
| timepointFU3 | 1 | 5 | 0.0064 | 0.0004 | LNacc | RNacc | -1.6400562 |
| timepointFU3 | 1 | 5 | 0.0064 | 0.0004 | RNacc | RpACC | -0.3729687 |
| timepointFU3 | 1 | 5 | 0.0064 | 0.0004 | LNacc | RsgACC | -1.5701425 |
| timepointFU3 | 1 | 5 | 0.0064 | 0.0004 | RNacc | RsgACC | -1.691749 |
| timepointFU3 | 1 | 5 | 0.0064 | 0.0004 | LsgACC | RsgACC | -3.0736749 |
| mean_fd | 1 | 39 | 0.0114 | 0.0012 | LNacc | RNacc | 5.775339 |
| mean_fd | 1 | 39 | 0.0114 | 0.0012 | LNacc | LpACC | 3.1374995 |
| mean_fd | 1 | 39 | 0.0114 | 0.0012 | RNacc | LpACC | 2.4008524 |
| mean_fd | 1 | 39 | 0.0114 | 0.0012 | LNacc | RpACC | 3.7851431 |
| mean_fd | 1 | 39 | 0.0114 | 0.0012 | RNacc | RpACC | 3.4416154 |
| mean_fd | 1 | 39 | 0.0114 | 0.0012 | LpACC | RpACC | 3.6295764 |
| mean_fd | 1 | 39 | 0.0114 | 0.0012 | LNacc | LsgACC | 3.8984595 |
| mean_fd | 1 | 39 | 0.0114 | 0.0012 | RNacc | LsgACC | 3.9005701 |
| mean_fd | 1 | 39 | 0.0114 | 0.0012 | LpACC | LsgACC | 0.3238557 |
| mean_fd | 1 | 39 | 0.0114 | 0.0012 | RpACC | LsgACC | 1.9279403 |
| mean_fd | 1 | 39 | 0.0114 | 0.0012 | LNacc | RsgACC | 4.4384875 |
| mean_fd | 1 | 39 | 0.0114 | 0.0012 | RNacc | RsgACC | 4.1648936 |
| mean_fd | 1 | 39 | 0.0114 | 0.0012 | LpACC | RsgACC | 0.819486 |
| mean_fd | 1 | 39 | 0.0114 | 0.0012 | RpACC | RsgACC | 1.5115836 |
| mean_fd | 1 | 39 | 0.0114 | 0.0012 | LsgACC | RsgACC | 2.2648785 |
| mean_fd | 1 | 39 | 0.0114 | 0.0012 | RNacc | LdlPFC | 0.5773882 |
| mean_fd | 1 | 39 | 0.0114 | 0.0012 | LpACC | LdlPFC | 0.7323083 |
| mean_fd | 1 | 39 | 0.0114 | 0.0012 | RpACC | LdlPFC | 0.167995 |
| mean_fd | 1 | 39 | 0.0114 | 0.0012 | LsgACC | LdlPFC | 0.8872511 |
| mean_fd | 1 | 39 | 0.0114 | 0.0012 | RsgACC | LdlPFC | 0.3704242 |
| mean_fd | 1 | 39 | 0.0114 | 0.0012 | LNacc | RdlPFC | 0.4957702 |
| mean_fd | 1 | 39 | 0.0114 | 0.0012 | LpACC | RdlPFC | 1.6490475 |
| mean_fd | 1 | 39 | 0.0114 | 0.0012 | RpACC | RdlPFC | 0.4290873 |
| mean_fd | 1 | 39 | 0.0114 | 0.0012 | LsgACC | RdlPFC | 0.365819 |
| mean_fd | 1 | 39 | 0.0114 | 0.0012 | RsgACC | RdlPFC | 0.2242117 |
| mean_fd | 1 | 39 | 0.0114 | 0.0012 | LNacc | mPFC | 1.9534294 |
| mean_fd | 1 | 39 | 0.0114 | 0.0012 | RNacc | mPFC | 1.1810599 |
| mean_fd | 1 | 39 | 0.0114 | 0.0012 | RpACC | mPFC | 0.2222279 |
| mean_fd | 1 | 39 | 0.0114 | 0.0012 | LsgACC | mPFC | 0.3323914 |
| mean_fd | 1 | 39 | 0.0114 | 0.0012 | LdlPFC | mPFC | 0.8139885 |
| mean_fd | 1 | 39 | 0.0114 | 0.0012 | RdlPFC | mPFC | 1.6530927 |
| mean_fd | 1 | 39 | 0.0114 | 0.0012 | LNacc | vmPFC | 4.397337 |
| mean_fd | 1 | 39 | 0.0114 | 0.0012 | RNacc | vmPFC | 4.1787986 |
| mean_fd | 1 | 39 | 0.0114 | 0.0012 | LpACC | vmPFC | 0.285378 |
| mean_fd | 1 | 39 | 0.0114 | 0.0012 | RpACC | vmPFC | 1.2395842 |
| mean_fd | 1 | 39 | 0.0114 | 0.0012 | LsgACC | vmPFC | 2.5937265 |
| mean_fd | 1 | 39 | 0.0114 | 0.0012 | RsgACC | vmPFC | 1.826765 |
| mean_fd | 1 | 39 | 0.0114 | 0.0012 | LdlPFC | vmPFC | 0.7778513 |
| mean_fd | 1 | 39 | 0.0114 | 0.0012 | RdlPFC | vmPFC | 0.7919335 |
| siteDUBLIN | 1 | 1 | 1 | 1 | LNacc | RNacc | 0.071469 |
| siteDUBLIN | 1 | 1 | 1 | 1 | LpACC | RpACC | 0.8482357 |
| siteDUBLIN | 1 | 1 | 1 | 1 | LpACC | mPFC | 1.4455046 |
| siteDUBLIN | 1 | 1 | 1 | 1 | RpACC | mPFC | 0.3159564 |
| siteDUBLIN | 1 | 1 | 1 | 1 | LpACC | vmPFC | 0.2399208 |
| siteDUBLIN | 1 | 1 | 1 | 1 | RpACC | vmPFC | 0.8511696 |
| siteDUBLIN | 1 | 1 | 1 | 1 | mPFC | vmPFC | 0.1260028 |
| siteHAMBURG | 7 | 1 | 1 | 0.8526 | LdlPFC | RdlPFC | 1.4333364 |
| siteLONDON | 1 | 1 | 1 | 0.9898 | LNacc | RNacc | 2.0463247 |
| siteLONDON | 1 | 1 | 1 | 0.9898 | LpACC | RpACC | 1.016494 |
| siteLONDON | 1 | 1 | 1 | 0.9898 | LsgACC | RsgACC | 0.1408131 |
| siteLONDON | 1 | 1 | 1 | 0.9898 | RpACC | RdlPFC | 0.2719668 |
| siteLONDON | 1 | 1 | 1 | 0.9898 | LpACC | mPFC | 0.2175884 |
| siteNOTTINGHAM | 1 | 1 | 1 | 1 | LNacc | RNacc | 0.0812449 |
| siteNOTTINGHAM | 1 | 1 | 1 | 1 | LpACC | LsgACC | 0.1283093 |
| siteNOTTINGHAM | 1 | 1 | 1 | 1 | RpACC | LsgACC | 0.5243697 |
| siteNOTTINGHAM | 1 | 1 | 1 | 1 | LpACC | RsgACC | 0.4012315 |
| siteNOTTINGHAM | 1 | 1 | 1 | 1 | RpACC | RsgACC | 0.1448605 |
| sitePARIS | 4 | 1 | 1 | 0.1556 | RpACC | vmPFC | 0.4388284 |

***Table S11:*** *Longitudinal Female model (CDT: p<0.01)*

| Variable | Cluster | No. Edges | p-FWE (Size) | p-FWE (Strength) | ROI1 | ROI2 | Strength |
| --- | --- | --- | --- | --- | --- | --- | --- |
| timepointFU3 | 3 | 1 | 0.2388 | 0.1082 | LpACC | RpACC | -0.3847387 |
| timepointFU3 | 3 | 1 | 0.2388 | 0.1082 | LsgACC | RsgACC | -2.9664587 |
| threat_z | 1 | 6 | 0.136 | 0.0978 | LNacc | LsgACC | 1.4318628 |
| threat_z | 1 | 6 | 0.136 | 0.0978 | RNacc | LsgACC | 0.5623726 |
| threat_z | 1 | 6 | 0.136 | 0.0978 | LNacc | RsgACC | 0.1760215 |
| threat_z | 1 | 6 | 0.136 | 0.0978 | LsgACC | mPFC | 0.2503055 |
| threat_z | 1 | 6 | 0.136 | 0.0978 | LNacc | vmPFC | 0.4543078 |
| threat_z | 1 | 6 | 0.136 | 0.0978 | RNacc | vmPFC | 0.6135654 |
| deprivation_z | 1 | 10 | 0.004 | 0.0046 | LNacc | RpACC | -0.2495723 |
| deprivation_z | 1 | 10 | 0.004 | 0.0046 | LNacc | LsgACC | -0.436863 |
| deprivation_z | 1 | 10 | 0.004 | 0.0046 | RNacc | LsgACC | -0.1256913 |
| deprivation_z | 1 | 10 | 0.004 | 0.0046 | LpACC | LsgACC | -1.0658042 |
| deprivation_z | 1 | 10 | 0.004 | 0.0046 | RpACC | LsgACC | -0.5732687 |
| deprivation_z | 1 | 10 | 0.004 | 0.0046 | LpACC | RsgACC | -0.0568471 |
| deprivation_z | 1 | 10 | 0.004 | 0.0046 | RpACC | RsgACC | -0.042466 |
| deprivation_z | 1 | 10 | 0.004 | 0.0046 | RNacc | mPFC | -0.042585 |
| deprivation_z | 1 | 10 | 0.004 | 0.0046 | LsgACC | mPFC | -1.1354232 |
| deprivation_z | 1 | 10 | 0.004 | 0.0046 | RsgACC | mPFC | -0.3909858 |
| mean_fd | 1 | 40 | 0.0396 | 0.0006 | LNacc | RNacc | 6.4865465 |
| mean_fd | 1 | 40 | 0.0396 | 0.0006 | LNacc | LpACC | 3.3644465 |
| mean_fd | 1 | 40 | 0.0396 | 0.0006 | RNacc | LpACC | 1.3816025 |
| mean_fd | 1 | 40 | 0.0396 | 0.0006 | LNacc | RpACC | 2.3303192 |
| mean_fd | 1 | 40 | 0.0396 | 0.0006 | RNacc | RpACC | 0.6574189 |
| mean_fd | 1 | 40 | 0.0396 | 0.0006 | LpACC | RpACC | 3.6030553 |
| mean_fd | 1 | 40 | 0.0396 | 0.0006 | LNacc | LsgACC | 5.5617793 |
| mean_fd | 1 | 40 | 0.0396 | 0.0006 | RNacc | LsgACC | 5.0488126 |
| mean_fd | 1 | 40 | 0.0396 | 0.0006 | LpACC | LsgACC | 1.3289375 |
| mean_fd | 1 | 40 | 0.0396 | 0.0006 | RpACC | LsgACC | 0.1688795 |
| mean_fd | 1 | 40 | 0.0396 | 0.0006 | LNacc | RsgACC | 3.9641569 |
| mean_fd | 1 | 40 | 0.0396 | 0.0006 | RNacc | RsgACC | 4.1249317 |
| mean_fd | 1 | 40 | 0.0396 | 0.0006 | LpACC | RsgACC | 0.3165213 |
| mean_fd | 1 | 40 | 0.0396 | 0.0006 | LsgACC | RsgACC | 4.685575 |
| mean_fd | 1 | 40 | 0.0396 | 0.0006 | LNacc | LdlPFC | 1.6340952 |
| mean_fd | 1 | 40 | 0.0396 | 0.0006 | RNacc | LdlPFC | 0.925673 |
| mean_fd | 1 | 40 | 0.0396 | 0.0006 | LpACC | LdlPFC | 2.9256972 |
| mean_fd | 1 | 40 | 0.0396 | 0.0006 | RpACC | LdlPFC | 2.2684829 |
| mean_fd | 1 | 40 | 0.0396 | 0.0006 | LsgACC | LdlPFC | 2.0080182 |
| mean_fd | 1 | 40 | 0.0396 | 0.0006 | RsgACC | LdlPFC | 1.2559459 |
| mean_fd | 1 | 40 | 0.0396 | 0.0006 | LNacc | RdlPFC | 1.5957399 |
| mean_fd | 1 | 40 | 0.0396 | 0.0006 | RNacc | RdlPFC | 2.2593693 |
| mean_fd | 1 | 40 | 0.0396 | 0.0006 | LpACC | RdlPFC | 3.5297 |
| mean_fd | 1 | 40 | 0.0396 | 0.0006 | RpACC | RdlPFC | 1.136965 |
| mean_fd | 1 | 40 | 0.0396 | 0.0006 | LsgACC | RdlPFC | 2.7668339 |
| mean_fd | 1 | 40 | 0.0396 | 0.0006 | RsgACC | RdlPFC | 1.4622151 |
| mean_fd | 1 | 40 | 0.0396 | 0.0006 | LNacc | mPFC | 3.7374603 |
| mean_fd | 1 | 40 | 0.0396 | 0.0006 | RNacc | mPFC | 2.1971903 |
| mean_fd | 1 | 40 | 0.0396 | 0.0006 | LpACC | mPFC | 2.7357258 |
| mean_fd | 1 | 40 | 0.0396 | 0.0006 | RpACC | mPFC | 2.166686 |
| mean_fd | 1 | 40 | 0.0396 | 0.0006 | LsgACC | mPFC | 0.5435735 |
| mean_fd | 1 | 40 | 0.0396 | 0.0006 | LdlPFC | mPFC | 2.7785295 |
| mean_fd | 1 | 40 | 0.0396 | 0.0006 | RdlPFC | mPFC | 3.0908914 |
| mean_fd | 1 | 40 | 0.0396 | 0.0006 | LNacc | vmPFC | 3.0659225 |
| mean_fd | 1 | 40 | 0.0396 | 0.0006 | RNacc | vmPFC | 3.7370763 |
| mean_fd | 1 | 40 | 0.0396 | 0.0006 | LpACC | vmPFC | 0.7319362 |
| mean_fd | 1 | 40 | 0.0396 | 0.0006 | LsgACC | vmPFC | 4.0584599 |
| mean_fd | 1 | 40 | 0.0396 | 0.0006 | RsgACC | vmPFC | 2.4524666 |
| mean_fd | 1 | 40 | 0.0396 | 0.0006 | LdlPFC | vmPFC | 1.138629 |
| mean_fd | 1 | 40 | 0.0396 | 0.0006 | RdlPFC | vmPFC | 0.879592 |
| siteDUBLIN | 1 | 10 | 0.99 | 0.7812 | LNacc | RNacc | 0.9424516 |
| siteDUBLIN | 1 | 10 | 0.99 | 0.7812 | LNacc | RpACC | 0.0573209 |
| siteDUBLIN | 1 | 10 | 0.99 | 0.7812 | LpACC | RpACC | 1.8112314 |
| siteDUBLIN | 1 | 10 | 0.99 | 0.7812 | LpACC | LsgACC | 0.817749 |
| siteDUBLIN | 1 | 10 | 0.99 | 0.7812 | RpACC | LsgACC | 1.5549362 |
| siteDUBLIN | 1 | 10 | 0.99 | 0.7812 | LpACC | RsgACC | 0.1369735 |
| siteDUBLIN | 1 | 10 | 0.99 | 0.7812 | RpACC | RsgACC | 1.2896577 |
| siteDUBLIN | 1 | 10 | 0.99 | 0.7812 | LsgACC | RsgACC | 1.1965877 |
| siteDUBLIN | 1 | 10 | 0.99 | 0.7812 | LdlPFC | RdlPFC | 1.4469795 |
| siteDUBLIN | 1 | 10 | 0.99 | 0.7812 | LpACC | mPFC | 1.2685463 |
| siteDUBLIN | 1 | 10 | 0.99 | 0.7812 | RpACC | mPFC | 0.2212853 |
| siteHAMBURG | 3 | 6 | 0.9974 | 0.0552 | LpACC | RsgACC | -0.0430881 |
| siteHAMBURG | 3 | 6 | 0.9974 | 0.0552 | LpACC | LdlPFC | 0.7135913 |
| siteHAMBURG | 3 | 6 | 0.9974 | 0.0552 | RpACC | LdlPFC | 1.2457281 |
| siteHAMBURG | 3 | 6 | 0.9974 | 0.0552 | LdlPFC | RdlPFC | 0.4570953 |
| siteHAMBURG | 3 | 6 | 0.9974 | 0.0552 | LdlPFC | mPFC | 1.6489561 |
| siteHAMBURG | 3 | 6 | 0.9974 | 0.0552 | LdlPFC | vmPFC | 0.4094802 |
| siteLONDON | 1 | 10 | 1 | 0.9836 | LNacc | RNacc | 4.3640692 |
| siteLONDON | 1 | 10 | 1 | 0.9836 | LNacc | RpACC | 0.3525196 |
| siteLONDON | 1 | 10 | 1 | 0.9836 | RNacc | RpACC | 0.1022143 |
| siteLONDON | 1 | 10 | 1 | 0.9836 | LpACC | RpACC | 1.9092055 |
| siteLONDON | 1 | 10 | 1 | 0.9836 | LsgACC | RsgACC | 3.2105243 |
| siteLONDON | 1 | 10 | 1 | 0.9836 | RpACC | LdlPFC | 0.5263962 |
| siteLONDON | 1 | 10 | 1 | 0.9836 | LsgACC | LdlPFC | 0.438343 |
| siteLONDON | 1 | 10 | 1 | 0.9836 | RsgACC | LdlPFC | 0.9288747 |
| siteLONDON | 1 | 10 | 1 | 0.9836 | LdlPFC | RdlPFC | 1.3309606 |
| siteLONDON | 1 | 10 | 1 | 0.9836 | LpACC | mPFC | 1.9335722 |
| siteMANNHEIM | 3 | 8 | 0.994 | 0.9974 | LpACC | LdlPFC | 1.2913925 |
| siteMANNHEIM | 3 | 8 | 0.994 | 0.9974 | RpACC | LdlPFC | 1.7660309 |
| siteMANNHEIM | 3 | 8 | 0.994 | 0.9974 | LsgACC | LdlPFC | 0.1295965 |
| siteMANNHEIM | 3 | 8 | 0.994 | 0.9974 | RsgACC | LdlPFC | 0.4053106 |
| siteMANNHEIM | 3 | 8 | 0.994 | 0.9974 | LpACC | RdlPFC | 0.5669747 |
| siteMANNHEIM | 3 | 8 | 0.994 | 0.9974 | RpACC | RdlPFC | 0.8539461 |
| siteMANNHEIM | 3 | 8 | 0.994 | 0.9974 | LdlPFC | mPFC | 1.3675629 |
| siteMANNHEIM | 3 | 8 | 0.994 | 0.9974 | LdlPFC | vmPFC | 0.5668845 |
| siteNOTTINGHAM | 1 | 1 | 1 | 1 | LNacc | RNacc | 0.6968045 |
| siteNOTTINGHAM | 1 | 1 | 1 | 1 | LsgACC | RsgACC | 2.7845864 |
| siteNOTTINGHAM | 1 | 1 | 1 | 1 | LpACC | mPFC | 0.1656363 |
| sitePARIS | 1 | 1 | 1 | 1 | LNacc | RNacc | -0.8753342 |
| sitePARIS | 1 | 1 | 1 | 1 | LpACC | RsgACC | -0.2289136 |
| sitePARIS | 1 | 1 | 1 | 1 | RpACC | RsgACC | -0.2118124 |
| sitePARIS | 1 | 1 | 1 | 1 | LpACC | LdlPFC | 0.5309627 |
| sitePARIS | 1 | 1 | 1 | 1 | RpACC | LdlPFC | 0.2863475 |
| sitePARIS | 1 | 1 | 1 | 1 | LdlPFC | RdlPFC | 0.0284714 |
| sitePARIS | 1 | 1 | 1 | 1 | LsgACC | mPFC | -0.0934044 |
| timepointFU3:threat_z | 1 | 7 | 0.0162 | 0.0362 | RNacc | LpACC | -0.111303 |
| timepointFU3:threat_z | 1 | 7 | 0.0162 | 0.0362 | LNacc | RpACC | -0.1091864 |
| timepointFU3:threat_z | 1 | 7 | 0.0162 | 0.0362 | RNacc | RpACC | -0.4247848 |
| timepointFU3:threat_z | 1 | 7 | 0.0162 | 0.0362 | LNacc | LsgACC | -0.6103386 |
| timepointFU3:threat_z | 1 | 7 | 0.0162 | 0.0362 | RNacc | LsgACC | -0.2653239 |
| timepointFU3:threat_z | 1 | 7 | 0.0162 | 0.0362 | LNacc | RdlPFC | -0.1211784 |
| timepointFU3:threat_z | 1 | 7 | 0.0162 | 0.0362 | RNacc | vmPFC | -0.3407069 |
| timepointFU3:deprivation_z | 1 | 1 | 0.2768 | 0.2554 | LpACC | LsgACC | 0.1492508 |
| timepointFU3:deprivation_z | 1 | 1 | 0.2768 | 0.2554 | LNacc | RdlPFC | 0.0503102 |
| timepointFU3:deprivation_z | 1 | 1 | 0.2768 | 0.2554 | LsgACC | mPFC | 0.3933732 |
| timepointFU3:deprivation_z | 1 | 1 | 0.2768 | 0.2554 | RsgACC | mPFC | 0.4152775 |

***Table S12:*** *Longitudinal Female model (CDT: p<0.005)*

| Variable | Cluster | No. Edges | p-FWE (Size) | p-FWE (Strength) | ROI1 | ROI2 | Strength |
| --- | --- | --- | --- | --- | --- | --- | --- |
| timepointFU3 | 3 | 1 | 0.16 | 0.12 | LpACC | RpACC | -0.1496326 |
| timepointFU3 | 3 | 1 | 0.16 | 0.12 | LsgACC | RsgACC | -2.7313526 |
| threat_z | 1 | 4 | 0.17 | 0.13 | LNacc | LsgACC | 1.1967567 |
| threat_z | 1 | 4 | 0.17 | 0.13 | RNacc | LsgACC | 0.3272665 |
| threat_z | 1 | 4 | 0.17 | 0.13 | LNacc | vmPFC | 0.2192017 |
| threat_z | 1 | 4 | 0.17 | 0.13 | RNacc | vmPFC | 0.3784593 |
| deprivation_z | 1 | 5 | 0.03 | 0 | LNacc | LsgACC | -0.2017569 |
| deprivation_z | 1 | 5 | 0.03 | 0 | LpACC | LsgACC | -0.8306981 |
| deprivation_z | 1 | 5 | 0.03 | 0 | RpACC | LsgACC | -0.3381626 |
| deprivation_z | 1 | 5 | 0.03 | 0 | LsgACC | mPFC | -0.9003171 |
| deprivation_z | 1 | 5 | 0.03 | 0 | RsgACC | mPFC | -0.1558797 |
| mean_fd | 1 | 39 | 0.04 | 0.01 | LNacc | RNacc | 6.2514404 |
| mean_fd | 1 | 39 | 0.04 | 0.01 | LNacc | LpACC | 3.1293404 |
| mean_fd | 1 | 39 | 0.04 | 0.01 | RNacc | LpACC | 1.1464964 |
| mean_fd | 1 | 39 | 0.04 | 0.01 | LNacc | RpACC | 2.0952131 |
| mean_fd | 1 | 39 | 0.04 | 0.01 | RNacc | RpACC | 0.4223128 |
| mean_fd | 1 | 39 | 0.04 | 0.01 | LpACC | RpACC | 3.3679492 |
| mean_fd | 1 | 39 | 0.04 | 0.01 | LNacc | LsgACC | 5.3266732 |
| mean_fd | 1 | 39 | 0.04 | 0.01 | RNacc | LsgACC | 4.8137065 |
| mean_fd | 1 | 39 | 0.04 | 0.01 | LpACC | LsgACC | 1.0938314 |
| mean_fd | 1 | 39 | 0.04 | 0.01 | LNacc | RsgACC | 3.7290508 |
| mean_fd | 1 | 39 | 0.04 | 0.01 | RNacc | RsgACC | 3.8898256 |
| mean_fd | 1 | 39 | 0.04 | 0.01 | LpACC | RsgACC | 0.0814152 |
| mean_fd | 1 | 39 | 0.04 | 0.01 | LsgACC | RsgACC | 4.4504689 |
| mean_fd | 1 | 39 | 0.04 | 0.01 | LNacc | LdlPFC | 1.3989891 |
| mean_fd | 1 | 39 | 0.04 | 0.01 | RNacc | LdlPFC | 0.6905669 |
| mean_fd | 1 | 39 | 0.04 | 0.01 | LpACC | LdlPFC | 2.6905911 |
| mean_fd | 1 | 39 | 0.04 | 0.01 | RpACC | LdlPFC | 2.0333768 |
| mean_fd | 1 | 39 | 0.04 | 0.01 | LsgACC | LdlPFC | 1.7729121 |
| mean_fd | 1 | 39 | 0.04 | 0.01 | RsgACC | LdlPFC | 1.0208398 |
| mean_fd | 1 | 39 | 0.04 | 0.01 | LNacc | RdlPFC | 1.3606338 |
| mean_fd | 1 | 39 | 0.04 | 0.01 | RNacc | RdlPFC | 2.0242632 |
| mean_fd | 1 | 39 | 0.04 | 0.01 | LpACC | RdlPFC | 3.2945939 |
| mean_fd | 1 | 39 | 0.04 | 0.01 | RpACC | RdlPFC | 0.9018589 |
| mean_fd | 1 | 39 | 0.04 | 0.01 | LsgACC | RdlPFC | 2.5317278 |
| mean_fd | 1 | 39 | 0.04 | 0.01 | RsgACC | RdlPFC | 1.227109 |
| mean_fd | 1 | 39 | 0.04 | 0.01 | LNacc | mPFC | 3.5023542 |
| mean_fd | 1 | 39 | 0.04 | 0.01 | RNacc | mPFC | 1.9620842 |
| mean_fd | 1 | 39 | 0.04 | 0.01 | LpACC | mPFC | 2.5006197 |
| mean_fd | 1 | 39 | 0.04 | 0.01 | RpACC | mPFC | 1.9315799 |
| mean_fd | 1 | 39 | 0.04 | 0.01 | LsgACC | mPFC | 0.3084674 |
| mean_fd | 1 | 39 | 0.04 | 0.01 | LdlPFC | mPFC | 2.5434234 |
| mean_fd | 1 | 39 | 0.04 | 0.01 | RdlPFC | mPFC | 2.8557853 |
| mean_fd | 1 | 39 | 0.04 | 0.01 | LNacc | vmPFC | 2.8308164 |
| mean_fd | 1 | 39 | 0.04 | 0.01 | RNacc | vmPFC | 3.5019702 |
| mean_fd | 1 | 39 | 0.04 | 0.01 | LpACC | vmPFC | 0.4968301 |
| mean_fd | 1 | 39 | 0.04 | 0.01 | LsgACC | vmPFC | 3.8233538 |
| mean_fd | 1 | 39 | 0.04 | 0.01 | RsgACC | vmPFC | 2.2173605 |
| mean_fd | 1 | 39 | 0.04 | 0.01 | LdlPFC | vmPFC | 0.9035229 |
| mean_fd | 1 | 39 | 0.04 | 0.01 | RdlPFC | vmPFC | 0.6444859 |
| siteDUBLIN | 1 | 1 | 1 | 1 | LNacc | RNacc | 0.7073455 |
| siteDUBLIN | 1 | 1 | 1 | 1 | LpACC | RpACC | 1.5761253 |
| siteDUBLIN | 1 | 1 | 1 | 1 | LpACC | LsgACC | 0.5826429 |
| siteDUBLIN | 1 | 1 | 1 | 1 | RpACC | LsgACC | 1.3198301 |
| siteDUBLIN | 1 | 1 | 1 | 1 | RpACC | RsgACC | 1.0545516 |
| siteDUBLIN | 1 | 1 | 1 | 1 | LsgACC | RsgACC | 0.9614816 |
| siteDUBLIN | 1 | 1 | 1 | 1 | LdlPFC | RdlPFC | 1.2118734 |
| siteDUBLIN | 1 | 1 | 1 | 1 | LpACC | mPFC | 1.0334402 |
| siteHAMBURG | 3 | 5 | 0.88 | 0.04 | LpACC | LdlPFC | 0.4784852 |
| siteHAMBURG | 3 | 5 | 0.88 | 0.04 | RpACC | LdlPFC | 1.010622 |
| siteHAMBURG | 3 | 5 | 0.88 | 0.04 | LdlPFC | RdlPFC | 0.2219892 |
| siteHAMBURG | 3 | 5 | 0.88 | 0.04 | LdlPFC | mPFC | 1.41385 |
| siteHAMBURG | 3 | 5 | 0.88 | 0.04 | LdlPFC | vmPFC | 0.1743741 |
| siteLONDON | 1 | 9 | 0.99 | 0.97 | LNacc | RNacc | 4.1289631 |
| siteLONDON | 1 | 9 | 0.99 | 0.97 | LNacc | RpACC | 0.1174135 |
| siteLONDON | 1 | 9 | 0.99 | 0.97 | LpACC | RpACC | 1.6740994 |
| siteLONDON | 1 | 9 | 0.99 | 0.97 | LsgACC | RsgACC | 2.9754182 |
| siteLONDON | 1 | 9 | 0.99 | 0.97 | RpACC | LdlPFC | 0.2912901 |
| siteLONDON | 1 | 9 | 0.99 | 0.97 | LsgACC | LdlPFC | 0.2032369 |
| siteLONDON | 1 | 9 | 0.99 | 0.97 | RsgACC | LdlPFC | 0.6937686 |
| siteLONDON | 1 | 9 | 0.99 | 0.97 | LdlPFC | RdlPFC | 1.0958545 |
| siteLONDON | 1 | 9 | 0.99 | 0.97 | LpACC | mPFC | 1.6984661 |
| siteMANNHEIM | 3 | 7 | 1 | 0.99 | LpACC | LdlPFC | 1.0562864 |
| siteMANNHEIM | 3 | 7 | 1 | 0.99 | RpACC | LdlPFC | 1.5309248 |
| siteMANNHEIM | 3 | 7 | 1 | 0.99 | RsgACC | LdlPFC | 0.1702045 |
| siteMANNHEIM | 3 | 7 | 1 | 0.99 | LpACC | RdlPFC | 0.3318686 |
| siteMANNHEIM | 3 | 7 | 1 | 0.99 | RpACC | RdlPFC | 0.61884 |
| siteMANNHEIM | 3 | 7 | 1 | 0.99 | LdlPFC | mPFC | 1.1324568 |
| siteMANNHEIM | 3 | 7 | 1 | 0.99 | LdlPFC | vmPFC | 0.3317784 |
| siteNOTTINGHAM | 1 | 1 | 1 | 1 | LNacc | RNacc | 0.4616984 |
| siteNOTTINGHAM | 1 | 1 | 1 | 1 | LsgACC | RsgACC | 2.5494803 |
| sitePARIS | 1 | 1 | 1 | 0 | LNacc | RNacc | -0.6402281 |
| sitePARIS | 1 | 1 | 1 | 0 | LpACC | LdlPFC | 0.2958566 |
| sitePARIS | 1 | 1 | 1 | 0 | RpACC | LdlPFC | 0.0512414 |
| timepointFU3:threat_z | 1 | 4 | 0.03 | 0.07 | RNacc | RpACC | -0.1896787 |
| timepointFU3:threat_z | 1 | 4 | 0.03 | 0.07 | LNacc | LsgACC | -0.3752325 |
| timepointFU3:threat_z | 1 | 4 | 0.03 | 0.07 | RNacc | LsgACC | -0.0302178 |
| timepointFU3:threat_z | 1 | 4 | 0.03 | 0.07 | RNacc | vmPFC | -0.1056008 |
| timepointFU3:deprivation_z | 5 | 2 | 0.06 | 0.09 | LsgACC | mPFC | 0.1582671 |
| timepointFU3:deprivation_z | 5 | 2 | 0.06 | 0.09 | RsgACC | mPFC | 0.1801714 |

**Sensitivity analysis: Sex-specific attrition in moderate to severe adversity participants**

**Table S13:** Logistic regression output from model predicting follow-up status (1 = available rsfMRI data at follow-up two and follow-up three, 0 = available rsfMRI data at follow-up two but missing at follow-up three).

| **Variable** | **Odds Ratio** | **SE** | **Statistic** | **p value** | **95%CI low** | **95%CI high** |
| --- | --- | --- | --- | --- | --- | --- |
| (Intercept) | 42.797753 | 2.445878 | 1.5358434 | 0.1245768 | 0.3570939 | 5324.3788 |
| SexMale | 1.0348405 | 0.1751593 | 0.1955211 | 0.844985 | 0.734235 | 1.4595894 |
| Moderate-Severe group | 1.6442223 | 0.3947895 | 1.2595763 | 0.2078222 | 0.7729934 | 3.6846934 |
| age_fu2 | 0.8510814 | 0.1306724 | -1.2339828 | 0.2172093 | 0.6579546 | 1.0994751 |
| siteDRESDEN | 0.4193679 | 0.3519837 | -2.4688829 | 0.0135536 | 0.2070333 | 0.8268687 |
| siteDUBLIN | 0.4225277 | 0.425635 | -2.0240352 | 0.0429665 | 0.1810472 | 0.9663462 |
| siteHAMBURG | 0.5659858 | 0.400132 | -1.4224961 | 0.1548823 | 0.2557132 | 1.2343114 |
| siteLONDON | 0.5160128 | 0.3447587 | -1.9190923 | 0.0549727 | 0.2584413 | 1.0036117 |
| siteMANNHEIM | 0.7073043 | 0.3788097 | -0.9141642 | 0.3606306 | 0.3330561 | 1.4784496 |
| siteNOTTINGHAM | 0.6469396 | 0.360855 | -1.2068625 | 0.2274851 | 0.3147919 | 1.3020883 |
| sitePARIS | 0.3178212 | 0.395082 | -2.9013374 | 0.0037157 | 0.1441762 | 0.6816606 |
| SexMale: Moderate-Severe group | 0.3466126 | 0.5874789 | -1.8035501 | 0.0713019 | 0.1065568 | 1.0786528 |

**Sex-specific associations between adversity and anhedonia at IMAGEN follow-up three**

**Table S14:** *Model output for linear regression examining associations of adversity, including sex*adversity interaction terms and controlling for site and age.*

| **Variable** | **Estimate** | **SE** | **Statistic** | **p value** | **95%CI low** | **95%CI high** |
| --- | --- | --- | --- | --- | --- | --- |
| (Intercept) | 0.5811991 | 0.8406727 | -0.6455092 | 0.518907 | 0.1114106 | 3.0319584 |
| threat_z | 1.0652983 | 0.0393625 | 1.6069822 | 0.1087186 | 0.9860087 | 1.150964 |
| SexMale | 0.9522229 | 0.0579433 | -0.8448984 | 0.3985903 | 0.8497517 | 1.0670511 |
| deprivation_z | 1.0543711 | 0.0496813 | 1.0656816 | 0.2871056 | 0.956307 | 1.1624911 |
| age_fu2 | 1.0456813 | 0.0451962 | 0.9883256 | 0.3234932 | 0.9568207 | 1.1427943 |
| siteDRESDEN | 0.7687192 | 0.1158746 | -2.2699496 | 0.0236551 | 0.6121879 | 0.9652743 |
| siteDUBLIN | 0.7756696 | 0.1388815 | -1.829103 | 0.068007 | 0.5904192 | 1.0190442 |
| siteHAMBURG | 0.741682 | 0.1324058 | -2.2569603 | 0.0244613 | 0.5717782 | 0.9620727 |
| siteLONDON | 0.706177 | 0.1114955 | -3.1202093 | 0.0019165 | 0.5672409 | 0.8791433 |
| siteMANNHEIM | 0.9428747 | 0.1212056 | -0.4853067 | 0.6276811 | 0.7430563 | 1.1964271 |
| siteNOTTINGHAM | 0.9842915 | 0.1171555 | -0.1351472 | 0.8925524 | 0.7818936 | 1.2390813 |
| sitePARIS | 0.8264263 | 0.1302295 | -1.4639114 | 0.1438755 | 0.6398397 | 1.0674244 |
| threat_z:SexMale | 0.9401135 | 0.0746063 | -0.8277411 | 0.4082302 | 0.8119216 | 1.0885452 |
| SexMale:deprivation_z | 1.1661232 | 0.0689612 | 2.2285691 | 0.0263069 | 1.0183465 | 1.3353444 |

**References**

1.	Bartra, O., McGuire, J. T. & Kable, J. W. The valuation system: A coordinate-based meta-analysis of BOLD fMRI experiments examining neural correlates of subjective value. *NeuroImage* **76**, 412–427 (2013).

2.	Kelly, A. M. C. *et al.* Development of Anterior Cingulate Functional Connectivity from Late Childhood to Early Adulthood. *Cereb. Cortex* **19**, 640–657 (2009).

3.	Dhami, P. *et al.* Prefrontal Cortical Reactivity and Connectivity Markers Distinguish Youth Depression from Healthy Youth. *Cereb. Cortex N. Y. NY* **30**, 3884–3894 (2020).

4.	Rupprechter, S. *et al.* Blunted medial prefrontal cortico-limbic reward-related effective connectivity and depression. *Brain J. Neurol.* **143**, 1946–1956 (2020).
